# Interpretable deep learning for deconvolutional analysis of neural signals

**DOI:** 10.1101/2024.01.05.574379

**Authors:** Bahareh Tolooshams, Sara Matias, Hao Wu, Simona Temereanca, Naoshige Uchida, Venkatesh N. Murthy, Paul Masset, Demba Ba

**Author notes:** BT and SM have contributed equally to this work. DB and PM have jointly supervised this work. **Data availability:** Data and code will be deposited in a public repository upon publication. **Competing interests:** The authors declare no competing interests.

## Abstract

The widespread adoption of deep learning to build models that capture the dynamics of neural populations is typically based on “black-box” approaches that lack an interpretable link between neural activity and network parameters. Here, we propose to apply algorithm unrolling, a method for interpretable deep learning, to design the architecture of sparse deconvolutional neural networks and obtain a direct interpretation of network weights in relation to stimulus-driven single-neuron activity through a generative model. We characterize our method, referred to as deconvolutional unrolled neural learning (DUNL), and show its versatility by applying it to deconvolve single-trial local signals across multiple brain areas and recording modalities. To exemplify use cases of our decomposition method, we uncover multiplexed salience and reward prediction error signals from midbrain dopamine neurons in an unbiased manner, perform simultaneous event detection and characterization in somatosensory thalamus recordings, and characterize the heterogeneity of neural responses in the piriform cortex and in the striatum during unstructured, naturalistic experiments. Our work leverages the advances in interpretable deep learning to gain a mechanistic understanding of neural activity.

## Introduction

Understanding the activity of neurons, both at the single neuron and population levels, in relation to features in the environment and the behaviour of an organism, is a key question in neuroscience. Recent technological advancements and experimental methods have allowed researchers to record from an increasingly large population of identified single neurons using high-throughput electrophysiology or imaging in animals performing complex tasks [1–3]. In such complex environments, external events might unfold on various timescales, giving rise to neural signals expressed over different timescales across the population of recorded neurons. Moreover, these neural representations show complex dynamics and differing levels of multiplexing. For example, single neurons across the cortical hierarchy exhibit varying degrees of mixed selectivity to task parameters depending on task structure and demands [4–9]. Neuromodulatory neurons, such as midbrain dopamine neurons, can respond to different environmental and internal variables [10, 11]. Additionally, single-neuron activity has been proposed to be composed of multiple components required for reward evaluation, such as valence and salience [12, 13].

To understand how multiple representations in single neurons enable population-level computations, we need scalable and reliable methods for decomposing their activity into overlapping and non-overlapping local components/events that can capture important intrinsic heterogeneity in the recorded populations. Here, we develop such a deconvolutional method within a deep learning framework.

A reasonable deconvolution method ought to meet several requirements. First, the method should be able to be implemented on single instantiations of the neural data without the need for averaging over trials or animals [14–16]. Preferably, it should apply to both structured and naturalistic tasks in which there is little or no trial structure [17–20]. Second, it should be flexible concerning the source signal (e.g., spike count data or a proxy signal such as calcium levels via a fluorescent indicator [21]). Third, the method should utilize an expressive class of mappings from latent representations to data, namely ones that can capture the complexity of neural data. Fourth and importantly, the method should be interpretable. In this manuscript, interpretability is addressed in two contexts: first, we aim for interpretability within the neural network, where its architecture, weights, and representations can be directly linked to the optimization, parameters, and representations of a generative model. The second aspect focuses on the model’s ability to provide a direct mapping between stimuli (internal or external) and latent variables. For example, a probabilistic generative model, adaptable to different neural data types, can capture the locality of neural responses and the temporal sparsity of stimuli, interpreted via neural impulse responses [22–24]. This level of interpretability depends on domain knowledge, such as task structure or hypotheses about the data. Finally, the method should be scalable and practical for analyzing large-scale neuroscience datasets.

Most classical neural data analysis methods are primarily designed for dimensionality reduction, aiming to find a low-rank structure governing the neural data globally, and thus are not suited for deconvolutional analysis. Examples include PCA [25, 26], NMF [27], GPFA [28], vLGP [29], mTDR [30], TCA [31, 32], and SCA [33]. In contrast, deconvolutional methods recover locally lowrank structures in the data. Methods such as Poisson GLM [34], convNMF[35, 36], seqNMF[37], and point process models for neural sequences (PP-Seq) [15] can fit neural data with locally low-rank structures and offer interpretability via direct mappings between data and latent representations. However, none fully meet all requirements, such as scalability or respecting the statistics of the source signal. For example, Poisson GLM uses predefined components, and seqNMF does not account for source signal statistics.

The limitations of classical methods, along with the scalability of deep learning, have inspired the community to explore deep learning approaches for neural data analysis. Indeed, prior work using deep learning has addressed, to varying extents, all the discussed desiderata, except the fourth, by extracting a low-dimensional latent space from the neural data through a non-linear deep neural architecture [38–42]. However, they do not provide a direct link between the contributions of single neurons or neuron types and the population level computation, owing to their “black-box” approach typical of deep networks [43, 44]. Our method complements these existing tools by extracting interpretable impulse-like responses of multiplexed signals from single neurons, recovering locally low-rank structures, which can be further used to characterize heterogeneity and homogeneity across neural populations.

In the deep learning literature, interpretability methods can be categorized into two groups [45]: explainable and interpretable. The former, also called *mechanistically interpretable* deep learning, develops interpretability methods to explain black-box models. For example, in computer vision, saliency maps are constructed to highlight input image pixels that are discriminative with respect to an output decision of a deep neural network [46, 47]. A more generalizable example is Local Interpretable Model-Agnostic Explanations (LIME), a framework for explaining predictions of any black-box model by learning a locally-interpretable model around the prediction of interest [48]. However, this class of models does not make the neural network interpretable in and of itself: the model tries to explain what the network does. First, this means that there is no direct mapping from the embedding to the data: for instance, the explainable model might conclude that the network is optimizing for a feature that is correlated with the learning objective of the network, missing the true understanding of the “black-box” system [49]. Second, this approach does not guide the neural network architecture to learn interpretable representations. That is, the network may perform discrimination based on non-generalizable spurious features [50]. The majority of deep neural networks [51–53], including many current methods used in neuroscience [38, 42], falls into this category. While recent work gains mechanistic insights into neural circuit computations [54, 55], such analysis was achieved from *a posteriori* interpretation and analysis of the network architecture.

In contrast, model-based interpretable deep learning [56] (Figure 1A) is an emerging technique to design deep neural networks that are inherently interpretable. In particular, algorithm unrolling [57], a sub-category of interpretable deep learning, offers deep neural networks whose weights and representations can be directly interpreted as parameters and variables of an underlying generative model [57, 58]. This one-to-one mapping between the neural weights and latent representations of a generative model introduces interpretability. These mappings can be learned using an iterative algorithm optimizing the model [58–60]. Importantly, this generative model does not require detailed assumptions about the data: it provides domain knowledge information without restricting the model’s output such that important data features are missed. Following seminal work in algorithm unrolling [58], numerous applications have been developed across several fields, including computational imaging (e.g., super-resolution [61] and image deblurring [62]), medical imaging [63, 64], identification of dynamical systems [65], remote sensing applications (e.g., radar imaging [66]) or source separation in speech processing [67].

**Figure 1.**
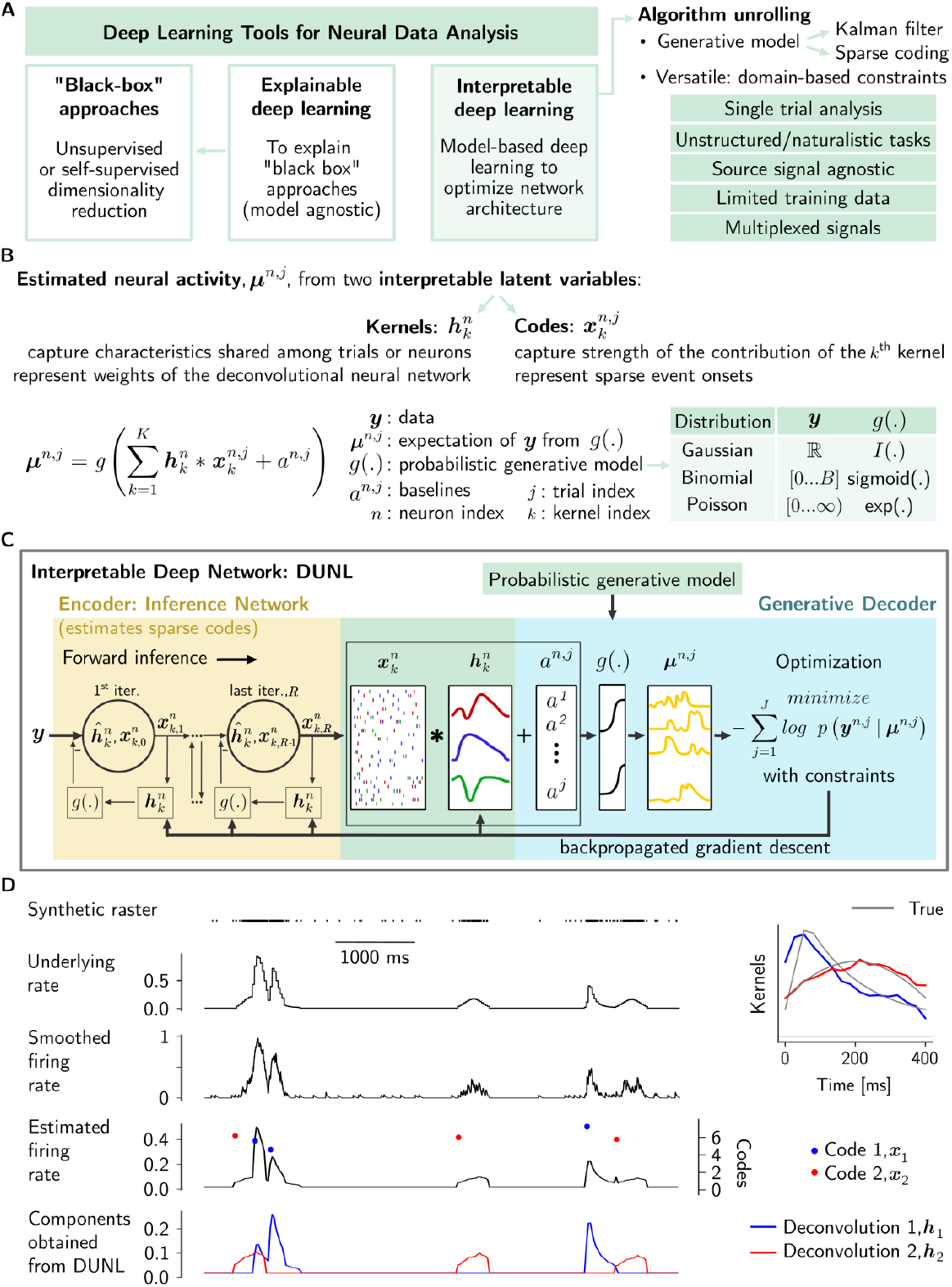
Interpretable deep learning with deconvolutional unrolled neural learning: DUNL. (A) Categorization of deep learning tools developed for neural data analysis and advantages of using algorithm unrolling. (B) Generative model used by DUNL to estimate neural activity as a function of the summation of convolutions between kernels and sparse codes. (C) Schematic representation of DUNL: the deep inference network (yellow), whose weights are the estimated kernels, estimates the sparse codes. Both the estimated codes and kernels serve as input to the generative decoder (blue; the overlapping area between encoder and decoder, with shared variables, is represented in green). The output of this decoder is used to optimize the network by feeding back the estimated kernels. More details are provided in the Supplementary Materials - Methods and Figure S1 (D) Demonstration of DUNL’s ability to deconvolve events from a synthetic neuron in an unstructured single-trial, where two recurrent events occur locally at random times and with varying amplitudes (left). DUNL recovers an estimate (colored) of the true kernels (gray) after training with 25 trials (right).

Here, we propose a novel framework combining algorithm unrolling with convolutional sparse coding (i.e., dictionary learning), called Deconvolutional Unrolled Neural Learning (DUNL), that ful-fills all the above-listed desiderata (Figure 1A). Our method offers a flexible framework to deconvolve single-trial neuronal activity into interpretable and local components; this interpretability is made possible by the choice of convolutional sparse coding as the generative model. Our source code is flexible, easy to use, and adaptable to various applications by simple modifications of neural network non-linearities and training loss functions without requiring the user to re-derive an optimization algorithm for their specific application.

To demonstrate the versatility and usefulness of DUNL, we apply it to the deconvolution of neural signals acquired in a wide range of experimental conditions. First, we show that it can deconvolve the salience and reward prediction error (RPE) components of naturally multiplexed reward signals encoded by dopamine neurons in the midbrain. Second, we demonstrate that it can deconvolve cue and outcome components of slow calcium signals recorded from dopamine neurons during associative learning. Third, we show simultaneous event detection and characterization of neural activity from the thalamus in a high signal-to-noise ratio (SNR) setting. Fourth, we demonstrate that in a low SNR setting, we can extract classes of neural responses from the piriform cortex in the presence of random and overlapping odor pulses. Fifth, we characterize axonal dopaminergic activity in the striatum during a single session of a naturalistic experiment. Finally, we perform a thorough characterization of our method and situate it within the plethora of decomposition methods available to neuroscientists, to show that local interpretability in a limited data regime and scalability are important features of our deconvolution method.

## Results

### Sparse deconvolutional learning uncovers structure in single-trial, single-neuron activity

We aim to decompose single-trial neural activity into local impulse-like responses to sparse yet recurring events; One may interpret the decomposition as finding locally low-rank structures in neural data. We assume that the observed neural activity is the result of a combination of recurring components – kernels of a “dictionary” – whose timing and magnitudes can vary on an event-by-event basis. Thus, we seek to reconstruct the neural data by optimizing a model that learns these components or kernels. To achieve this, the neural activity is modeled as the linear combination of convolutions of these kernels and a set of vectors representing the timing and the strength of neural response to recurring sparse events. We refer to each of the vectors as a sparse code. Stochasticity in the neural activity is accounted for by passing this convolved signal through a link function, used to model the mean activity under a probability distribution of the natural exponential family (e.g., Gaussian, Binomial, and Poisson, see Table S1) to match the signal statistics.

More specifically, we model (Figure 1B) the observations ***y***^*n,j*^ from neuron *n* at trial *j* using the natural exponential family [68, 69] (e.g., Binomial or Poisson for spiking and Gaussian for calcium signals) with distribution mean of ***μ***^*n,j*^. We impose a generative model on the *n*^th^ neuron’s mean activity at trial *j*, ***μ***^*n,j*^, and express it as the convolution of *K* localized kernels 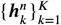 and sparse codes (representations) 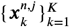,along with a background, baseline, measured activity level *a*^*n,j*^ (Figure 1B). The convolutional aspect enables the identification of local patterns occurring across time, and the discovery of *locally* low-rank structures. While this manuscript focuses on the locality of the kernels, DUNL can also be used to capture long-range temporal features. Kernel learning via DUNL goes beyond the *pre-defined* family of basis functions used in previous methods like Poisson GLM regression for neural data [34]. Moreover, kernels and codes are interpretable in the following sense. Kernels capture characteristics shared among trials : they characterize a neuron’s response to time-sensitive sparse and asynchronous inferred events. When the support of the inferred sparse codes accurately represents the stimuli, this characterization is considered the neuron’s response to the stimuli. Alternatively, DUNL’s flexible design allows for sharing kernels across neurons to obtain population-level impulse responses to synchronous events, where the neuron index is dropped: 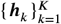 while code amplitudes remain neuron-, trial-, and event-specific. In this case, the optimization is done jointly for all neurons. The nonzero entries of the sparse latent representation 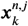 represent the time when the event associated with the kernel *k* occurs in trial *j*; their amplitude captures the strength of the neural response. In relation to the functional identification of a system, this model characterizes the system in terms of cause-effect relationship: the code aims to capture the timing of a stimulus applied locally in time [70]; the kernel captures the impulse response of the neuron [34].

Thus, the kernels in the model are non-parametric and learned fully from data, i.e., they do not obey a user-specified parametric form, and the codes are sparse in time. We learn the kernels and infer the codes by minimizing the negative data log-likelihood 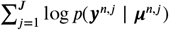 regularized/pe-nalized by terms encouraging desired properties on the codes and kernels. We impose a sparsity prior, to promote a few code activations in time, and an optional second-order covariance structure on the codes to capture dependencies among kernels (e.g., discouraging activation of two event types simultaneously). Where needed, we apply smoothing regularization on the kernels [71, 72] (Supplementary Materials - Methods). We note that the location of these codes can be fully inferred by DUNL during the optimization process, or the support (their locations) can be provided as an input, in which case only the code amplitudes need to be estimated. Whether the support was provided or inferred is specified for each example application in the respective figures below. We map the optimization into an encoder/decoder neural architecture following the algorithm unrolling approach [57] (Figure 1C, Figure S1). We call this framework Deconvolutional Unrolled Neural Learning (DUNL), an application of algorithm unrolling to convolutional dictionary learning [73–75]. The encoder is a deep-structured recurrent convolutional neural network. Unlike sequential deep encoder approaches, this encoder shares the same parameters as the generative model/decoder. The encoder takes single neuron single-trial observation ***y***^*n,j*^ as input and encodes it into a set of sparse representations 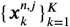 that explain the data via a set of temporally local kernels 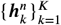. As described above, this latent code corresponds to estimated event/stimuli onsets and the strength of neural response to the event. The decoder is a shallow network based on the proposed generative model. This decoder maps the estimated time series of sparse representations into a time-series estimate of the mean neural activity. Both the encoder and decoder are characterized by the kernels 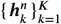 from the generative model (i.e., kernels are weights of the artificial deep neural network and can be trained by backpropagation). While this work focuses on sparsity and local low-rank structures, DUNL can, in principle, be extended to learn dense representations covering the entire trial length, alongside the sparse local representations. This approach is known as dense and sparse autoencoders [76]. Finally, training DUNL involves both a forward pass (inference for codes) and a backward pass (training to learn the kernels; see Supplementary Materials - Methods), parallelizable over neurons and trials.

To demonstrate the capabilities of DUNL, we first apply it to synthetic spiking data. The experiment consists of two event types characterized by local kernels. Within one trial, the events happen 3 times uniformly at random. In this unstructured experiment, the goal is to recover the underlying kernels and the timing and magnitude of the associated events, independently of whether these are single events or composed events (superposition of more than one kernel). DUNL successfully decomposes the synthetic neural data into kernels and codes (Figure 1D), and is able to do so with a limited amount of data (see following sections).

To summarize, we introduce a novel framework to recover the statistics of time series data as a sparse superposition of kernels (Figure 1D), that is akin to a convolutional generalization of Generalized Linear Models (GLMs), in which both coefficients and kernels are learnable, contrary to GLMs in which the kernels, as a set of basis, are user-defined [34]. Importantly, our method outputs a response amplitude for each individual occurrence of an event per trial per neuron, independently of the length of the trials. These features are absent from other encoding methods.

### DUNL uncovers salience and value signals from single dopamine neurons

We first apply DUNL to deconvolve multiplexed signals in the responses of dopamine neurons. The activity of dopamine neurons in the midbrain has long been an interest of neuroscientists, both in fundamental and clinical research, given their involvement in motivated behaviours, learning, and multiple physiological functions. A subset of these neurons located in the Ventral Tegmental Area (VTA) has been described as encoding a reward prediction error (RPE) from temporal difference (TD) reinforcement learning algorithms [77–82]. This computation requires the neural representation of the value of rewards in the environment: a transient positive RPE response signals an unexpected increase in the value of the environment. However, reward is a subjective quantity that is non-linearly modulated along multiple dimensions of reward (e.g., probability, size, etc.). It has been suggested that the reward responses of dopamine neurons multiplex two sequential and overlapping signals [83], the first one carrying information related to the salience of the reward and the second one carrying subjective value information, or utility, of the reward [12]. This distinction is important from a computational point of view because only the value-like component matches the reward prediction error signal driving learning in TD algorithms. However, in practice, most studies of dopamine neurons ignore this potential multiplexing by averaging dopamine responses over a single time window following reward delivery [84, 85], or, at best, apply user-defined ad-hoc windows to try to isolate these two contributions [86]. We used DUNL to find, in a data-driven manner, whether the reward responses of dopamine neurons can be decomposed into two components and whether these are differently modulated by reward value.

We used electrophysiological data from 40 optogenetically identified dopamine neurons [84, 85] recorded in mice performing a classical conditioning task as part of a previous study [87] (Figure 2a and see Methods). In *“Unexpected”* trials, a reward of varying size (i.e., 0.1 to 20 *μl*) was delivered without a cue, and in *“Expected”* trials, an odor cue preceded reward delivery by 1.5 s (Figure 2A,B). Although the cue predicted the timing of the reward, it provided no information about its magnitude. We modeled the data with three non-negative kernels: one to characterize the response to the odor cue, and another two for the reward event (Figure 2C,D top) [12]. The kernels are shared across the neural population, hence jointly optimized across neurons. DUNL was provided with the timing of the cue and reward events but not the trial types (reward amounts). The ground-truth event onsets are used in the encoder to identify the support of the code ***x***. Notably, the data was not aligned across trials to any events, as DUNL can handle trials of varying lengths. The goal is to recover the generating kernels, associated with the cue and reward events, given only raw spiking data and the timing of these events. Having shared kernels across the population of neurons, the codes are individualized for each neuron in single trials (Figure 2D bottom), such that each neuron is characterized by its own decomposition of its estimated firing rate. In this setting, DUNL’s encoder outputs single-trial, single-neuron specific sparse codes and background firing rate estimates.

**Figure 2.**
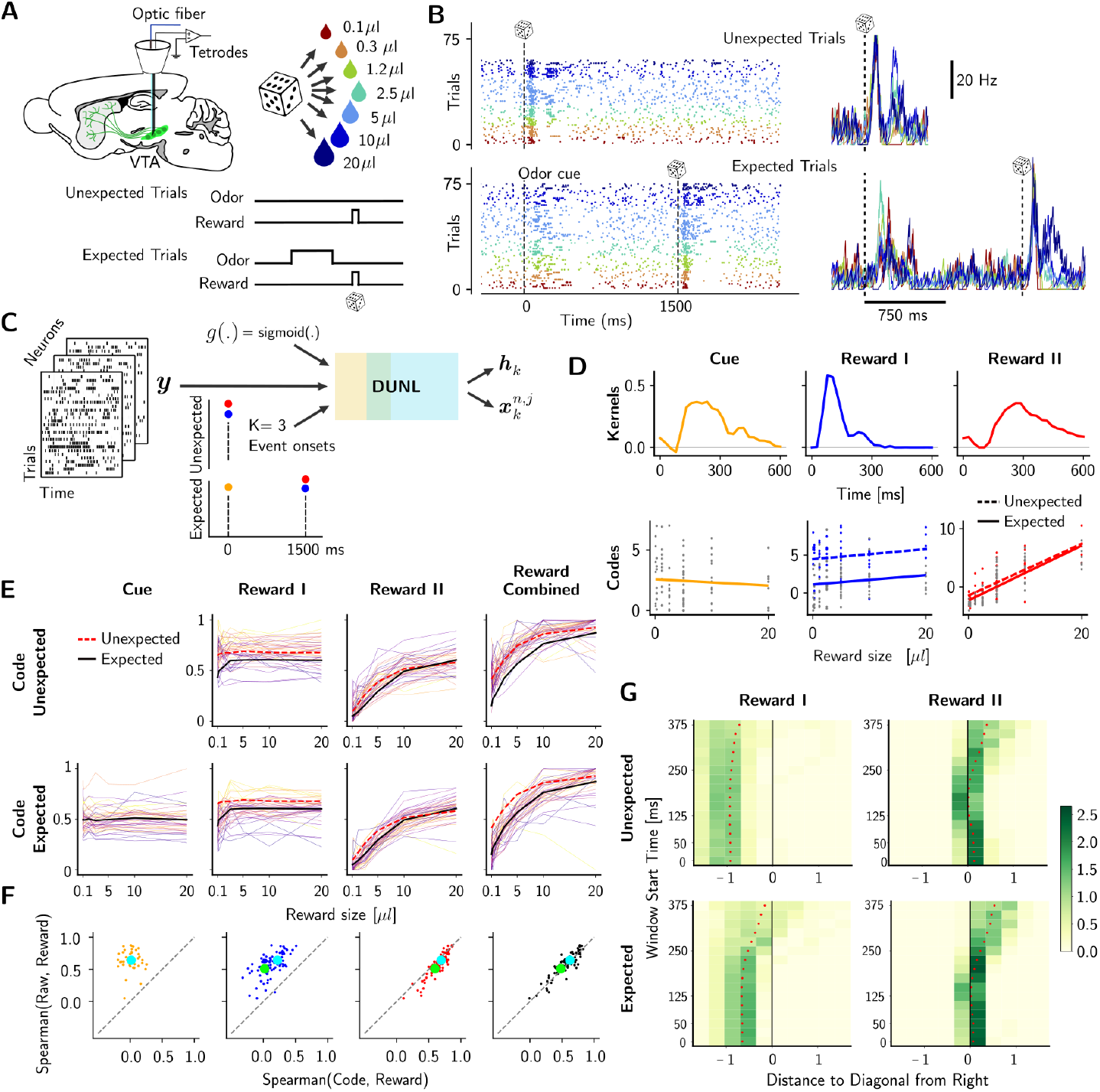
DUNL uncovers salience and value signals from single dopamine neurons’ reward responses. (A) Experimental setup showing optical fiber and tetrode recordings on a sagittal slice of the mouse brain (top left), distribution of reward sizes (top right), and task structure (bottom). (B) Raster plot (each dot is a spike) from one neuron (left) and corresponding firing rate averaged across trials of the same reward size. (C) Representation of the input information used to run DUNL in this dataset: neuron activity across time and trials, timing of stimuli, number of kernels to learn, probabilistic generative model for spiking data. (D) Learned kernels shared across neurons (top) and inferred code amplitudes for one example neuron (bottom, each dot responds to a single-trial code, the line is the linear regression of the codes over reward sizes). (E) Diversity of neural encodings (code amplitudes) as a function of reward size for expected and unexpected trials; each line represents one neuron; the black line is the average for expected trials, and the red dashed line is the average for unexpected trials. The lines are normalized per neuron, and the normalization constants are shared across trial types and codes. (F) Spearman’s rank correlation between codes and reward size vs. the windowed averaged firing rates and reward size within the full 600 ms window. The alignment of the red dots (Reward II) under the diagonal line illustrates that the value-like code is more informative about the reward size (each neuron is represented by two dots (expected and unexpected); the average of all neurons is shown by the larger circle (t-test: *p* = 0.050 expected (blue), *p* = 6.19 *∗* 10^−5^ unexpected (green)). (G) The comparison of Spearman’s correlation between the windowing and DUNL methods is mapped by measuring the distance from the diagonal (as seen in panel F) while varying the start time for the windowing method. The y-axis shows the delay from reward delivery for the start of the 600 ms window. We observe that any start time for the windowing method contains more information about value than Reward I (denser green on the left). Conversely, Reward II correlates more with value than any window from the windowing method (denser green on the right). A positive distance corresponds to values below the diagonal in panel F. The color bar represents the normalized probability density function for each bin, with the integral over each row equal to 1. For experiment results on limited data (< 8% of current analyses data) see Supplementary materials (Figure S5).

DUNL’s output showed that, as expected, the magnitude of the associated code obtained using the kernel for cue responses is essentially invariant to the reward size. More importantly, although we did not instruct DUNL to retrieve salience and value-related components separately [12], DUNL obtained two reward-related kernels which can be characterized as responding to salience (blue, Reward I) and value (red, Reward II) (Figure 2D,E,F). When we plot the code values for each kernel as a function of reward size we observe that codes corresponding to salience (blue, Reward I) are modulated by expectation (unexpected vs. expected), but almost invariant to the reward size, and codes corresponding to the value (red, Reward II) are strongly positively correlated with reward size, both for individual neurons and across the population average (Figure 2E). In fact, the value code carries more information about the reward size than the mean firing rate over a traditional ad-hoc window (Figure 2F,G). Furthermore, combining the two reward kernels (Reward I and II) (i.e., linearly adding their amplitudes) does not improve the information about the reward size, indicating that the salience-like code does not contribute to value information. We also found that as the ad-hoc window shrinks to exclude the first spike(s) traditionally attributed to salience, the ad-hoc window method improves in the representation of reward size (for Reward II, the best ad-hoc window approximately excludes the first 125 (expected) and 150 (unexpected) ms of data from the reward onset). Still, DUNL’s code is more informative of the reward value (Figure 2G).

DUNL’s successful decomposition of neural responses to the reward, as opposed to spike counts from ad-hoc windows, indicates that the code amplitudes from the value kernel in single trials are a powerful measure of the neurons’ tuning to reward size. Additionally, we ran variations of this experiment, each with different initial conditions, and obtained similar results to those in Figure 2. Specifically: 1) when kernels were learned individually for each neuron Figure S2); 2) when baseline activity was estimated from the pre-odor period instead of inferred by DUNL (Figure S3, with example neuron decomposition in Figure S4); and 3) when learning/inference was performed in a data-limited regime (< 8% of the data used in current analyses, Figure S5). To quantify the quality of our decomposition as a function of the number of trials used for training, we simulated dopamine neurons in the same experimental settings. We found that in our simulated dopamine data, we could recover well-fitted kernels with as little as 14 trials Figure S6). In summary, we showed that DUNL can discover two components in the reward responses of dopamine neurons in a systematic, data-driven approach, recovering a first salience-like component that is not modulated by reward size, while the second value-like component is. We note that although the choice of the number of kernels, in this case two for reward events, is a hyperparameter to set *a priori*, it can be tuned using validation sets (see Figure 7). Overall, DUNL will empower future studies to precisely quantify the relative contribution of salience-like and value-like components to the reward prediction error response of single dopamine neurons in an unbiased manner.

### DUNL deconvolves cue, salience, and value signals from single dopamine neurons in two-photon calcium recordings

To demonstrate DUNL’s flexibility and applicability to other data modalities beyond spike trains, we next applied DUNL to two-photon calcium imaging data [82, 88]. To this goal, we recorded the activity of 56 dopamine neurons in mice using two-photon calcium imaging with a gradient refractive index (GRIN) lens (Figure 3A) in a classical conditioning task with the same structure as in the above experiment [85, 87], but with a longer delay between the cue and the reward delivery. In unexpected trials, rewards of different sizes (i.e., 0.3 to 11 *μl*) were delivered once at a random time. In expected trials, an odor cue was delivered 3 s before the reward delivery. There is diversity in the responses of neurons to the cue and to the multiple reward sizes and, in general, we see modulation by expectation and reward size (Figure 3B).

**Figure 3.**
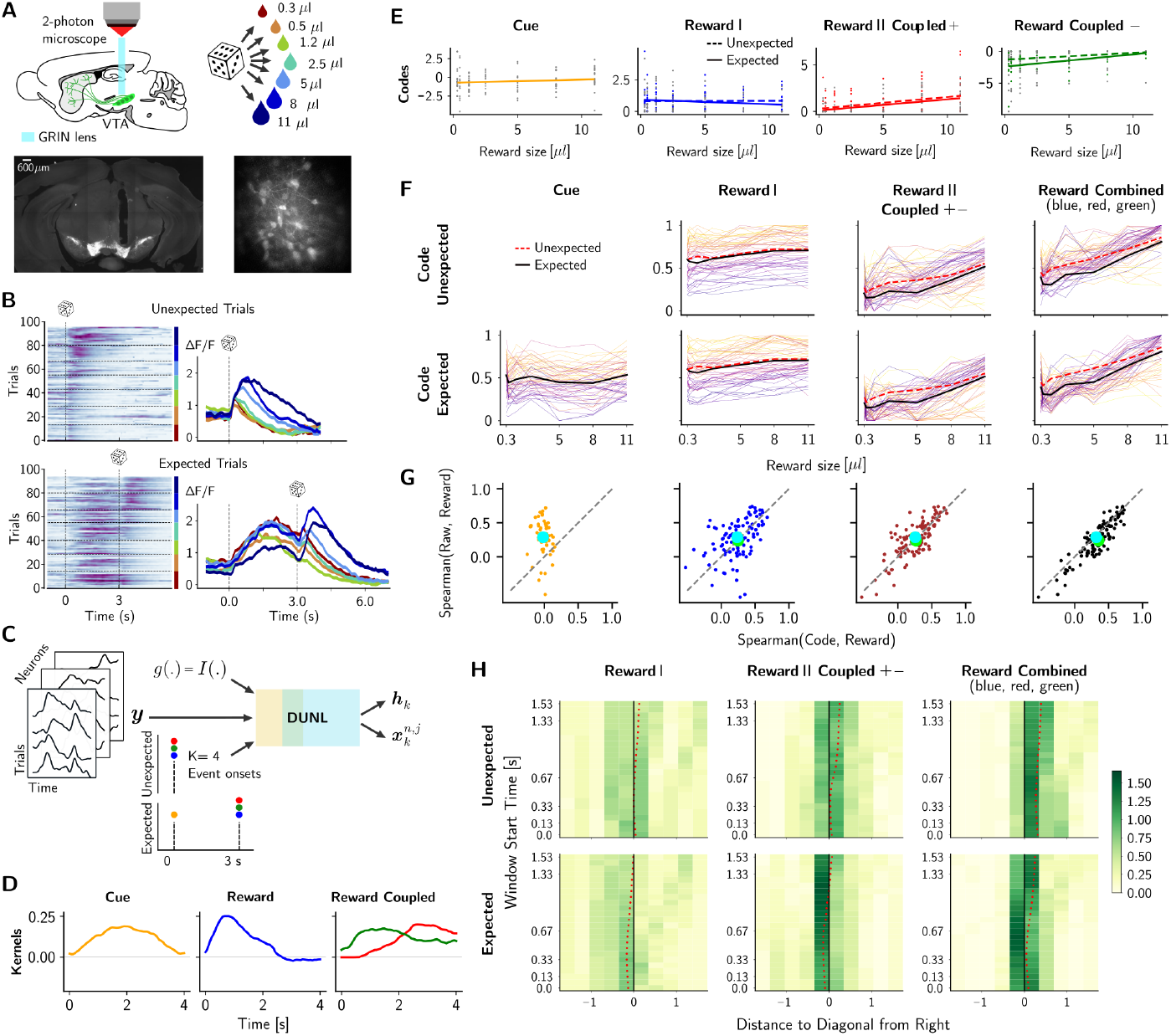
Deconvolution of the cue, salience, and value components from dopamine calcium data. (A) Top: experimental setup depicted on a sagittal slice of the mouse brain with dopamine neurons represented in green; bottom: histology images showing dopamine neurons expressing the fluorescent calcium indicator GCaMP6m: coronal slice of the mouse brain showing GRIN lens track over the VTA (left, scalebar: 600*μ*m), and projection image of the field of view obtained during an experimental acquisition under the two-photon microscope showing individual neurons (right). (B) Left: Heatmap of time-aligned trials at the reward onset. The trials are ordered from low to high reward size, with a horizontal line separating the different trial types. Right: Averaged time-aligned activity of an example neuron for each reward size. (C) Inputs used to run DUNL in this dataset: calcium activity across time and trials, timing of stimuli, number of kernels to learn, and probabilistic generative model for continuous calcium data. (D) Kernel characterization of cue and reward events. Three kernels were used to estimate the reward response: one for salience (blue) and two non-concurrent kernels for positive or negative value (green and red). (E) Code amplitude as a function of the reward size for an example neuron (each dot corresponds to the code inferred from single-trial neural activity; these values are fitted by linear regression). (F) Diversity of neural encodings as a function of reward size for unexpected (top) and expected trials (bottom): each line represents one neuron, the black line shows the average for expected trials, and the red dashed line average for unexpected trials. Activity is normalized per neuron and across trial types, and codes for comparison across subfigures. (G) Spearman correlation of the codes (x-axis) and the windowed average activity of 4 seconds (y-axis) with respect to the reward sizes: each dot represents one neuron and the average across all neurons are shown by blue (expected) and green (unexpected) large marker (Reward Combined has *p* = 0.008, and *p* = 3.468 *∗* 10^−9^ t-test, respectively). The third panel (brown) from the left combines the code from Reward Coupled kernels (positive and negative, depending on the trial). The right panel combines all the reward-related codes (salience-like Reward and value-like Reward-Coupled). (H) Heatmap of the distance of the blue (unexpected) and green (expected) circle markers in (F) to the diagonal as a measure of the increased Spearman’s correlation between codes and reward size, compared to the windowing method, as the interval chosen for the ad-hoc window is modified: it shrinks from the bottom to the top of the y-axis to gradually exclude the early activities after the onset. Positive values are located below the diagonal. On the right panel, for reward combined, the marker is closest to the diagonal when 0.4 s of activity at the reward onset is excluded in the ad-hoc window approach. Colorbar: normalized probability density function at each bin, such that the integral over each row is 1.

We characterized the neural activity using four kernels (Figure 3C), as follows. The response to the odor cue in expected trials was characterized by one kernel (blue), and the reward response (at the reward onsets) in both unexpected and expected trials was modeled by three kernels: the blue kernel can be freely active with a positive code, while the red/green kernels were positive and their codes were positive and negative, respectively. Unlike the analysis of spiking activity from dopamine neurons discussed earlier, the kernels, in this case, are 4 seconds long, causing the cue and reward kernels to overlap. As a result, window-based non-convolutional methods like PCA and NMF are not suitable. The choice of 4-second kernels is based on the raw data (Figure 3B), where neural activity in response to the cue does not return to baseline within 3 seconds, and activity in response to the reward returns to baseline after about 4 seconds. Moreover, we coupled red and green kernels such that only one of them is encouraged to be active on each trial. To achieve this, we used structured representation learning (see Methods Section in Supplementary Materials). This structural regularization is motivated to capture the different response dynamics of the calcium signal to increases versus decreases in the underlying firing of the neurons due to the different onset and offset dynamics of the sensor. DUNL’s output shows that the blue reward kernel resembles the salience response, and the red/green reward coupled kernels resemble the value response (Figure 3D). The inferred single-trial codes from a single neuron (Figure 3E) and across the population (Figure 3F) show that the salience-like kernel (Reward I) is almost invariant to reward size, while the combination of the value-components kernels (Reward II Coupled) correlates positively with reward size.

To understand this choice of kernel characterization, we looked into the interactions of the chosen kernels in the decomposition of the raw data (Figure S7). In this dataset, we observed that many neurons lack an obvious salience-like response (i.e., an early transient increase of the neural activity that is invariant to the reward size), probably because the cue-related calcium signal has not yet decayed to the baseline, potentially masking the salience response. Due to the calcium sensor’s faster onset than offset dynamics, we observed faster salience-contaminated positive responses for high-reward trials, and a very slow negative response for low-reward trials. Given the different temporal dynamics of positive and negative signals, the decomposition of reward signals into only two kernels (salience and value-like) would result in a combination of salience and valence information for both kernels, such that both kernels would be correlated with the reward size.

We computed the Spearman correlation between the Cue code, the Reward I code (blue), the Reward II coupled (red + green) code, as well as all the reward codes combined (blue + red + green) with the reward size (we combine the codes by linearly adding up their amplitude). These correlation values were then compared to the ad-hoc approach where the correlation was computed using the 4 s windowed averaged activity at the reward onset. This analysis showed that only when all reward codes (salience-like + value-like) are combined, the codes become more informative of the reward size than the ad-hoc windowing approach (the distribution of points is below the identity line, Figure 3G). This can also be noticed in the average population activity (Figure 3F). Regardless of the window size used for computing the value component of the reward response in traditional approaches, the Reward Combined code is significantly more informative of the reward size than the windowing approach in both unexpected and expected trials (Figure 3H). We attribute this success to the denoising capability of DUNL: it performs deconvolution of the cue response from the reward response, which is important in these slow calcium signals. Similar results were obtained when the baseline activity of each neuron was estimated from the pre-odor period instead of inferred by DUNL (Figure S8). Comparing these results with those obtained in electrophysiological recordings shown in Figure 2 also highlights tge flexibility of DUNL with respect to signal statistics, allows to compare neural tuning and dynamics inferred from different recording modalities [89]

### DUNL learns a population-level kernel, event onsets and infers neuron-specific event amplitudes in somatosensory thalamus recordings

Given its algorithm unrolling foundation, DUNL is a versatile framework whose inputs can be adjusted according to the application. In our previous examples, we provided DUNL with the expected number of kernels and expected times of events to guide the learning process. However, this information might be completely omitted, and we can use DUNL to perform simultaneous onset detection and learn local kernels in an unsupervised manner. This approach will be more successful in a high signal-to-noise ratio (SNR) setting.

To demonstrate this capability, we applied DUNL to electrophysiological recordings from the somatosensory thalamus of rats recorded in response to periodic whisker deflections. The whisker position was controlled by a piezoelectric stimulator using an ideal position waveform [90]. The experiment was designed with trials starting/ending with a 500 ms baseline; in the middle 2000 ms, 16 deflections of the principal whisker were applied, each with a period of 125 ms (Figure 4A). We considered the whisker position to be the stimulus and attributed a particular phase of the whisker position as the event of interest to detect. The goal is to detect the onset of the events, and characterize the neural response to the stimulus using one kernel (shared across neurons). In this experiment, onsets of events were unknown, and DUNL looks for up to 18 events in each trial (Figure 4B; the additional 2 events were added to adjust for unknown activities outside the known 16 deflection stimuli). In place of vanilla sparsity, DUNL uses group sparsity across neurons to encourage the co-incidence of event onsets across the population. We refer the reader to the Methods section and Table S4 for more information on the unsupervised DUNL method and training.

**Figure 4.**
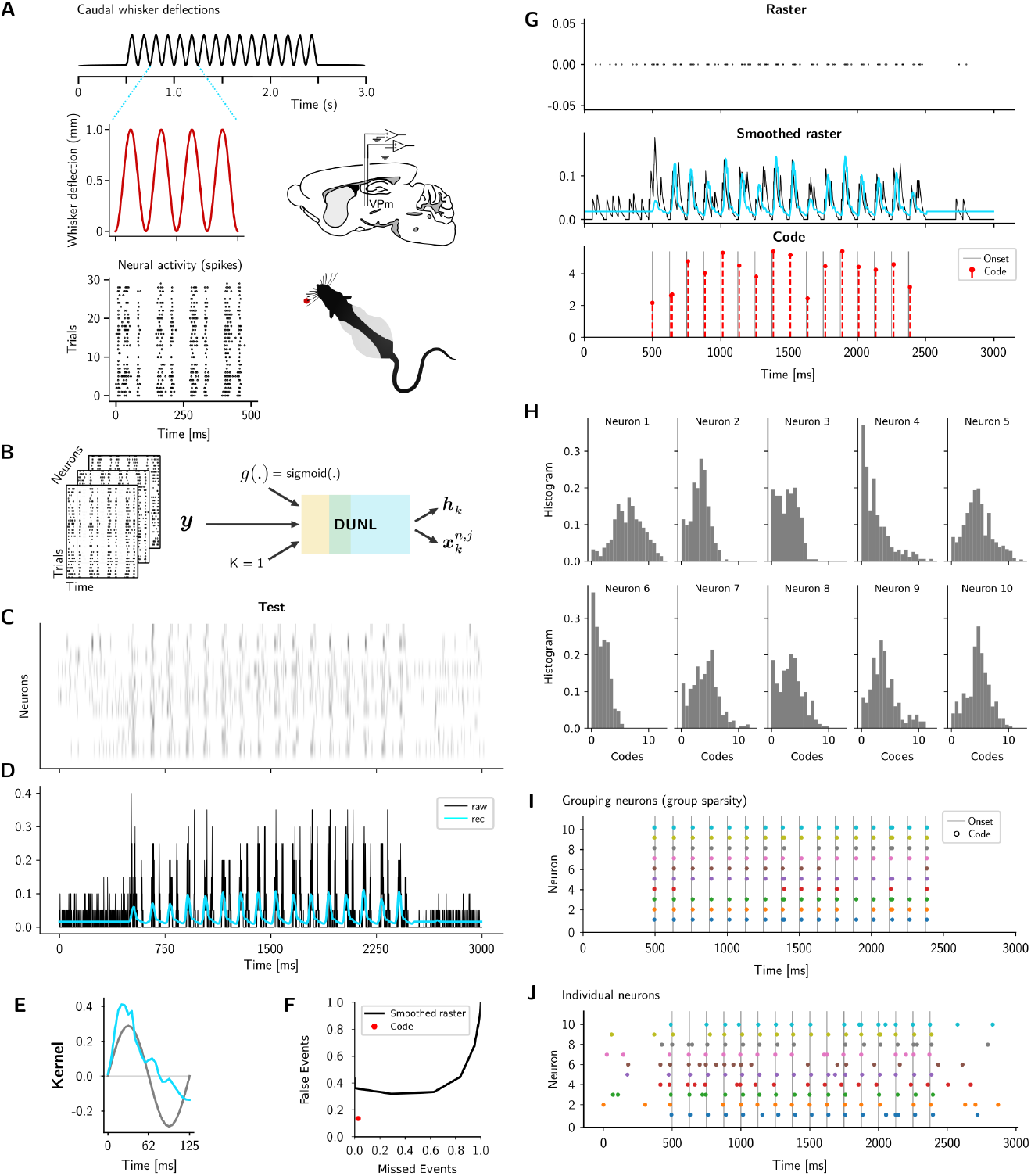
Applied to somatosensory thalamus neurons, DUNL learns a population-level kernel, event onsets and infers neuron-specific event amplitudes. (A) Experimental setup [90]: periodic whisker deflections of constant velocity are imposed in each trial (top), resulting in phase-locked neural activity (bottom left) in the ventral posteromedial (VPm) nucleus of the thalamus in anesthetized rats (bottom right). (B) DUNL setup to detect one kernel across the entire population. (C) Raster plot from 10 neurons for one trial (only test trials are shown): 16 deflections with a period of 125 ms are applied to the whisker of the rat for a total of 2000 ms. (D) Peristimulus time histogram from one example neuron (black) and DUNL estimate of the firing rates (blue). (E) Kernel characterization of the whisker motion (blue). The gray sinusoid is the first derivative of stimulus motion. (F) Quantification of the miss/false events detected by DUNL. The red dot represents the performance of DUNL when events are detected on single-trials. The black curve shows the performance of a peak-finding algorithm on the smoothed spike trains for a range of thresholds. We used a tolerance of 10*ms* (2-time bins) while computing the false/miss events. The peak finding algorithm is part of python scipy.signal package. (G) Spikes in one example trial (top), smoothed spike rate in black, and spike rate estimation in blue (middle), with the inferred code on the detection of 16+2 events in time (bottom). Each event has its own code amplitude, contrary to previously used methods. (For more information on the analyzed neurons and stimuli in relation to the original paper collecting the data [90], see Supplementary Materials). (H) Histogram of code amplitudes for individual neurons: DUNL outputs one code amplitude per event per neuron. Inferred codes can still show diverse neural responses despite the group sparsity regularization. (I) Whisker manipulation onsets detected by DUNL on a single trial and in each one of the recorded neurons when using group sparsity. (J) Whisker manipulation onsets detected by DUNL on the same trials as (H) across all neurons when not using group sparsity (i.e. inference is done separately for each neuron). Event detection is more robust when groups sparsity is used.

We divided the data in a training and test set to show that DUNL simultaneously characterizes the shape of neural spiking modulation (Figure 4C-E) and infers (detects) the event onsets at single-event, single-trial level (Figure 4F-J). The learned kernel suggests that the measured neurons encode, and are modulated by, the whisker velocity, as previously reported [91, 92].

Moreover, unlike prior work analyzing these data by averaging the time-aligned trials [68], DUNL makes inference on single-trial data (Figure 4G) with bin spike counts of only 5 ms. The heterogeneity of the inferred code amplitudes (Figure 4G,H) indicates the intrinsic variability of the neural response to the stimulus. These event, trial and neuron specific sparse codes can be used to plot a distribution of codes and perform further analysis. This feature is absent in previously published GLM analyses [91, 92], which assume the neural responses are constant across deflections. For event detection, we showed that DUNL performs significantly better than a peak-finding algorithm (Figure 4F). Here, instead of DUNL, a peak-finding algorithm can be applied to a smoothed raster and will work in this case because the neural impulse response showed a sharp increase followed by a single peak. However, unlike DUNL, peak-finding is not suitable when kernels have multiple peaks, non-sharp peaks, or when more than two kernel types are present (see Simulation Study III). We also note that event detection is more reliable when group sparsity is imposed across neurons, encouraging their activation to occur simultaneously. This group regularization aligns event onsets across neurons (Figure 4I) rather than allowing them to occur at different times for each neuron (Figure 4J). With group sparsity, a code amplitude is assigned to each event and neuron, but event detection remains robust even if individual neurons miss spikes or activity. See Figure S9 for DUNL’s results without group sparsity. With group sparsity, DUNL achieves an ***R***^2^ of 0.1612 for detecting 16 events and 0.1766 for detecting 16 + 2 events. In contrast, a GLM approach, where event onsets are known, results in ***R***^2^ = 0.1136 and 0.147 for fixed and variable code amplitudes across events, respectively (Figure S10). This experiment highlights the ability of DUNL to detect event onset while simultaneously characterizing the neural response to the event, a feature absent in prior GLM frameworks [34].

### DUNL captures intrinsic heterogeneity in neural responses during unstructured, naturalistic experiments

Finally, we highlight how DUNL can be used for exploratory data analysis. We applied DUNL to two unstructured experiments, one using known support (input the timing of events), and one with unknown support (no information about the timing of events). First, we analyzed electrophysiological data recorded from the piriform cortex of mice engaged in an olfactory task in which short (50 ms) odor pulses occur at random times (sampled from a Gamma distribution) across trials, mimicking the statistics of natural odor plumes [93]. We recorded and isolated 770 neurons from mice’s anterior piriform cortex (Figure 5A-B). The structure of piriform cortex neural responses to sequences of odor pulses are largely unexplored, and here we use DUNL to characterize them. To model neural responses, we aligned the non-zero elements of the sparse code to the timing of the odor pulses, and spike counts were modeled with a Poisson process (Figure 5C). Each neuron was characterized by one kernel. We learned both the kernels and the code amplitudes for all recorded neurons, allowing us to characterize the diversity of temporal response dynamics across piriform neurons. Using k-means clustering on the zero-mean normalized kernels, we identified 4 clusters in the population (Figure 5D); Figure 5F shows the first two principal components for visualization purposes. Three of these neural populations correspond to neurons whose activity increases following an odor pulse, albeit with different dynamics, while the other cluster corresponds to neurons whose activity is inhibited by the olfactory pulses. Similar results are obtained using a different number of clusters (Figure S11).

**Figure 5.**
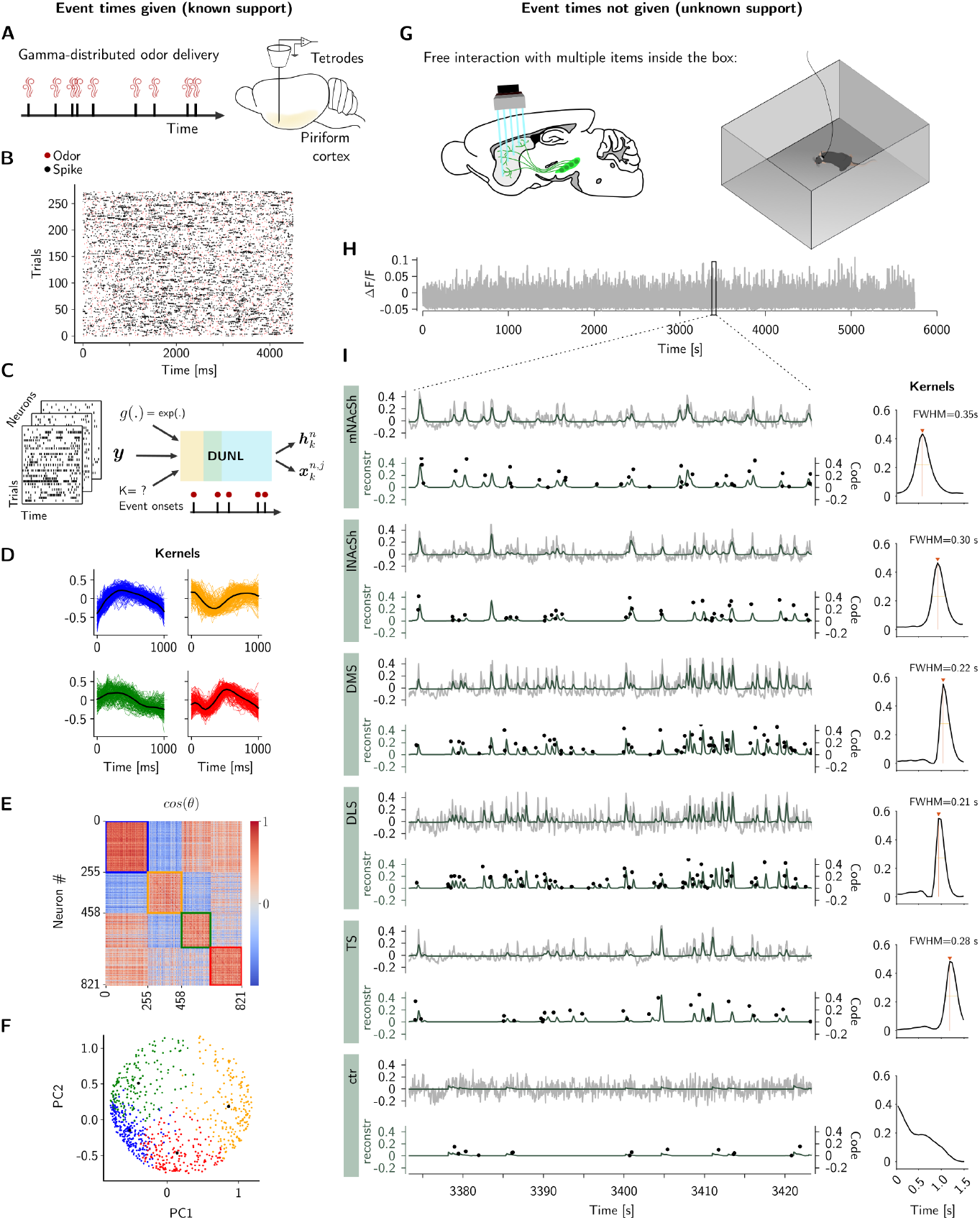
Characterizing the structure of neural activity in unstructured, naturalistic experiments. (A) Unstructured experiment with known event times: experimental task used for recording of piriform neurons: odor plumes were delivered randomly in time according to a gamma distribution. (B) Raster plot from one neuron in this unstructured trial-based experiment. The red dots denote 50 ms Gamma-distributed odor pulses, and the black dots represent single spikes. (C) Schematic of the inputs to run DUNL. (D) Kernel characterization of single neurons. The kernels (zero-mean, normalized) are shown for the four clusters. For K-means with 3 and 5 clusters, see Figure S11. (E) The cosine similarity matrix corresponding to the clusters of learned kernels. (F) Visualization of the four clusters of kernels using Principal Component Analysis (principal components 1 and 2). We note that clustering is performed on zero-mean normalized kernels, and the principal components are provided for visualization only. See Supplementary Materials for more details. (G) Unstructured experiment with unknown event times: A mouse was exposed to multiple events and objects during a 90-minute session inside a box, while calcium fluorescence signals were simultaneously recorded from dopamine axons across several striatum regions (bottom left). (H) Fluorescence trace in the nucleus accumbens lateral shell throughout the session. (I) Left: Close-up of a 50-second snippet from 5 example regions. The gray trace represents the original fluorescence input to DUNL, and the dark green trace is DUNL’s rate estimation. Codes are overlaid on the estimated neural signal (reconstruction). Right: Estimated kernels for each fiber, including full width at half maximum for each kernel. mNAcSh and lNAcSh: medial and lateral nucleus accumbens shell; DMS: dorsomedial striatum; DLS: dorsolateral striatum; TS: tail of the striatum; ctr: control fiber patch not reporting brain activity.

This application demonstrates how any type and shape of kernels can be learned by DUNL, without any assumptions guiding the shape of the kernels. Thus, DUNL can capture a diversity that may not be recoverable when using a hand-crafted family-of-basis, highlighting the value of non-parametric temporal characterization of neural responses [94].

Second, we analyzed multi-fiber photometry recordings from a single session of a behaving mouse exposed to multiple events and objects in an unstructured experiment where events were not pre-timed (Figure 5G). These recordings captured calcium concentration changes in dopamine axons across different striatal regions, each monitored by a separate optical fiber. DUNL was used to identify a single kernel for each fiber over the entire 90-minute session (Figure 5H). We observe that the width of the kernels learned by DUNL is larger in the ventral striatal regions and smaller in dorsal striatal regions, as previously reported in structured experiments [95](Figure 5I). This application demonstrates DUNL’s ability to detect events that trigger dopaminergic signaling across the striatum and to capture signal heterogeneity across regions during unstructured naturalistic behaviors. If no structure is detected in the signal for a given a fiber, DUNL outputs unstructured kernels with zero or near-zero amplitude (“ctr”: control in Figure 5I).

### Model characterization

To assess the reliability of the results reported here and guide DUNL’s end users, we characterized the performance of DUNL on a wide range of simulated data, focusing on the spiking-data modality. This section includes three distinct simulation studies mimicking scenarios commonly occurring in neuroscience experiments.

#### Simulation study I: data generation

In scenario I, we focused on a setting where a neuron responds to two different types of events, characterized by two distinct kernels of length 400 ms. In each trial, 3 different events from each kernel can occur. The timing is unstructured, such that event onsets are chosen from a uniform distribution over time, with a minimum distance of 200 ms between two events of the same type. Events of different types can occur simultaneously, thus convolving their activity (Figure 6A, blue and red events). The strength of the neural responses of the neuron was generated by the Gaussian distribution with a mean of 50 and variance of 2 for blue events and with a mean of 55 and variance of 2 for red events. The baseline firing rate was chosen to be 8 Hz.

**Figure 6.**
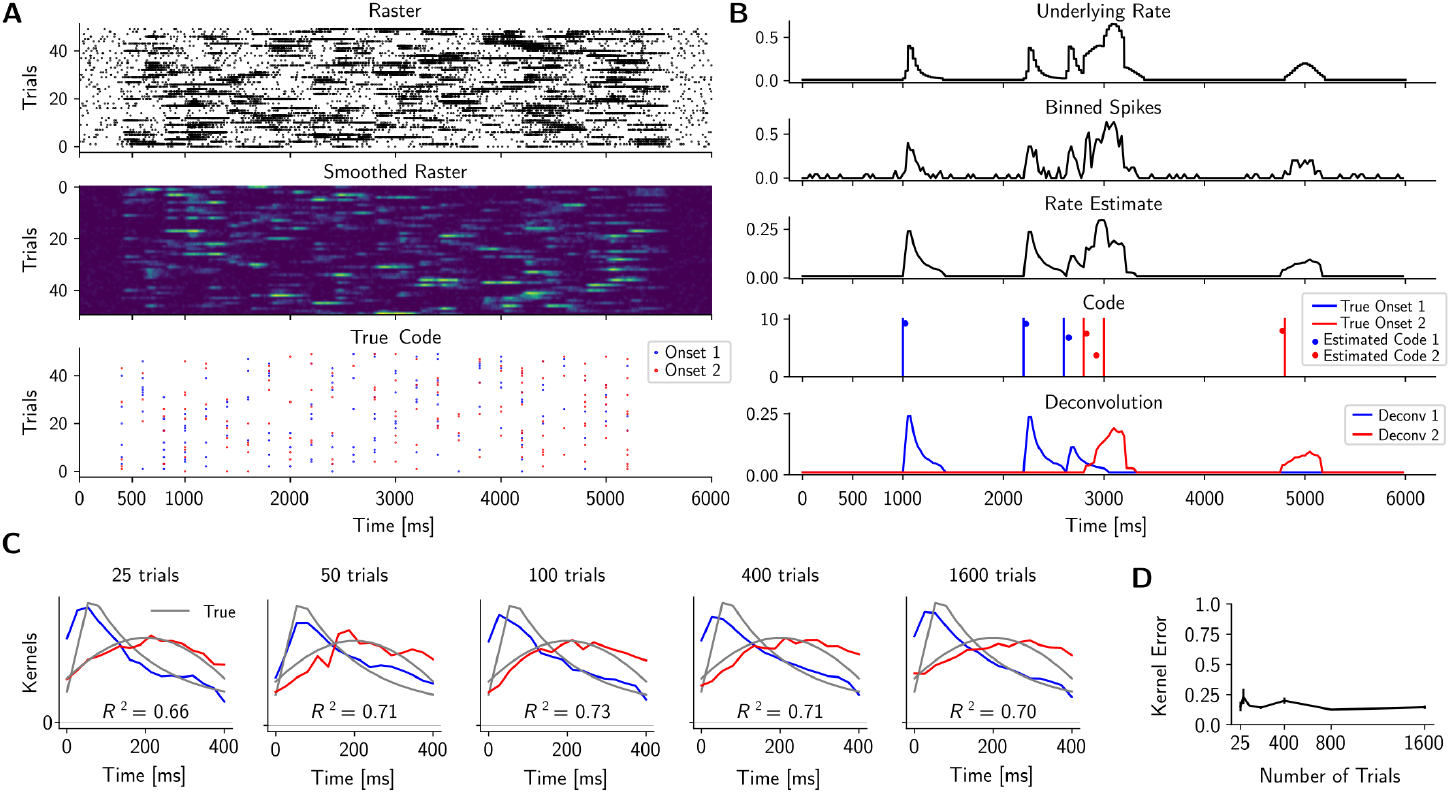
DUNL model characterization with two kernels in an unstructured experiment. (A) Test trials for this unstructured synthetic experiment: raster plot of spiking events (top), smoothed raster (middle), and true event onsets (bottom). (B) Example trial showing original firing rate (top), estimated firing rate by DUNL (middle), estimated codes and kernels (bottom). (C) Estimated kernels for 25, 50, 100, 400, and 1600 training trials. Gray traces represent the true kernels used to generate the data. The test ***R***^2^ fit score, evaluated on the binned spikes are 0.66, 0.71, 0.73, 0.74, 0.71, 0.62, 0.70 for each training size (25, 50, 100, 200, 400, 800 and 1600 trials, respectively). (D) Kernel recovery error (i.e., 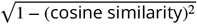,as a function of number of training trials. Specifically, we computed maximum cross-correlation to compute the cosine similarity to account for learning shifted kernels. See Figure S12 for more single trials and decomposition examples. The event detection hit rate is, on average, 80.22% and 80.68% for the smallest and largest training datasets with a tolerance of 2 time bins (with a tolerance of 3 time bins, the hit rate is increased to 91.58% and 93.34%, respectively. We report hit rate with tolerance to account for learned shifted kernels (see the Supplementary Materials - Methods for more info on the shift/delay property of DUNL).

**Figure 7.**
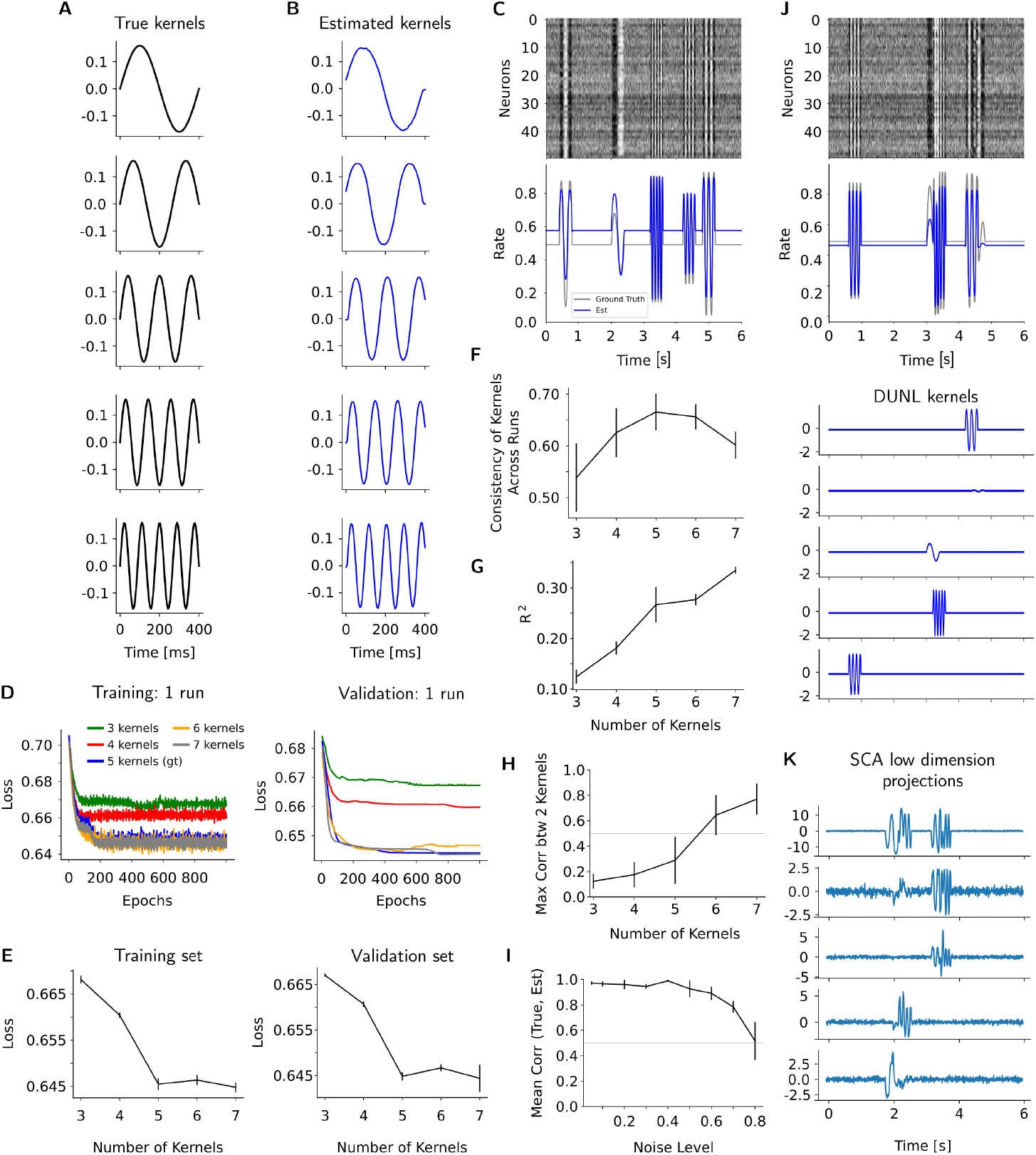
DUNL model characterization with 5 orthogonal kernels. (A) 5 orthogonal kernels used to generate synthetic data consisting on an unstructured experiment in which the 5 kernels appear once in each trial, at random time points. (B) Kernels reconstructed by DUNL using 100 trials, 20 unrollings. (C) Top: raster plot of an example trial showing 50 out of 1000 simulated neurons. The baseline firing rate and amplitude of the response varies across neurons, but they all respond to the kernels. Bottom: True underlying rate (gray) and average reconstructed rate (blue) of one neuron of the same trial shown in B. (D) Loss as a function of epoch in the training dataset (left) and in the validation dataset (right), for multiple number of kernels requested (3-7), in a single run. (E) Average loss vs. the number of kernels for the training dataset (left) and for the validation dataset (right). The reported loss is computed across 30 runs of different initialization for various number of trials (i.e., 20, 50, 100, 200, 500, and 1000. (F) Consistency of kernels across runs as a function of the number of kernels. (G) ***R***^2^ as a function of the number of kernels used to run DUNL. (H) Maximal correlation between all pairs of kernels recovered by DUNL as a function of the number of kernels used to run DUNL. The correlation approaches 1 for a number of kernels above the true number. (I) Mean correlation between the true and estimated kernels as a function of additive noise. Related to Figure S16. (J) Example trial (raster on top and underlying rate from one neuron below) with two overlapped local kernels that are recovered by DUNL (bottom). (K) 5 low-dimensional projections obtained with SCA [33] for the same trial as J. Despite the orthogonality of the kernels, SCA cannot deconvolve local kernels within these single trials.

#### Simulation study I: fitting using DUNL

We trained DUNL with these synthetic data using bin size resolution of 25 ms while the number of trials available for training varies from 25 to 1600 (results in Figure 6B and Figure S12 are from a test set with 500 trials). The number of events in each trial was known, but the timing of the events was unknown to DUNL. DUNL estimated the firing rate of the neuron and deconvolved it into two components, corresponding to each event type. Moreover, the magnitude of the sparse codes inferred by DUNL encoded the local activity of each event (kernel) within the trial (Figure 6B). Lastly, the result held with small kernel recovery error in the limited data regime, i.e., 25 training trials (Figure 6C,D, see Supplementary Materials - Methods section for optimization properties of DUNL).

#### Simulation study II: data generation

In this scenario, we restricted ourselves to the setting of a single neuron and a single event to assess how well DUNL can learn the kernel and codes associated with a neuron given its general firing properties. This model characterization allows end users to carefully assess the reliability of DUNL based on their data statistics. We evaluated the performance of DUNL as a function of the background firing rate (i.e., 2, 5, 8, 11, 14, and 17 Hz), the bin-size the model used to count spikes (i.e., 5, 10, 25, and 50 ms) (Figure S13), and the number of trials (i.e., 10, 25, 250, 500, 1000), available to learn the model parameters (i.e., kernels) (Figure S14).

We simulated multiple trials of activity, a subset of which we used for training, and the other for testing. Each trial was 4000 ms long with 1 ms resolution. In each trial, 5 similar events happen uniformly at random with a minimum distance of 200 ms. We assumed the neural response to the event is 500 ms long. We modeled the strength of the neural response using a Gaussian distributed code amplitude of mean 30 with variance 2. Given the code and kernel, the firing rate of the neuron was constructed based on the DUNL generative model using the Binomial distribution. The test set consisted of 100 trials following similar statistics to the training set. For a low background firing rate, a few spikes were observed in each trial (e.g., for 2 Hz, only 29 spikes were observed on average in each trial, whereas for 17 Hz firing rate, 202 spikes were observed on average in each trial). Hence, learning and inference were challenging when the neuron was very silent.

#### Simulation study II: fitting using DUNL

We considered two scenarios: a) known timing of events (known support), and b) unknown timing of events with a known number of events (unknown support) (Figure S15). The dopamine (Figures 2 and 3) and olfactory (Figure 5A-F) experiments from earlier sections correspond to *known support* scenario, while the whisker deflection (Figure 4) and multi-fiber photometry (Figure 5G-I) experiments corresponds to *unknown support*. When the onsets were known, the inference was reduced to estimating the amplitude of the sparse codes and the shape of the kernel was learned during training. When the onsets were unknown, the inference was more challenging: it involved estimating the event onsets in addition to the neural strength response (the code amplitude) and the kernel shape. In this case, the reliability of learning the kernel was entangled with the reliability of estimating the event onsets.

We showed that when the event onsets are known (Figure S14B top), DUNL’s kernel recovery is relatively robust to the baseline firing rate and can successfully be achieved with few trials (e.g., 25 trials). In this setting, high-temporal resolution (e.g., 5 ms bin size) can be used, regardless of the size of the data. If data are very limited (e.g., 25 trials), increasing the bin size slightly (e.g., 5 ms to 10 ms) is important to implicitly learn a smoother kernel (Figure S15A) (we note that one can also tune the kernel smoothing hyperparameter in the DUNL training framework for better results with very limited data). When the event onsets are unknown (Figure S14B bottom), the bin-size imposes a limit on how well the kernel can be learned (Figure S15B). This challenge comes from the fact that the lower the bin size, the harder the event detection (Figure S14A). We recommend using a larger bin size to match the user’s tolerance for event detection errors and temporal definition. In summary, the higher the number of trials, the higher the firing rate, and the larger the bin-size, the better DUNL’s ability to learn kernels and infer event onsets. Our analyses can help practitioners explore in which regime their experimental data lies and assess which parameters of the model can be recovered from the data.

#### Simulation study III: data generation

In this scenario, we simulated 1000 neurons, each with spiking activity measured during 6-second trials. Up to 1000 trials were run, with neurons responding to 5 distinct events occurring randomly in time. These events stimulated the neurons’ firing rates locally, using a set of 5 orthogonal kernels (Figure 7A,C). While orthogonality is not required by DUNL, it was chosen to allow a one-to-one comparison with Sparse Component Analysis (SCA, [33]) applied to the same dataset. Since events occur randomly in each trial, event alignment across trials is not applicable. Therefore, the neural population data exhibits only a *local* low-rank structure, limiting the use of conventional methods designed to recover global low-rank structures (e.g. PCA [25, 26], NMF [27], GPFA [28], vLGP [29], mTDR [30] TCA [31, 32], SCA [33]).

#### Simulation study III: fitting using DUNL

Using this data, we demonstrate that DUNL effectively learns the local low-rank structure, successfully recovering the 5 underlying kernels (Figure 7B) and decomposing single trials into their local components, even when two kernels overlap in time (Figure 7C,J). When the number of kernels is unknown, the training and validation loss over epochs can guide users to determine the correct number (Figure 7D,E), as the loss plateaus once the kernel count exceeds the true underlying rank (i.e., 5 kernels). Additionally, kernel consistency is highest when the correct number (5) is chosen (Figure 7F), helping users identify the most stable kernel count in experimental data. Moreover, despite the test ***R***^2^ increasing with additional kernels, the slope of this increase flattens significantly after reaching 5 kernels (Figure 7G). We also observe that the maximal cross-correlation between estimated kernels exceeds 0.5 when more than 5 kernels are used (Figure 7H), suggesting redundancy as DUNL learns highly correlated kernels that likely characterize the same event. DUNL remains robust to noise, recovering kernels across a range of spiking additive noise levels (Figure 7I, Figure S16).

There is a large body of work on stable recovery of sparse representation from noisy data [96– 98] but DUNL’s robust performance in this dataset is also due to its ability to share kernels across the neuron population, where group sparsity outperforms individual sparsity [99–101]. Since DUNL is fast and scalable (see Figure S19), it is not burdensome to iterate over different kernel counts to find the best decomposition. Furthermore, in datasets where local kernels are unaligned and lack a shared sequential pattern across trials, global low-dimensional methods like LFADS [38], TCA [102], and SCA [33] fail to detect latent factors (Figure 7J,K). Finally, peak-finding algorithms are inadequate for differentiating between event types in this experiment. For more details and the effect of unrolling iterations on DUNL’s performance, see Supplementary Materials and Figure S17.

### Comparison with other decomposition methods

DUNL is a versatile deconvolutional method that can extract directly interpretable latent representations from time-series data. Its main strengths are its ability to learn multiple local kernels within single trials, i.e., its locality, its capacity to do so in a limited data regime, and its scalability. To emphasize how our framework fills a gap in the space of functionalities provided by other decomposition methods, we evaluate DUNL’s performance to those, while for each comparison, we focus on one specific aspect.

#### Learning temporally low-rank structures

We first focus on DUNL’s ability to learn temporally local figures. We use a synthetic dataset to show that LFADS [38], a deep learning framework for inferring latent factors from single-trial neural activity, does not solve the same problem as DUNL, despite the fact that both are deep-learning-based methods for single-trial analysis of neural data.. LFADS [38] is developed for population analysis of simultaneously recorded neurons to capture dynamics covering the whole trial length; hence, it is not suitable for the deconvolution of asynchronous, local activity within single trials of single neurons. We design the experiment in its simplest form, where the trials are time-aligned and structured, with two different types of non-overlapping events occurring in single trials. We evaluated the interpretability of DUNL and LFADS from their ability to recover the local structure and deconvolve the single-trial neural activity into two traces, each corresponding to one underlying event type. We generated such trials for a training dataset and a test dataset (Figure 8A).

**Figure 8.**
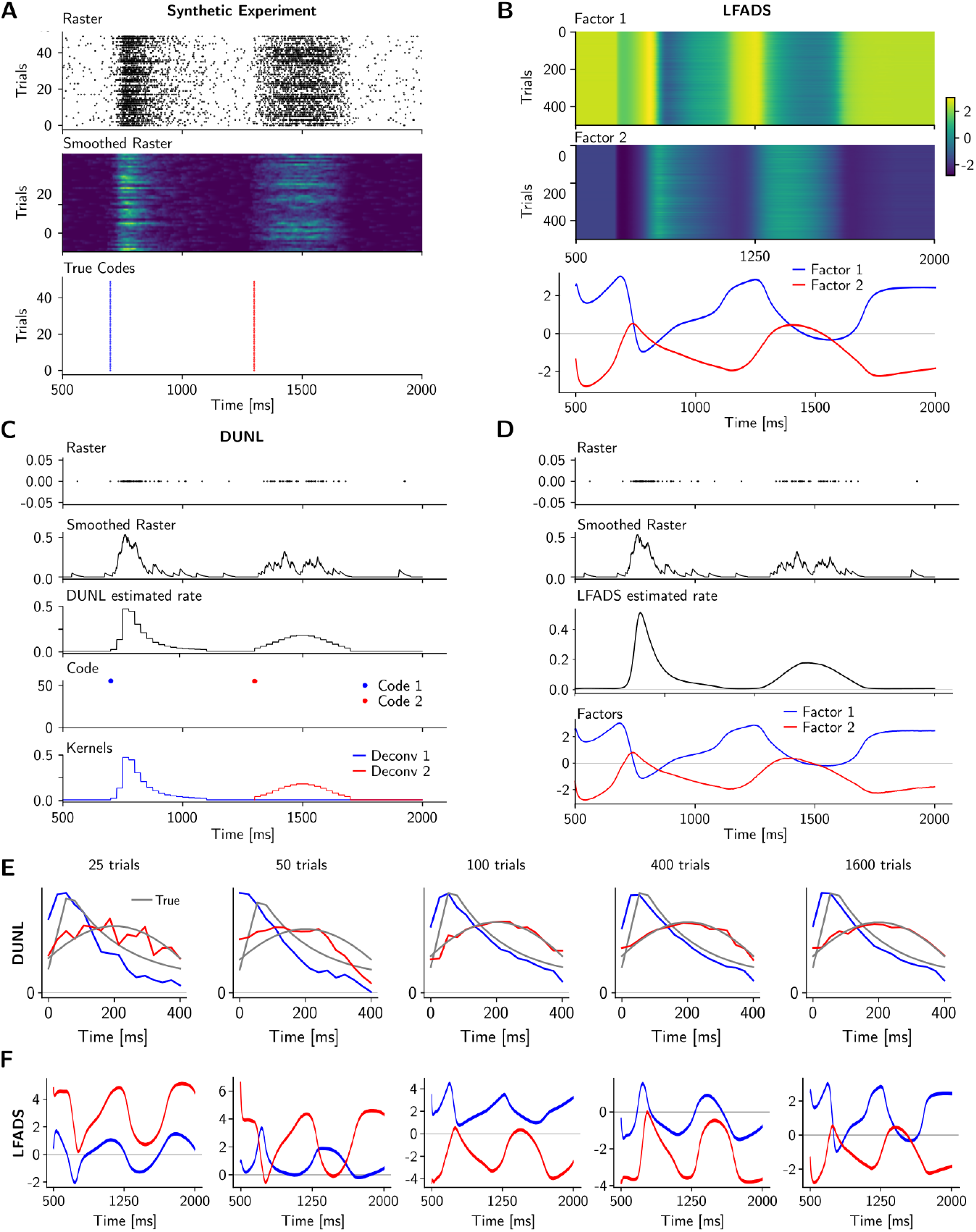
Comparison of DUNL with LFADS. DUNL finds local components in single trials. (A) Experimental setting implemented for simulated spiking neural activity: this is a multiple trial-based experiment in which two events occur in single trials (top and middle). The event onsets for the two events in each trial of the test set (bottom). (B) LFADS estimates two factors for this dataset, which span the entire trial duration. (C-D) In an example trial (raster and smoothed rate on top), DUNL estimates the firing rate of this simulated neuron in 25 ms bins ((C), middle) while LFADS estimates the firing rate in the original rate ((D)), middle). DUNL finds local codes for the two kernels ((C), bottom), while LFADS finds two kernels that span the entire trial duration ((D), bottom). We note that DUNL has estimated the firing rate using 25 ms bin resolution, while the visualization is shown at the original sampling rate at 1 ms resolution; hence, the estimated rate is shown as piece-wise constant at the high-resolution time scale. (E) Kernels found by DUNL using 25, 50, 100, 400, or 1600 trials for training (compared to the true underlying kernels used to generate the data in gray). (F) Average LFADS factors across the training scenarios.

Both DUNL and LFADS were able to estimate the underlying firing rate of the simulated neuron (Figure 8B,C,D). DUNL successfully recovers the underlying kernels (Figure 8E) Despite their success in rate estimations, LFADS: a) finds factors that span the entire trial duration, lacking the locality provided by DUNL, and b) fails to deconvolve the neural activities excited by events of different types from one another (Figure 8F). Overall, unlike DUNL, which provides a functional relation between kernels and firing rates of neurons via a probabilistic generative model, LFADS inference is based on a recurrent neural network, whose encoder and decoder are not tied to one another, thus lacking a direct link between spiking data and the latent factors of single neurons (for an additional comparison of DUNL with SCA [33] to emphasize learning temporally low-rank structures see Figure 7K). Overall, The two examples demonstrate the ability of DUNL to perform local characterization and deconvolution in single neurons.

#### Scalability

DUNL can handle trials of varying lengths due to its convolutional and batch-based inference approach. It learns via accelerated gradient-based optimizers like Adam [103] and is scalable, running batches of data on GPU. To demonstrate DUNL’s scalability, we compared its training runtime with two methods: SCA [33] and seqNMF [37]. Both methods only handle data in a (Neurons × Trial Length) matrix format, limiting their ability to analyze trials of different lengths or multiple trials without concatenation. While SCA can run on GPUs, its full-batch approach allows fitting a maximum of 8 trials, and it takes significantly longer to train than DUNL (Figure S19A). SeqNMF, running on CPU, can fit more trials but still requires much more time than DUNL. Beyond scalability, seqNMF is designed to sort neurons based on population response structure. However, it struggles to deconvolve local kernels that span the same neurons but appear separately in time. In experiments requiring the detection of two temporally local kernels, seqNMF identified only one, and its code did not align well with event onsets, failing to infer a highly sparse representation despite the sparse regularization provided (Figure S18).

#### Decomposition property

Next, we highlight DUNL’s kernel learning capabilities and demonstrate how accounting for the statistical properties of the data (e.g., binary spiking data) allows DUNL to learn more interpretable kernels compared to classical and deep dimensionality reduction methods. To focus on kernel learning, we applied the baseline to local data windows, bypassing the sparse coding/event detection step. Specifically, we analyzed 600 ms windowed dopamine spiking data, time-aligned to the reward onset. The result showed that Principal Component Analysis (PCA), and Non-negative Matrix Factorization (NMF) can be used to extract components from windowed data (Figure S20A,C). They do not offer the salience/value interpretability that DUNL provides. The coefficients extracted from each trial and the Spearman’s rank correlation between each neuron’s coefficients across all trials are not aligned with the decomposed codes from DUNL (Figure S20B,D). For PCA, this is due to the dissimilarity of the learned kernels to what we know from the salience and value responses. For NMF, the kernels are semi-similar to the learned kernels in DUNL (non-negative kernels are learned in the spiking scenario). However, the NMF coefficients are not capable of capturing the dip in neural responses due to their non-negativity constraint, resulting in a lower Spearman’s rank correlation to the reward sizes. Finally, we note that one may use Poisson-functional PCA [104] to improve the decomposition interpretability of PCA for spiking data. However, we note that the advantages of DUNL go beyond decomposition property; DUNL is scalable and can find locally low-rank structures and can do event detection, properties that are missing in PCA, NMF, and Poisson-functional PCA [104].

Additionally, we again compared DUNL with LFADS using the same window of dopamine spiking data. Using two factors, we found that despite LFADS’s success in estimating the firing rate of neurons in each trial, the learned factors lack salience-value interpretability (Figure S20E). The comparison of the Spearman’s rank correlation analysis on the LFADS factors and DUNL’s codes for the reward response (Figure S20F) further supports the absence of salience-like characterization in the LFADS method: both factors incorporate value information. These effects could be due to the more expressive architecture of LFADS that overfits the data at the expense of a parsimonious and interpretable description (See Figure S20G-H in Supplementary materials for LFADS analysis in the limited data-regime).

#### Choice of Basis

Finally, We highlight that DUNL allows users to *learn* kernels using sparse representations to characterize neural responses, addressing two key points: a) it eliminates the need for domain knowledge in selecting a set of basis functions, and b) it provides a highly sparse set of kernels (e.g., 1 or 2 per event, unlike time-delayed bases), enhancing the interpretability of representations. In this regard, we applied GLM with a family of basis functions [34] using a similar time-bin resolution of 25 ms as DUNL to the dopamine spiking experiment. We guide the frame-work to fit the data using a set of bases at the onset of the reward for a duration of 600 ms. We show results from two scenarios, each with a different set of bases, i.e., non-linear raised cosines and raised cosines. We found that although GLM with pre-defined bases has smooth fitting curves, it cannot deconvolve single-trials into components that are interpretable from the perspective of salience and value in this experiment (Figure S21). We argue that this is primarily due to the dissimilarity of the pre-defined bases to the kernels learned by DUNL. One may use the DUNL kernels within the GLM framework to perform the deconvolution of interest, thus taking advantage of the interpretability of learned kernels in place of pre-defined bases. However, without a kernel-learning framework, such interpretable kernels are unknown a *priori*. Overall, in this setting, DUNL shares similarities with approaches like GP-GLM, where a temporal set of kernels is learned rather than designed [94].

## Discussion

We introduce the use of unrolled dictionary learning-based neural networks [73–75] to recover locally low-rank structures and deconvolve multiplexed components of neural activity that are relatable to human-interpretable latent variables. This is a technique, based on algorithm unrolling [57, 58], to design an interpretable deep neural network.parUsing this technique, we show for the first time that temporal convolutional sparse coding techniques can be used in neural recordings to detect and characterize sparse latent events on time scales from millisecond to second. Our method, DUNL (Deconvolutional Unrolled Neural Learning), fulfills important desiderata of a decomposition method: it can be implemented in single instantiations of the neural data, it can be trained with a limited dataset, it is flexible in regard to the source signal, it generates a mapping between data and latent variables, and, importantly, from these latent variables to human-interpretable variables. This is achieved through the use of a generative model that guides the architecture of the inference deep neural network during the optimization process (Figure 1) and the choice of that generative model, i.e., convolutional dictionary learning to capture the data statistics. Consequently, domain knowledge is incorporated into the deep neural network. DUNL is a deconvolutional method that can look for and encode overlapping local events within a single trial. The end goal might be to detect/characterize either localized asynchronous events in single-neurons, or localized synchronous events across the population (e.g., cue and reward components in the dopamine experiments or multiple deflections in the whisker experiment). This goal contrasts with most neural data analysis tools, which typically seek latent representations in the neural population of simultaneously recorded neurons, functioning as dimensionality reduction methods across the entire trial duration. As a result, many classical methods and their successors—such as PCA [25, 26], sparse coding [105], SCA [33], NMF [27], or LFADS [38, 39]—provide components or factors that cover the whole trial or sequential activity in the neural population. Compared to other convolutional methods like convNMF [35, 36] and seqNMF [37], which focus on local characterization of neuron-by-time matrices, DUNL stands out for its scalability and ability to analyze data tensors across neurons, time, and trials. (See Supplementary Discussion for a detailed comparison of DUNL and prior works.)

Our work, while sharing the convolutional property with a subset of these methods, and the sparsity and group sparsity constraint with another subset, adds in the statistical nature of generalized linear models (GLM) [34, 94], and the use of deep learning. This combination enables: a) finding locally low-rank structures by learning kernels (covariates) from the data and b) using deep learning and backpropagation for data fitting, such that the typical response function of neurons and their amplitudes to each event in single trials are directly obtained from the network weights and latent representations. Our method is particularly useful when multiplexed signals are encoded by individual neurons or populations, and event detection is needed, especially when users prefer not to impose constraints on the kernels (such as orthogonality in PCA, non-negativity in NMF, or a predefined basis set in GLM). However, constraints and regularization (e.g., enforcing or discouraging kernel co-activation, non-negativity, group sparsity, or smooth kernels) can easily be added to the optimization problem if needed.

Our method owes its efficiency and scalability to the combination of algorithm unrolling with convolutional sparse coding to provide temporally low-rank structure to the analysis of the neural data. Exogenous stimuli, behaviour, and neural activity all share time as a fundamental variable, and sparse coding has a rich history in neuroscience as an interpretable theory of early sensory processing in many brain regions and systems [106, 107]. By adding temporal structure to sparse coding, we obtain an expressive artificial neural network that can deconvolve single-trial neuronal activity into interpretable components, because they correlate with exogenous stimuli and/or behaviour. First, the obtained latent representations are aligned with time and, second, they are sparse. Our method’s deconvolution of neural response components can be seen as resulting in an input/output characterization of the functional properties of a system (neuron(s) in this case).

The appeal of approaches such as GLMs comes from the fact that, in some sense, they provide such input/output descriptions of neural responses. Signal processing theory [108] has well-established links between such descriptions and computations (e.g., differential equations). The GLM-like statistical nature of our models linking latent, learnable, representations and neuronal data, together with their translation, via algorithm unrolling, into interpretable deep-learning architectures, leads to a powerful approach for analyzing single-trial neural data that satisfies the desiderata put forth. Our framework is designed to be flexible, which requires setting a few hyperparameters (e.g., sparse regularizer, number of unrolling iterations, smooth regularization, number of kernels). We provide guidance for selecting these based on the data. For example, to determine the right number of kernels, users can evaluate the consistency of output kernels across different initializations and quantify the maximal cross-correlation between any two kernels. Importantly, only the number of kernels and time window duration need to be specified – no other parameterization is required, as DUNL can learn any kernel shape unlike previous methods [34], but see [94]. When the approximate number of events is known, DUNL employs a k-sparse operation within its unrolling architecture, making it relatively robust to choosing the sparsity parameter *λ*. Without the k-sparse operation, a validation set can be used to tune *λ*.

One limitation of convolutional sparse coding methods is that two kernels may not be distinguishable if they are perfectly co-active and have similar shapes; following Occam’s razor, one may choose to model them as a single kernel. Additionally, in convolutional dictionary learning, higher kernel correlation makes a recovery more difficult [109]. While kernels do not need to be orthogonal, more orthogonality facilitates the learning and inference process [75, 109, 110]. Although DUNL can recover relatively coherent kernels when initialized randomly, theoretical analyses of dictionary learning rely on an incoherence assumption and a relatively good kernel estimate at initialization [110, 111]. The current design of DUNL infers representations using kernels of the same length for a single data modality and cannot perform time-series prediction. Additionally, DUNL cannot yet handle multiple recording modalities simultaneously, though this is feasible in principle. Future work could extend DUNL to learn representations shared across multiple modalities using multi-scale kernels or use the unrolling technique more broadly on a different iterative algorithm [57]. While prior literature addresses missing data through dictionary learning from incomplete data [112, 113], and recent work explores unrolled dictionary learning from compressed Gaussian-distributed measurements [114], DUNL does not currently support handling missing samples, but this could be implemented in future versions.

To show the versatility of our model, we applied it to a diverse set of experimental settings in which we highlight functionalities that our method provides and that where not available with previously published tools. To conclude, we point out that the unrolling framework can be extended to provide interpretable latent representations under other regimes, besides the sparsity one used here. More complex generative models can be used, for instance, Kalman filtering-based neural networks [57, 65], 2D filters with other constraints. Our work is a first step towards leveraging the advances in interpretable deep learning via convolutional dictionary learning to gain a mechanistic understanding of neural dynamics and underlying computations.

## Acknowledgment

We thank the reviewers for their constructive feedback. We thank Ignacio Sanguinetti-Scheck for conceptual and technical collaboration in the dopamine recordings in naturalistic conditions. We thank all members of the Uchida, Murthy, and Ba laboratories for helpful discussions, particularly Jacob Zavatone-Veth, Adam S. Lowet, and Andrew H. Song for feedback on the manuscript. We thank Gil Costa and Scidraw.io for the rat schematic (doi.org/10.5281/zenodo.3926343). We thank the funding agencies that supported our work: the US DOD, UK MOD, and UK Engineering and Physical Research Council (EPSRC) for the ARO Grant (W911NF-16-1-0368 to DB), the NIH (grant 5R01DC017311 to NU and VNM), the Human Frontier Science Program (LT000801/2018 to SM), the Harvard Brain Science Initiative (Young Scientist Transitions Award to SM); the Brain and Behavior Research Foundation (NARSAD Young Investigator Grant no.30035 to SM); and the Harvard Mind Brain Behavior Interfaculty Initiative (to PM). This research was undertaken thanks in part to funding from the Canada First Research Excellence Fund, awarded to PM through the Healthy Brains, Healthy Lives initiative at McGill University.

## Author contributions

Author contributions are summarized in the table.

**Table 1.** Author contributions

| Authors | BT | SM | HW | ST | NU | VM | PM | DB |
| --- | --- | --- | --- | --- | --- | --- | --- | --- |
| Study conception | ✓ | ✓ |  |  |  |  | ✓ | ✓ |
| Methodology | ✓ |  |  |  |  |  |  | ✓ |
| Formal analysis | ✓ | ✓ |  |  |  |  |  |  |
| Investigation: performed *in silico* experiments | ✓ |  |  |  |  |  |  |  |
| Investigation: performed *in vivo* experiments |  | ✓ | ✓ | ✓ |  |  |  |  |
| Data curation | ✓ | ✓ | ✓ | ✓ |  |  |  |  |
| Writing - initial draft and final manuscript | ✓ | ✓ |  |  |  |  | ✓ | ✓ |
| Writing - critical review and revision | ✓ | ✓ |  | ✓ | ✓ | ✓ | ✓ | ✓ |
| Writing - data presentation | ✓ | ✓ |  |  |  |  |  |  |
| Supervision |  |  |  |  |  |  | ✓ | ✓ |

## Supplementary Materials - Methods

### Notation

Scalars are denoted by non-bold-lower-case *a*. Vectors and matrices are denoted by bold-lowercase ***a*** and upper-case letters ***A***, respectively. We let *n* = 1, …, *N* index neurons, and *j* = 1, 2, …, *J* the number of trials. We assume that the time series representing the activity from neuron *n* at trial *j* comprises *T* measurements, which we denote ***y***^*n,j*^. We denote the full measurement tensor by ***Y*** with *T* × *J* × *N* dimensions (time/bin measurements, number of trials, number of neurons). We note that the tensor structure of the data is for ease of notation and the proposed framework can work with measurement data ***Y*** of varying size across the time dimension *T* on a batch basis. We denote the convolution operator by ∗ and its transpose operator (correlation) by ⋆. Finally, we use superscript T for transpose of a matrix ***A***^T^.

### Data Distribution

For spiking data, the spikes at each trial are binned at *B* ms resolution. Hence, each entry of ***y***^*n,j*^ represents a spike count ranging from 0 to *B*. We model the observations using the natural exponential family [68, 69], i.e., ***y***^*n,j*^ ∼ Poisson(***μ***^*n,j*^) and ***y***^*n,j*^ ∼ Binomial(*B*, ***μ***^*n,j*^), where ***μ***^*n,j*^ models the mean of the distribution for neuron *n* at trial *j*. For continuous-valued data, such as Calcium fluorescence data, we model the time series ***y***^*n,j*^ ∈ ℝ^*T*^ as a Gaussian distribution with mean ***μ***^*n,j*^. We construct the data log-likelihood of the natural exponential family as [68, 69]

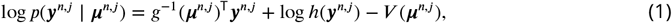

where *h*(***y***^*n,j*^) is a non-negative function as the base measure. Condition on *μ*^*n,j*^, we assume the entries of ***y***^*n,j*^ are independent. The functions *g* (i.e., inverse link), and *V* depend on the particular choice of distribution (see Table S1).

**Table S1.** Natural exponential family data log-likelihood specifications.

| | $\mathbf{y}$ | $V(\mathbf{z})$ | $g(\cdot)$ |
| --- | --- | --- | --- |
| Gaussian | $\mathbb{R}$ | $\mathbf{z}^\top \mathbf{z}$ | $I(\cdot)$ |
| Binomial | $[0 \dots B]$ | $-\mathbf{1}^\top \log(\mathbf{1} - \mathbf{z})$ | $\text{sigmoid}(\cdot)$ |
| Poisson | $[0 \dots \infty)$ | $\mathbf{1}^\top \mathbf{z}$ | $\exp(\cdot)$ |

### Generative Model

We follow the perspective of analysis-by-synthesis [24] and Bayesian generative modelling [71].

For each neuron *n*, we impose a generative model on the neuron’s activity (i.e., the firing rate in the spiking setting) and model it as a function of a baseline mean activity level *a*_*n,j*_ and a set of *K* localized kernels 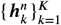 characterizing the neuron’s response to events that occur sparsely in time. We let the sparse vector 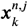 encode the onsets of events associated with the kernel *k* in trial *j*: its nonzero entries represent the times when events occur, and their amplitude the strength of the contribution of the *k*-th kernel to the neuron’s response. Similar to ***y***^*n,j*^, the entries of the sparse code 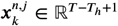 and the filter 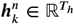 are both indexed across time. We note that *T*_*h*_ is chosen based on the locality of the neural responses and the sampling rate. *T*_*h*_ is not a function of *T* ; an increase in duration of the trial should not affect the length of kernel *T*_*h*_.

Mathematically, we can express this *convolutional sparse coding* model as follows

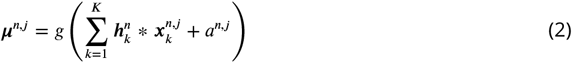

Although the model results in an estimate of each neuron’s firing rate on a trial basis, the kernels capture characteristics that are shared among trials and can be distinct across neurons or shared across the neural population. At times, we may use the terminology dictionary element to refer to the kernels. For the scenario where we share the kernels across neurons, we simplify the dictionary notation to ***h***_*k*_.

### Smooth Sparse Deconvolutional Learning

#### Optimization

Given the set of observations from all trials 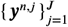 for each neuron *n*, we learn the kernels and codes by minimizing the negative log-likelihood with a sparse prior on the codes, i.e.,

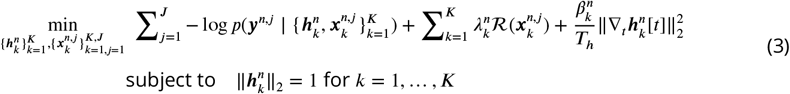

where 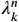 controls the regularization of the codes For example, setting 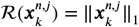 regularizes the sparsity of the codes (i.e., the frequency of onsets in time) for the kernel (event-type) *k* and neuron *n*. This regularization can also be defined as a group sparsity [99–101] at each time step (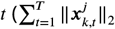 where 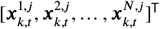) across neurons to encourage neurons to fire together and detect events concurrently with one another. Moreover, 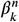 controls the smoothness of the kernels, achieved by regularizing the first derivative of the kernel with respect to time samples *t* [72, 115]. We use the general notion of sparsity to refer to both sparsity and group sparsity in the text, and call the above optimization smooth sparse deconvolutional learning (SSDL).

SSDL is a variant of dictionary learning, also referred to as sparse coding, and has widespread application outside of neuroscience. Dictionary learning is widely known in statistics and signal processing communities [116, 117]. The sparse coding model was initially introduced by Olshausen and Field [105] to model early layers of visual processing. Prior works used sparse coding for modeling neural connectivity and dynamics of early sensory systems [106, 107, 118–120]. Moreover, for imaging transcriptomics, the model is used to learn representations of gene expression [121, 122]. Within the neuroscience literature, sparse coding also appears in different contexts for analysis of EEG signal [123], calcium imaging [124, 125], auditory encoding [126], and spike sorting [127, 128], neuronal assembly detection [129], to name a few.

#### Alternating minimization

Equation (3) is a bi-convex optimization problem and can be solved by an iterative alternating-minimization algorithm [110]. Letting *l* denote its iterations, the algorithm alternates between a *sparse-coding* step, that computes an estimate of the codes 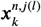 given an estimate 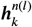 of the dictionary, and a *dictionary-update* step, that uses this new estimate of the codes to obtain refined estimates 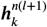 of the kernels. Mathematically, we can express the two steps as follows

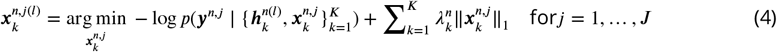

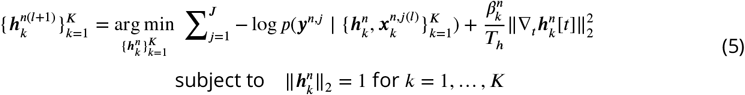

The first step explains the sparse coding step which is separable over trials and neurons. For group sparsity where the code is jointly optimized across neurons, the step optimizes a modified objective as following

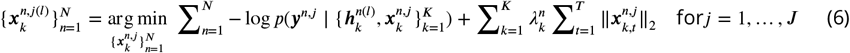

Compared to classical sparse coding, both our sparse-coding and dictionary-update steps have convolutional structure. Intuitively, the convolutional structure of our model enables the identification of patterns that occur *across* time.

#### Sparse coding step

The sparse coding step can be solved using the iterative shrinkage-thresholding algorithm (ISTA) [130, 131] where we have implicitly absorbed the noise variance into the sparse regularizer. One iteration of this proximal gradient descent algorithm proceeds as follows

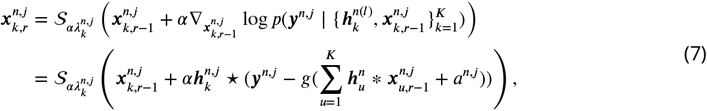

where the so-called shrinkage operator 𝒮 _*b*_(*z*) ≜ sign(*z*) max (|*z*| − *b*, 0) is a nonlinear, sparsifying, thresholding operation, and *r* denotes the sparse coding iteration *r*. For non-negative sparse coding, i.e., when the entries of the sparse code can only take a non-negative value, the shrinkage operation 𝒮 _*b*_(*z*) reduces to the celebrated ReLU_*b*_ (*z*) = (*z* − *b*) · **1**_*z*≥*b*_ nonlinearity. The converged code estimate from the iterative update in (8) is a minimizer of the sparse coding step (7). For group sparse coding, the nonlinear activation function takes the form 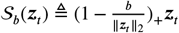 where ***z***_*t*_ is the code vector from each defined group at time *t* across neurons and (·)_+_ = *max*(·, 0). In applications where the onsets of events are known (i.e., code support is given), we apply an additional indicator function of events 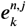 at every iteration. Thus, the iterative updates compute estimates of the strength of *k*-th kernel contribution to neural activity at known event-onset times.

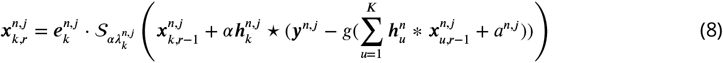

#### Dictionary learning step

We use gradient-based methods to update the dictionary. In its simpler form, the update is of stochastic projected gradient descent.

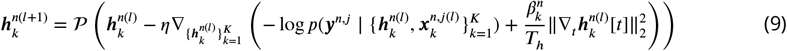

where 𝒫 (***z***) = ***z***/ ‖***z***‖ _2_ performs a norm projection, and η is the learning rate. This step concludes SSDL formulation and optimization. We note that SSDL, especially for non-Gaussian distributed data, is not a published framework, and there are no publicly available optimization toolboxes that can solve SSDL. Given the scalability challenges of current toolboxes, we next discuss how the optimization of SSDL can be mapped into the training of a deep neural network, which we call deconvolutional unrolled neural learning (DUNL). We note that the transition from SSDL to DUNL is key for a) the practical applicability of the proposed framework on large-scale neural data and b) the flexibility of the framework to encourage/impose a variety of constraints on parameters and representations (e.g., input event onset, structured representation, *k*-sparse codes).

### Interpretable deconvolutional unrolled neural learning (DUNL)

#### Inference network

The alternating minimization procedure explained above can be mapped into an encoder/decoder neural architecture. Specifically, we use algorithm unrolling [57] to map the sparse coding step (4) into an encoder. This is similar to the network architectures proposed in [68, 128] for dictionary learning. In this architecture, each sparse coding iteration (7) is interpreted as one layer of a neural network with a particular recurrent convolutional structure and shrinkage or ReLU non-linearity. We refer to this encoder as an inference network that maps the single-neuron, single-trial time series ***y***^*n,j*^, into estimates of the time series sparse codes 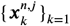,encoding event onsets and their contribution to explain the data (see Figure S1a).

**Figure S1.**
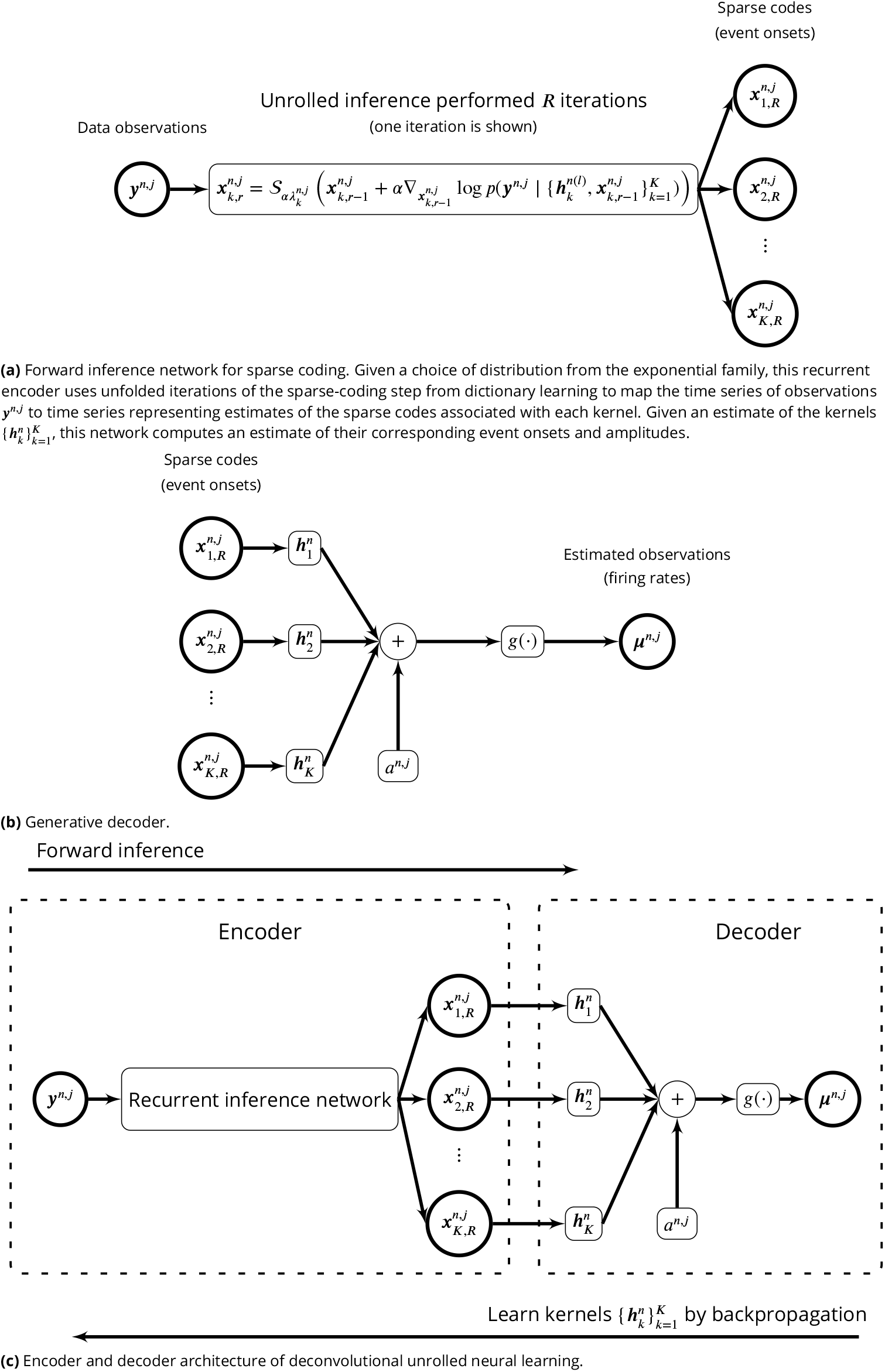
Deconvolutional unrolled neural learning (DUNL).

#### Generative decoder of DUNL

Given the codes from the inference network, we construct a decoder based on the generative model (2) (Figure S1b). This decoder maps the estimated time series of sparse codes from a given neuron into a time-series observation estimate (e.g., a time series representing firing rate in the case of spiking data).

#### Deconvolutional Unrolled Neural Learning (DUNL)

We combine the inference network and the generative decoder to construct an interpretable network which can be trained by backpropagation. Training lets us learn the kernels 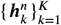 that characterize the neural response to events coded by 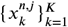. As detailed in the introduction, The interpretability of this network is two-fold: the network trainable parameters are directly related to the Kernels 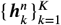, and the encoder latent representation corresponds to the event onsets and their strengths.

Training this network involves both a forward pass (inference) and a backward pass (training to learn the dictionary), both of which are embarrassingly parallelizable over neurons and trials. Therefore, the interpretation of the sparse-coding and dictionary-update steps as a network enables to take advantage of the parallelism offered by GPUs seamlessly. The benefits of DUNL compared to SSDL extend beyond GPU utilization. Convergence analysis of DUNL, when kernels span the entire trial and in the Gaussian case, has shown that with limited unrolling layers (approximate sparse coding), DUNL recovers the underlying dictionary better than SSDL [75]. In this Gaussian setting, DUNL also reduces estimation bias in model recovery compared to alternating-minimization-based optimization [75]. These advantages have been demonstrated in spike sorting [128], and theoretical analysis exists for the Binomial case with one unrolled layer [68]. Additionally, DUNL benefits from insights in sparse coding, which has a rich literature on its performance in the presence of model mismatch [132], noise [96–98], and adversarial attacks [133]. For example, [68] studies model mismatch where spiking data is modeled as Gaussian instead of Binomial, showing that while mismatch increases recovery error, the learned parameters still correlate positively with the ground truth.

#### Structured representation

Motivated by biological constraints/prior knowledge, we may want to impose structure on the codes in addition to sparsity. Calcium fluorescence, for instance, does not encode electrical activity linearly: the signal exhibits different dynamics when the firing rate of a neuron increases than when it decreases. Even though the dynamics of the underlying firing rate might be the same, the measured calcium signal will be different, but these dynamics can be captured through the addition of structured representations in our framework. Consider a version of our model for fluorescence data, with one kernel for each neuron. This model could not capture such a nonlinear relationship because both positive (increased activity above baseline) and negative (decreased activity) codes would need to use the same filter/kernel. To overcome this challenge, we can introduce an additional filter and a prior on the codes of both filters that prevent the co-occurrence of event onsets, i.e., that prevents both filters from contributing to neural activity at the same onset times. As we will demonstrate in our analyses of fluorescence data from dopamine neurons, such priors allow us to capture the nonlinear relation between activity level and fluorescence. We can either enforce such latent structure on the codes or learn it by incorporating an additional term into the original optimization. Mathematically, our modified optimization solves

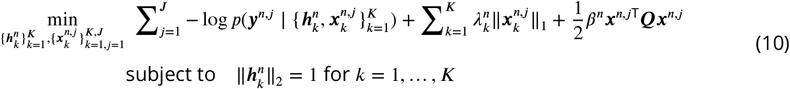

##### Algorithm 1

Iterative power method to approximate unrolled step size α.

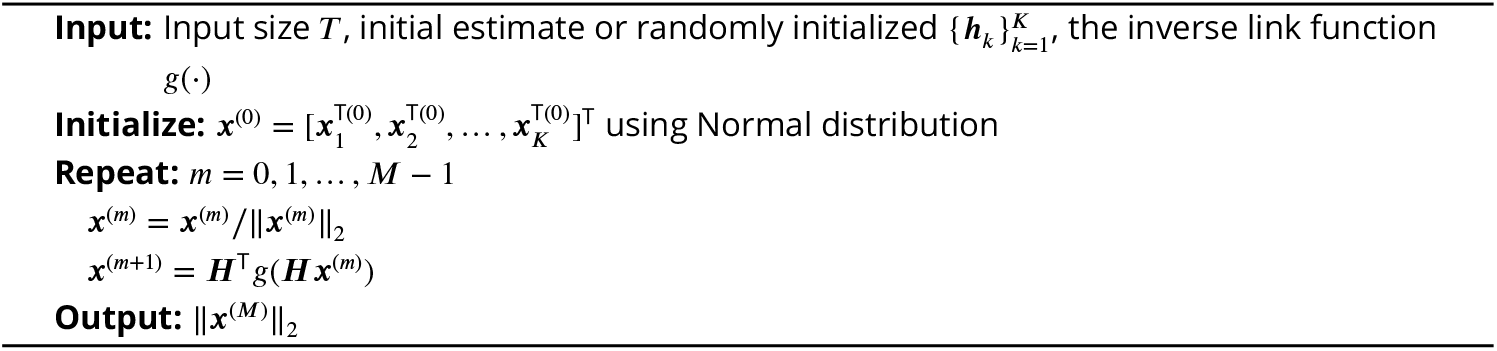

where 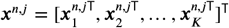,and 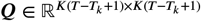 is a symmetric matrix with block structure. For example, given two kernels (*K* = 2) when codes are non-negative, 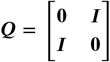 enforces a structure such that the kernels ***h***_1_ and ***h***_2_ are discouraged to get activated simultaneously. Variations of such latent regularization are 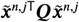 where ***Q*** ∈ ℝ^*K*×*K*^, and 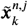 captures the energy of the code 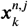 (e.g.,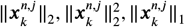, etc.). Although we treat ***Q*** as a hyperparameter, it can be related and is proportional to the negative inverse covariance matrix of the code ***x***^*n,j*^, hence it can be learned. Overall, this regularization modifies the recurrent inference network to

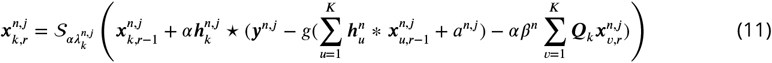

where ***Q***_*k*_ is the *k*^th^ column block of ***Q***. Our source code is written such that this regularization can be enforced not at every unrolled layer but with a chosen period. In addition to the residuals, the kernel codes are now interconnected to one another through ***Q***. From (11), we see that the amplitudes of 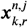 is damped when ***Q***_*k,v*_ is positive, and 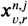 has high activity. In the above coding formulations, we assume that the baseline activity *a*^*n,j*^ is given by the user (e.g., pre-estimated from the beginning of the trial). However, DUNL also has the option of inferring the baseline activity of each neuron at each trial within its unrolling encoder; in this case, the baseline is updated at each layer of the encoder as follows:

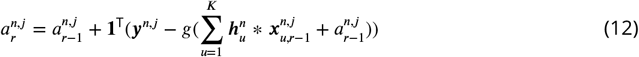

where **1**^T^ computes the average of the residual across time, and in (11), 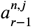 is used, accordingly.

#### Network parameters

In this section, we explain the network parameters and the effect of each on the performance of DUNL.

### Unrolled step size α

This is the step size inside the unrolled network. For stability purposes, α < 1/*σ*_max_(***H***), where *σ*_max_ is the maximum singular value, ***H*** = [***H***_1_ |***H***_2_ |***⋯*** |***H***_*K*_], and ***H***_*k*_ is the linear Toeplitz matrix corresponding to the convolution kernels ***h***_*k*_. An upper bound on the step size α can be approximated by the iterative power method shown in Algorithm 1. We note that the performance is not very sensitive to this hyperparameter as long as the network is stable.

### Sparse regularizer *λ*

This is the regularization parameter to enforce sparsity on the latent representation estimated at the encoder. This parameter can be set to 0 or a small value when the event onsets are known and enforced by the indicator 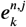 at the unrolled iterations. However, its presence is crucial without known support (event onsets). In the case of group sparsity, *λ* operates on the combined activation of the code across neurons to form a group structure. In this case, increasing *λ* discourages the activation of the code for individual neurons when most neurons are inactive.

### Baseline a^n,j^

Our model assumes that there is a baseline activity constant over time in each trial for each neuron. This can be estimated by taking the mean activity at the beginning of each trial prior to the appearance of events of interest, followed by the link function *g*^−1^(·). Alternatively, the user has the option of inferring the baseline activity within the unrolling encoder along with the code. This parameter is easy to infer or pre-estimated.

### Unrolled layers *R*

The inference network (Figure S1a) is equivalent to the optimization (4) when ***R → ∞***. However, given computational limitation ***R*** is finite. We recommend setting ***R*** on the order of 100 when the code support is known, and 1000 when the support is not known in the case of finding kernels that may show correlation (e.g., see Simulation Study I). However, it can be reduced to lower than 20 when local kernels are incoherent (e.g., see Simulation Study III). We recommend that the user increase ***R*** for improved deconvolution and sparser representation.

### Number of kernels *K*

Domain knowledge can help to set this hyperparameter. For example, if there are 2 distinct event types that are happening locally in time across trials, then it is recommended to set *K* = 2 or overestimate slightly. See Simulation Study III, for tips to choose the suitable number of kernels.

#### Learning parameters

This section explains the parameters involved during the learning/training stage. Additionally, we specify which parameters are hyperparameters (i.e., to be set by the user), which are learned during training, and which are estimated per inference.

### Kernel smoother regularization 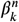

This is the regularization parameter used during training. Larger values encourage the smoothness of learned kernels. Using the kernel smoother regularization is highly recommended for spiking data when the chosen bin size *B* is small [115]. We note that one may reduce the need for kernel smoother by increasing the available data or the bin size for spike count; the latter can help improve event detection at the cost of learning temporally low-resolution kernels. Unlike 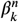,which directly operates on the kernel, the bin size *B* pre-processes the data which is used during training and inference. We recommend setting 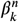 to a small value. A validation set can be utilized to tune this parameter; however, the authors recommend using their domain knowledge to visually tune this parameter to reduce the effect of learning noisy kernels. Additionally, the correlation between kernels of different runs can be used to find the minimum 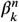 at which the correlation is maximized, indicating learning of stable and less noisy kernels.

### Gradient-based optimizer (ADAM) [103]

DUNL is optimized by gradient-based ADAM optimizer, which is very efficient in finding approximate solutions very fast. This optimizer is mainly designed for deep learning to ensure the optimization does not converge to sharp local minima [134]. Additionally, there is an active line of research on improving generalization properties of ADAM [135]. For DUNL, we have increased one of the optimizer’s parameters *ε* to have better converging solutions. Through our source code, the user can choose other optimizers. Moreover, increasing the batch size or performing fine-tuning toward the end of the training can improve kernel recovery and reduce the error.

#### Evaluation

We note that in simultaneous learning of the kernels and sparse codes in DUNL, it is possible to fit the same neural firing rate with a right/left-shifted kernel in time along with a left/right-shifted sparse code. Indeed, for kernels with decaying ends to baseline, this can frequently happen and the two rate models are equivalent. Thus, when we evaluate DUNL, we account for this by using cross-correlation as a metric for kernel recovery, and by using a tolerance when computing the event detection hit rate.

## Supplementary Materials - Training

### Dopamine spiking experiment

There are *N* = 40 optogenetically identified dopamine neurons. Across neurons, the number of trials ranges from *J* = 121 to *J* = 302. For surprise trials, we analyze the data from 1 s before the reward onset and 2.1 s after the onset. Similarly, for expected trials, we consider the data from 1 s before the cue onset up to 0.6 s after the reward onset. For expected trials, the reward is delivered after 1.5 s from the cue onset. We refer the reader to [87] for more information on data acquisition. We use the Binomial distribution with time-bin resolution of 25 ms for data modeling. We set *K* = 3 to learn three non-negative kernels shared across all neurons and all trials; one to characterize the neural response to the cue, and the other two to characterize salience and value for the reward prediction error responses. Each kernel is 600 ms long in time, and the baseline firing rate *a*_*n,j*_ is estimated (or inferred) for a single trial using the 1 s data prior to the event onset (bins with estimated baseline lower than 0.001 are set to 0.001 for stability purposes prior passing through the log link function. Given the learned kernels, from each neuron at each trial, we infer three codes to identify the neural strength response to cue (*k* = 1) and reward (*k* = 2, 3).

In addition to the norm projection 𝒫 (·), we apply element-wise ReLU_0_(·) projection after every backpropagation (kernel updates) to enforce kernel non-negativity. To enforce the known support for each kernel, the indicator vector for cue code 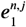 is set to 1 at the cue onset. Similarly, 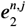 and 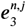 are set to 1 at the reward onset for each neuron *n* at each trial *j* (the event indicators are zero at other time-points). The data, model, and training parameters are summarized in Table S2. For the data-limited scenario, only 685 out of 8,786 total number of trials are used for training; in this case, the kernel smoother penalty is set to 0.0005.

### Dopamine calcium experiment

The data is captured from 3 different sessions, each with *N* = 6, 20, and 30 neurons. Data, model, and training information are summarized in Table S3. Below, we explain the modeling in detail.

Given the continuous domain of calcium imaging, we model the data using Gaussian distribution. We learn *K* = 5 kernels; one kernel to characterize the neural activity in response to the odor cue without reward (we call this regret), another kernel for the odor cue in the expected trial prior to the appearance of the reward, and three kernels to model RPEs. Specifically, for RPEs, we use one kernel with non-negative code to model salience, two kernels to model value (one with non-negative and another with non-positive code). We note that we do not enforce any other constraint for the kernels to explicitly model salience or value; the decomposition is natural upon training. Each kernel is 4 s long in time. The baseline firing rate is estimated from the 1 s data interval prior to the first event onset for every trial.

To attribute the kernels to the specific event of interest, we set the indicator vector for cue regret code 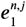 to 1 at the cue onset on regret trials, and zero, otherwise. Similarly, 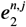 is set to 1 on expected trials at the cue onset, and zero, otherwise. This attribute kernel ***h***_1_ characterizes the neural response to cue in the absence of a reward, and ***h***_2_ represents the neural response to cue in expected trials. Furthermore, 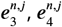,and 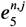 are all 1-sparse for trials with reward, and they are non-zero at the reward onset.

We use the structured representation optimization formulation described earlier to discourage codes ***x***_4_ and ***x***_5_ to be active at the same time. We use 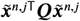 regularization variation with 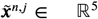 capturing the amplitude of the non-zero entry of each code, i.e.,

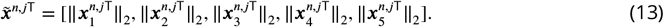

Moreover, we set ***Q***_4,5_ = ***Q***_5,4_ = 2.5 and other entries of ***Q*** is set to 0.

### Whisker thalamus spiking experiment

The training and modeling parameters of the whisker spiking experiment are summarized in Table S4. The original data contains data from 17 pairs of neurons and their activities in response to three types of stimuli. Considered neurons are from pair/neuron of 1/2, 2/1, 4/1, 5/1, 6/1, 8/2, 10/2, 16/2, 17/1 excluding non-responsive neurons or those with very low signal-to-noise ratio [90]. Additionally, the neural characterization is done for stimulus number 3, where the deflection velocity is constant.

**Table S2.** Parameters for dopamine spiking experiment.

| Data |  |  |  |
| --- | --- | --- | --- |
| Sampling rate | 1 ms | Trial length | [121, 302] |
| Number of neurons | 40 | Number of trials | [60-156] |
| Total number of neurons | 40 | Total number of examples | 8,786 |
| Code |  | Kernel |  |
| Non-negativity | False | Non-negativity | True |
| Sparse regularizer $\lambda$ (network) | 0 | Normalization | True |
| Sparse regularizer $\lambda$ (loss) | 0 | Numbers | 3 |
| Code support knowledge | True | Length | 600 ms (24) |
| Code Q regularization | False | Smoother | False |
| Code Q regularization matrix | - | Smoother penalty | - |
| Code Q regularization period | - | Initialization | Random Normal |
| Q regularization scale | - | Share among neurons | True |
| Q regularization norm type | - |  |  |
| Top k sparsity | - |  |  |
| Top k period | - |  |  |
| Adam optimizer |  | Other network parameters |  |
| Number of epochs | 15 | Model distribution | Binomial |
| Batch size | 32 | Time bin resolution | 25 ms |
| Learning rate | 0.01 | Unrolling non15lin | Shrinkage |
| Learning rate decay | False | Unrolling number | 100 |
| Learning rate decay step | - | Unrolling mode | FISTA |
| Adam eps | 0.001 | Unrolling alpha | 0.1 |
| Backpropagation type | Truncated |  |  |
| Truncated iterations | 10 |  |  |

In our analysis, we characterized the response of neurons to the stimuli by one kernel. For more re-fined characterization, one may choose to learn two kernels; prior work discussed that each whisker cycle could evoke two response types (one that encodes the caudal direction whisker movement and another that captures the rostral direction movement on the way back to the neutral whisker position) [90].

### Olfactory experiment

An experimental session consists of ≈ 250 trials. In each trial, a custom device delivered 50 ms odor pulses of the same peak concentration to the animal’s nose at a Poisson-distributed pulse rate between 0.5-4 pulse/s for 5 s. Neural activity in the animal’s anterior piriform cortex was recorded with a custom-built 32-channel tetrode drive at a 30 kHz sampling rate using the Open Ephys recording system [136]. The data are downsampled to 1 ms resolution for analysis. Single-unit spiking activities were isolated using Kilosort2 [137]. We isolated 5-40 single units in each recording session. At the end of each session, the entire bundle of tetrodes was lowered by 40 *μm* to obtain a new set of neurons for the subsequent session. We recorded *C* = 770 neurons during *S* = 17 behavioural sessions from 3 mice. The data and model parameters for this experiment are summarized in Table S5.

The clustering analysis is based on K-means. Figure S11 shows K-means with 3, 4, 5, 6 clusters.

**Table S3.** Parameters for dopamine calcium experiment. For code non-negativity, -1,1,2 are for negative, positive, and two-sided, respectively. For kernel non-negativity flag, 0 is for negative/positive, and 1 is for positive.

| Data |  |  |  |
| --- | --- | --- | --- |
| Sampling rate | 15 Hz | Trial length | 4.53 - 15.2 s |
| Number of neurons | {6, 20, 30} | Number of trials | {252, 299, 195} |
| Total number of neurons | 56 | Total number of examples | 13,342 |
| Code |  | Kernel |  |
| Non-negativity | [2,2,1,1,-1] | Non-negativity | [0,0,0,1,1] |
| Sparse regularizer $\lambda$ (network) | 0 | Normalization | True |
| Sparse regularizer $\lambda$ (loss) | 0 | Numbers | 5 |
| Support knowledge | True | Length | 4 s (60) |
| Q regularization | True | Smoother | False |
| Q regularization matrix | see text | Smoother penalty | - |
| Q regularization period | 1 | Initialization | Random Normal |
| Q regularization scale | 2.5 | Share among neurons | True |
| Q regularization norm type | 2 |  |  |
| Top k sparsity | - |  |  |
| Top k period | - |  |  |
| Adam optimizer |  | Other network parameters |  |
| Number of epochs | 15 | Model distribution | Gaussian |
| Batch size | 8 | Time bin resolution | 1 ms |
| Learning rate | 0.01 | Unrolling nonlin | Shrinkage |
| Learning rate decay | False | Unrolling number | 100 |
| Learning rate decay step | - | Unrolling mode | FISTA |
| Adam eps | 0.001 | Unrolling alpha | 0.1 |
| Backpropagation type | Truncated |  |  |
| Truncated iterations | 10 |  |  |

We have used 90% of the neurons at random (repeated 40 times) to compute the adjusted random index (ARI); on average, ARI is 0.94, 0.96, 0.95, 0.77, for 3, 4, 5, 6 clusters, respectively. We observed similar clustering effect using spectral clustering with a Radial Basis Function (RBF) kernel.

### Dopamine axon multi-fiber photometry in the striatum

The training and modeling parameters of the multi-fiber photometry experiment are summarized in Table S6. The original data contains simultaneously recorded fluorescence traces from 14 fibers and a control patch in a single session of one freely behaving mouse (resulting in 15 independent runs of DUNL, though with the same parameters).

In our analysis, we performed event detection and kernel characterized with a single kernel per fiber, to find a particular feature of the signals that has been reported in the literature (kernel width [95]). For a more refined characterization, one may choose to learn more kernels. This is the subject of current work by a subset of the authors of this manuscript.

### Simulated unstructured spiking experiment with overlapping events

Simulation study I: This section summarizes the information on the experiment demonstrating the ability of DUNL to detect and locally characterize events appearing at random. In this experiment, there are two types of events, each happening three times in a trial. While there is a 200 ms minimum distance between events of the same type, events of different types are allowed to fully overlap. Table S7 summarizes the parameters of this experiment.

**Table S4.** Parameters for whisker thalamus experiment.

| Data |  |  |  |
| --- | --- | --- | --- |
| Sampling rate | 1 ms | Trial length | 4000 |
| Number of neurons | 10 | Number of Trials | (25 train, 25 test) |
| Total number of neurons | 10 | Total number of examples | (250 train, 250 test) |
| Code |  | Kernel |  |
| Non-negativity | True | Non-negativity | False |
| Sparse regularizer $\lambda$ (network) | 0.03 | Normalization | True |
| Sparse regularizer $\lambda$ (loss) | 0.03 | Numbers | 1 |
| Support knowledge | False | Length | 125 ms (25) |
| Q regularization | False | Smoother | True |
| Q regularization matrix | - | Smoother penalty | 0.003 |
| Q regularization period | - | Initialization | Stimuli velocity |
| Q regularization scale | - | Share among neurons | True |
| Q regularization norm type | - |  |  |
| Top k sparsity | 18 |  |  |
| Top k period | 10 |  |  |
| Adam optimizer |  | Other network parameters |  |
| Number of epochs | 120 | Model distribution | Binomial |
| Batch size | 30 | Time bin resolution | 5 ms |
| Learning rate | 0.01 | Unrolling nonlin | Shrinkage |
| Learning rate decay | False | Unrolling number | 800 |
| Learning rate decay step | - | Unrolling mode | FISTA |
| Adam eps | 0.001 | Unrolling alpha | 0.5 |
| Backpropagation type | Truncated |  |  |
| Truncated iterations | 0 |  |  |

### LFADS experiment

We used the source code https://github.com/tensorflow/models/tree/master/research/lfads to run experiments for LFADS. We used the default parameters while we modified FACTORS_DIM to match the number of kernels in the experiment.

### Simulated model characterization experiment

Simulation study II: Table S8 summarized all the modeling and training parameters for simulation on DUNL characterization. For all spiking data simulations, we use convolutional sparse coding explained in Methods to model the mean firing rates of the neurons; then, the firing rates are used for each time point to sample from a Bernoulli distribution.

### Simulated model characterization with five orthogonal kernels

Table S9 summarizes the parameters of the DUNL model characterization experiment with five orthogonal kernels. We note that the number of kernels and their length are overestimated for the noisy experiments, i.e., DUNL learns 6 kernels of length 84. Moreover, for noisy experiments, the noise, with a spiking probability ranging from 0.05 to 0.8, is sampled from a Binomial distribution for each time bin window. The noise is then either added or subtracted from the ground-truth spike counts. Finally, the noisy data is made sure to be within 0 to 5, the maximum possible number of spikes in each bin.

**Table S5.** Parameters for olfaction experiment.

| Data |  |  |  |
| --- | --- | --- | --- |
| Sampling rate | reduced to 1 ms | Trial length | 4500 ms |
| Number of neurons | 770 | Number of trials | ≈ 250 |
| Total number of neurons | - | Total number of examples | - |
| Code |  | Kernel |  |
| Non-negativity | True | Non-negativity | False |
| Sparse regularizer $\lambda$ (network) | varies | Normalization | True |
| Sparse regularizer $\lambda$ (loss) | 0 | Numbers | 1 |
| Support knowledge | True | Length | 1000 ms (20) |
| Q regularization | False | Smoother | True |
| Q regularization matrix | - | Smoother penalty | 0.1 |
| Q regularization period | - | Initialization | Aligned raster |
| Q regularization scale | - | Share among neurons | False |
| Q regularization norm type | - |  |  |
| Top k sparsity | - |  |  |
| Top k period | - |  |  |
| Adam optimizer |  | Other network parameters |  |
| Number of epochs | 500 | Model distribution | Poisson |
| Batch size | Full-batch | Time bin resolution | 50 ms |
| Learning rate | 0.01 | Unrolling nonlin | Shrinkage |
| Learning rate decay | False | Unrolling number | 100 |
| Learning rate decay step | - | Unrolling mode | FISTA |
| Adam eps | 0.001 | Unrolling alpha | 0.1 |
| Backpropagation type | Full |  |  |
| Truncated iterations | - |  |  |

We used this dataset to compare DUNL to SCA [33]. The parameters used for SCA are summarized in Table S10.

### Simulated structured spiking experiment with non-overlapping events

This section summarizes the information on the data used for the comparison of DUNL and LFADS in their ability to capture local characteristics from single trials. Table S11 summarizes the parameters of this experiment.

**Table S6.**
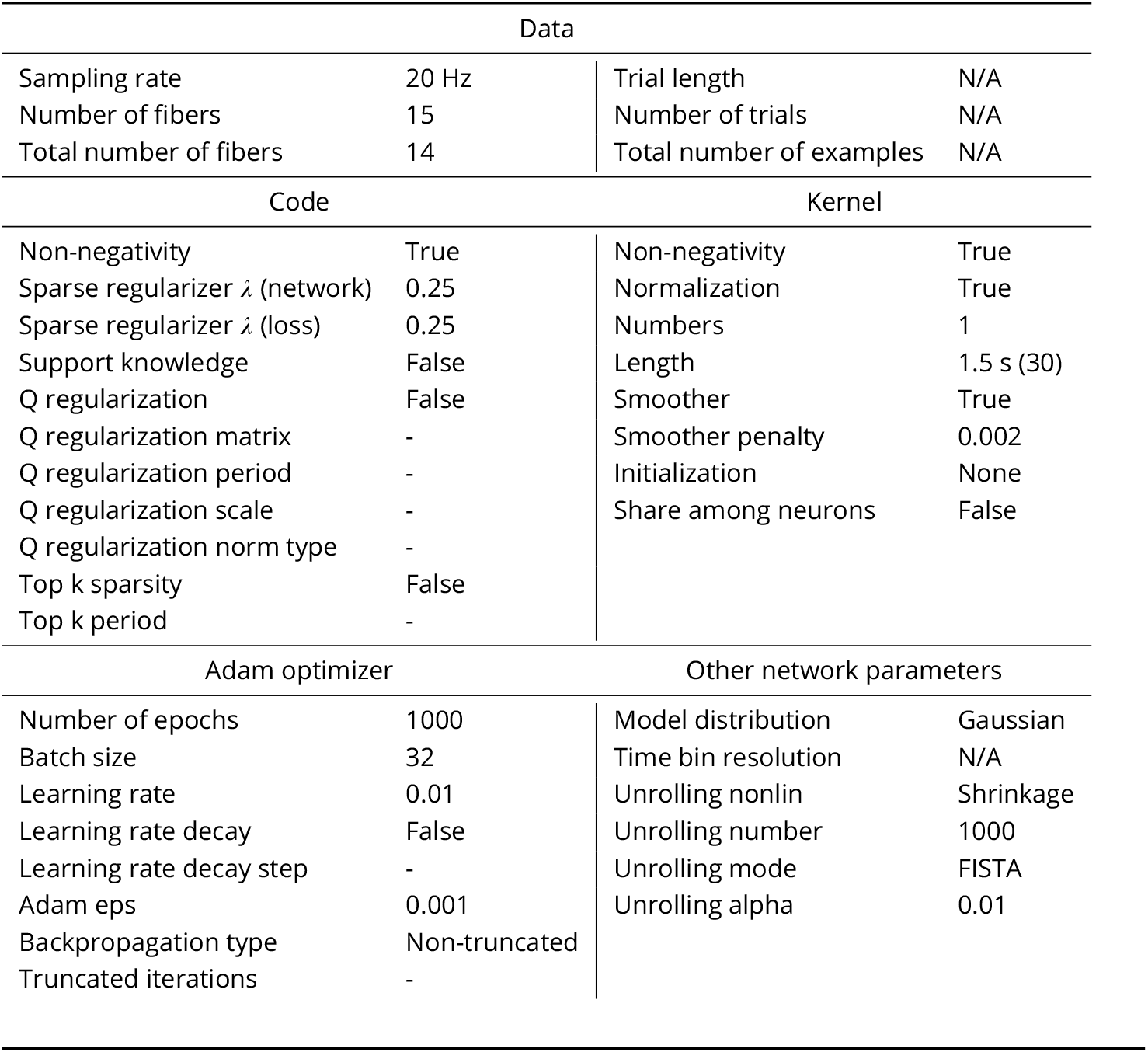
Parameters for multi-fiber photometry experiment.

| Data |  |  |  |
| --- | --- | --- | --- |
| Sampling rate | 20 Hz | Trial length | N/A |
| Number of fibers | 15 | Number of trials | N/A |
| Total number of fibers | 14 | Total number of examples | N/A |
| Code |  | Kernel |  |
| Non-negativity | True | Non-negativity | True |
| Sparse regularizer $\lambda$ (network) | 0.25 | Normalization | True |
| Sparse regularizer $\lambda$ (loss) | 0.25 | Numbers | 1 |
| Support knowledge | False | Length | 1.5 s (30) |
| Q regularization | False | Smoother | True |
| Q regularization matrix | - | Smoother penalty | 0.002 |
| Q regularization period | - | Initialization | None |
| Q regularization scale | - | Share among neurons | False |
| Q regularization norm type | - |  |  |
| Top k sparsity | False |  |  |
| Top k period | - |  |  |
| Adam optimizer |  | Other network parameters |  |
| Number of epochs | 1000 | Model distribution | Gaussian |
| Batch size | 32 | Time bin resolution | N/A |
| Learning rate | 0.01 | Unrolling nonlin | Shrinkage |
| Learning rate decay | False | Unrolling number | 1000 |
| Learning rate decay step | - | Unrolling mode | FISTA |
| Adam eps | 0.001 | Unrolling alpha | 0.01 |
| Backpropagation type | Non-truncated |  |  |
| Truncated iterations | - |  |  |

### Simulated unstructured spiking experiment with overlapping events

This section summarizes the information on the data used for the comparison of DUNL and SeqNMF in their ability to capture local characteristics from single trials. Table S12 summarizes the parameters of DUNL. We note that there are two underlying kernel in this dataset, where DUNL overestimates and learn 3 kernels from the dataset. For SeqNMF, we used the github source code https://github.com/FeeLab/seqNMF, and key parameters are summarized in Table S13.

### Simulated dopamine spiking experiment

The dopamine spiking simulation follows closely the data from the dopamine spiking real experiment. Table S14 summarizes the parameters of this experiment.

## Supplementary Materials - Data Acquisition

### Two-photon calcium imaging

#### Surgeries

Stereotaxic viral injections and GRIN lens implantation: Surgeries were performed under aseptic conditions. Mice were anesthetized with isoflurane (1–2 at 0.5–1 L.min^−1^), and local anesthetic (lidocaine (2%)/bupivacaine (0.5%) 1:1 mixture, subcutaneous (s.c.)) was applied at the incision site. Analgesia (buprenorphine for pre-operative treatment, 0.1 mg.kg^−1^, intraperitoneal (i.p.); ketoprofen for post-operative treatment, 5 mg.kg^−1^, i.p.) was administered for 3 days after surgery. A custom-made head plate was placed on the well-cleaned and dried skull with adhesive cement (C&B Metabond, Parkell) containing a small amount of charcoal powder. To express the calcium indicator GCaMP in dopamine neurons, AAV5-CAG-FLEX-GCaMP7f (1.8 × 10^13^ particles per ml) was injected unilaterally in the VTA (300 nl, bregma - 3.0 mm AP, 0.5 mm ML, 4.6 mm DV from dura) in two DAT-Cre mice (*Slc6a3*^*tm*1.1(*cre*)*Bkmn*^, Jackson Laboratory, 006660) [138] respectively. A third mouse was a double transgenic resulting from crossing DAT-Cre with Ai148D (B6.Cg-Igs7^*tm*148.1(*tetO*−*GCaMP* 6*f, CAG*−*tT A*2)*Hze*/*J*^, Jackson Laboratory, 030328) [139] for expression of GCaMP6f in dopamine neurons. The injection was done at a rate of approximately 20 nl.min^−1^ for a total of 300 nl using a manual plunger injector (Narishige). For both DAT-Cre and DAT-Cre;Ai148 double transgenic mice, a GRIN lens (0.6 mm in diameter, 7.3 mm length; 1050-004597, Inscopix) was slowly inserted above the VTA after insertion and removal of a 25-gauge needle. The implants were secured with C&B Metabond adhesive cement (Parkell) and dental acrylic (Lang Dental).

**Table S7.** Parameters for simulated unstructured spiking experiment for detection and deconvolution of two event types.

| Data |  |  |  |
| --- | --- | --- | --- |
| Sampling rate | 1 ms | Trial length | 6000 ms |
| Number of neurons | 1 | Number of Trials | 25 - 1600 |
| Total number of neurons | 1 | Total number of examples | 25 - 1600 |
| Code |  | Kernel |  |
| Non-negativity | True | Non-negativity | False |
| Sparse regularizer $\lambda$ (network) | 0.1 | Normalization | True |
| Sparse regularizer $\lambda$ (loss) | 0.1 | Numbers | 2 |
| Support knowledge | False | Length | 400 ms (16) |
| Q regularization | False | Smoother | True |
| Q regularization matrix | - | Smoother penalty | 0.015 |
| Q regularization period | - | Initialization | Random Normal |
| Q regularization scale | - | Share among neurons | True |
| Q regularization norm type | - |  |  |
| Top k sparsity | 3 |  |  |
| Top k period | 10 |  |  |
| Adam optimizer |  | Other network parameters |  |
| Number of epochs | 200-2000 | Model distribution | Binomial |
| Batch size | 128 | Time bin resolution | 25 ms |
| Learning rate | 0.01 | Unrolling nonlin | Shrinkage |
| Learning rate decay | False | Unrolling number | 800 |
| Learning rate decay step | - | Unrolling mode | FISTA |
| Adam eps | 0.001 | Unrolling alpha | 0.25 |
| Backpropagation type | Truncated |  |  |
| Truncated iterations | 5 |  |  |

#### Behavioral training and testing protocol

Mice were water-deprived in their home cage 1–2 days before the start of behavioral training, two or more weeks after surgery. During water deprivation, each mouse’s weight was maintained above 85% of its original value. Mice were habituated to the head-fixed setup by receiving water every 4 s (6 *μ*l drops) for 3 days, after which association between odors and outcomes started. A mouse lickometer (1020, Sanworks) was used to measure licking as infrared beam breaks. Water valves (LHDA1233115H, The Lee Company) were calibrated, and a custom-made olfactometer based on a valve driver module (1015, Sanworks) and a valve mount manifold (LFMX0510528B and LHDA1221111H valves, The Lee Company) was used for odor delivery. All components were controlled through a Bpod state machine (1027, Sanworks). Odors were diluted in mineral oil (Sigma-Aldrich) at 1:10, and 30 *μ*l of each diluted odor was placed inside a syringe filter (2.7-*μ*m pore size, 6823-1327, GE Healthcare). Odorized air was further diluted at 1:10 and delivered at 1,000 ml.min^−1^. Odors used for each association were randomly assigned from the following list of odors: isoamyl acetate, p-cymene, ethyl butyrate, (+)-carvone, (±)-citronellal, α-ionone, L-fenchone. One of these odors was associated with a distribution of reward sizes while a second odor was not paired with any outcome (nothing). For the rewarded odor, after a 2-s trace period, a reward was delivered whose size was taken randomly from a uniform distribution of the following sizes: 0.3, 0.5, 1.2, 2.5, 5.0, 8.0, 11.0 *μ*l. Variable-size non-cued rewards taken from the same distribution were also delivered throughout the sessions. Mice completed one session per day.

**Table S8.** Parameters for simulated model characterization experiment. The information in the parenthesis are for different time bin resolution scenario of (5 ms, 10 ms, 25 ms, 50 ms).

| Data |  |  |  |
| --- | --- | --- | --- |
| Sampling rate | 1 ms | Trial length | 4000 |
| Number of neurons | 1 | Number of trials | 25, 50, 100, 250, 500 |
| Total number of neurons | 1 | Total number of examples | 25, 50, 100, 250, 500 |
| Code |  | Kernel |  |
| Non-negativity | True | Non-negativity | False |
| Sparse regularizer $\lambda$ (network) | 0.03 | Normalization | True |
| Sparse regularizer $\lambda$ (loss) | 0.03 | Numbers | 1 |
| Support knowledge | - | Length | 500 ms (100, 50, 20, 10) |
| Q regularization | False | Smoother | True |
| Q regularization matrix | - | Smoother penalty | (0.2, 0.01, 0.004, 0.0002) |
| Q regularization period | - | Initialization | Sinusoidal shape |
| Q regularization scale | - | Share among neurons | True |
| Q regularization norm type | - |  |  |
| Top k sparsity | 5 |  |  |
| Top k period | 10 |  |  |
| Adam optimizer |  | Other network parameters |  |
| Number of epochs | 15-100 | Model distribution | Binomial |
| Batch size | 128 | Time bin resolution | (5, 10, 25, 50) ms |
| Learning rate | 0.01 | Unrolling nonlin | Shrinkage |
| Learning rate decay | False | Unrolling number | 800 |
| Learning rate decay step | - | Unrolling mode | FISTA |
| Adam eps | 0.001 | Unrolling alpha | 0.25 |
| Backpropagation type | Truncated |  |  |
| Truncated iterations | 20 |  |  |

#### Image acquisition

Imaging was performed using a custom-built two-photon microscope. The microscope was equipped with a diode-pumped, mode-locked Ti:sapphire laser (Mai-Tai, Spectra-Physics). All imaging was done with the laser tuned to 920 nm. Scanning was achieved using a galvanometer and an 8-kHz resonant scanning mirror (adapted confocal microscopy head, Thorlabs). Laser power was controlled using a Pockels Cell (ConOptics 305 with M302RM driver). The average beam power used for imaging was 40–120 mW at the tip of the objective (Plan Fluorite ×20, 0.5 NA, Nikon). Fluorescence photons were reflected using two dichroic beamsplitters (FF757-Di01-55×60 and FF568-Di01-55×73, Semrock), were filtered using a bandpass filter (FF01-525/50-50, Semrock), and were collected using GaAsP photomultiplier tubes (H7422PA-40, Hamamatsu), whose signal was amplified using transimpedance amplifiers (TIA60, Thorlabs). Microscope control and image acquisition were done using ScanImage 4.0 (Vidrio Technologies). Frames with 512×512 pixels were acquired at 15 Hz. Synchronization between behavioral and imaging acquisitions were achieved by triggering microscope acquisition in each trial to minimize photobleaching using a mechanical shutter (SC10, Thorlabs).

#### Data pre-processing

Acquired images were pre-processed in the following manner. (1) Movement correction was performed using phase correlation image registration implemented in Suite2P[140]. (2) Region-of-interest (ROI) selection was performed manually in FIJI from the mean and standard deviation projections of a subset of frames from the entire acquisition, as well as a movie of the frames used to build those projections. (3) Neuropil decontamination was performed with FISSA[141] using four regions around each ROI. The neuropil decontaminated fluorescent signal was then filtered with a 12 point Gaussian kernel with 0.6875 standard deviation. Drift along the session was corrected using the running maximum of the running minimum of a 120 s time window. Then Δ*F* /*F*_0_ was calculated as 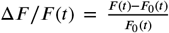 using as *F*_0_ the 6^*th*^ running percentile in a window of 40 s. This fluorescent trace was used for further data processing and analysis in the network.

**Table S9.** Parameters for model characterization with five orthogonal kernels.

| Data |  |  |  |
| --- | --- | --- | --- |
| Sampling rate | 1 ms | Trial length | 6000 ms |
| Number of neurons | 1000 | Number of trials | 20 - 1000 |
| Total number of neurons | 1000 | Total number of examples | 20 - 1000 |
| Code |  | Kernel |  |
| Non-negativity | True | Non-negativity | False |
| Sparse regularizer $\lambda$ (network) | 0.005 | Normalization | True |
| Sparse regularizer $\lambda$ (loss) | 0.005 | Numbers | 5 |
| Support knowledge | False | Length | 400 ms (80) |
| Q regularization | False | Smoother | True |
| Q regularization matrix | - | Smoother penalty | 0.01 |
| Q regularization period | - | Initialization | Random Normal |
| Q regularization scale | - | Share among neurons | True |
| Q regularization norm type | - |  |  |
| Top k sparsity | 1 |  |  |
| Top k period | 10 |  |  |
| Group sparse regularizer | 0.05 |  |  |
| Adam optimizer |  | Other network parameters |  |
| Number of steps | 1000 | Model distribution | Binomial |
| Batch size | 5 | Time bin resolution | 5 ms |
| Learning rate | 0.01 | Unrolling nonlin | Shrinkage |
| Learning rate decay | False | Unrolling number | 20 |
| Learning rate decay step | - | Unrolling mode | FISTA |
| Adam eps | 0.001 | Unrolling alpha | 0.25 |
| Backpropagation type | Truncated |  |  |
| Truncated iterations | 5 |  |  |

**Table S10.** Parameters of SCA in experiment with five orthogonal kernels.

|  |  |
| --- | --- |
| Number of components | 5 |
| Number of epochs | 3000 |
| Learning rate | 0.001 |
| Initialization of weights | PCA |
| Sparsity penalty | 0.1 |
| Orthogonality bool | True |
| Orthogonality penalty | 0.1 |

**Table S11.** Parameters for simulated structured spiking experiment for comparison of DUNL with LFADS for their ability to learn local characterization from data.

| Data |  |  |  |
| --- | --- | --- | --- |
| Sampling rate | 1 ms | Trial length | 2000 ms |
| Number of neurons | 1 | Number of trials | 25 - 1600 |
| Total number of neurons | 1 | Total number of examples | 25 - 1600 |
| Code |  | Kernel |  |
| Non-negativity | True | Non-negativity | False |
| Sparse regularizer $\lambda$ (network) | 0.1 | Normalization | True |
| Sparse regularizer $\lambda$ (loss) | 0.1 | Numbers | 2 |
| Support knowledge | False | Length | 400 ms (16) |
| Q regularization | False | Smoother | True |
| Q regularization matrix | - | Smoother penalty | 0.015 |
| Q regularization period | - | Initialization | Random Normal |
| Q regularization scale | - | Share among neurons | True |
| Q regularization norm type | - |  |  |
| Top k sparsity | 1 |  |  |
| Top k period | 10 |  |  |
| Adam optimizer |  | Other network parameters |  |
| Number of epochs | 125-1500 | Model distribution | Binomial |
| Batch size | 128 | Time bin resolution | 25 ms |
| Learning rate | 0.01 | Unrolling nonlin | Shrinkage |
| Learning rate decay | False | Unrolling number | 800 |
| Learning rate decay step | - | Unrolling mode | FISTA |
| Adam eps | 0.001 | Unrolling alpha | 0.25 |
| Backpropagation type | Truncated |  |  |
| Truncated iterations | 5 |  |  |

### Multi-fiber photometry

#### Surgeries

Optical fiber implantation: Surgeries were performed under aseptic conditions as described above for two-photon imaging. One DAT-Cre::Ai148::tdTomato triple transgenic mouse, which expresses GCaMP6s and tdTomato in dopamine neurons, was implanted with a custom-made head plate and custom 14-optical fiber implant inserted above the striatum. The implants were secured with C&B Metabond adhesive cement (Parkell) and dental acrylic (Lang Dental).

#### Behavioral testing protocol

After performing other tasks not reported in this manuscript, which are the subject of separate investigations, the mouse was put inside a box to interact freely with multiple stimuli introduced in the box by the experimenter.

#### Fluorescence acquisition and pre-processing

To collect fluorescence signal from dopamine axons across the striatum we used a bundle-imaging fiber photometry system with two integrated CMOS cameras and 3 integrated light-emitting diodes (LEDs) (BFMC6_LED(410-420)_LED(460-490)_CAM(500-550_LED(555-570)_CAM(580-680), Doric Lenses).

The three LEDs were triggered sequentially. Image acquisition was collected in the cameras so that camera 1 collected the fluorescence emitted by GCaMP when excitated by LED 1 (410-420 nm, isosbestic wavelength) and 2 (460-490 nm, calcium-dependent modulation), while camera 2 collected fluorescence from tdTomato (excited by LED 3(555-570 nm)). Each wavelength was acquired at 20Hz in a 640×640 pixel image. The objective of the BFMC6 was focused on the top FC connector of an optical fiber bundle. The cameras imaged the top of this fiber bundle and regions-of-interest (ROI) corresponding to each optical fiber were defined in the Doric Neuroscience Studio software (Doric Lenses). The average pixel count was used as a measure of fluorescence emitted through each fiber. The bundled fiber patchcord was connected to the fiber receptacle implanted in the mouse brain to deliver excitation light to the brain and collect the fluorescence emission signals from it simultaneously. The green and red fluorescence signals from the brain were spectrally separated from each other and from the excitation lights using dichroic mirrors integrated in the BFMC6. LED driving currents were chosen so that each excitation wavelength provided around 100 *μ*W at the tip of each optical fiber in the patchcord bundle.

**Table S12.** Parameters of DUNL for the experiment comparison with seqNMF.

| Data |  |  |  |
| --- | --- | --- | --- |
| Sampling rate | 1 ms | Trial length | 6000 ms |
| Number of neurons | 500 | Number of trials | 50 - 2000 |
| Total number of neurons | 500 | Total number of examples | 50 - 2000 |
| Code |  | Kernel |  |
| Non-negativity | True | Non-negativity | False |
| Sparse regularizer $\lambda$ (network) | 0.005 | Normalization | True |
| Sparse regularizer $\lambda$ (loss) | 0.005 | Numbers | 2 |
| Support knowledge | False | Length | 400 ms (80) |
| Q regularization | False | Smoother | True |
| Q regularization matrix | - | Smoother penalty | 0.01 |
| Q regularization period | - | Initialization | Random Normal |
| Q regularization scale | - | Share among neurons | True |
| Q regularization norm type | - |  |  |
| Top k sparsity | 1 |  |  |
| Top k period | 10 |  |  |
| Group sparse regularizer | 0.025 |  |  |
| Adam optimizer |  | Other network parameters |  |
| Number of steps | 1000 | Model distribution | Binomial |
| Batch size | 10 | Time bin resolution | 5 ms |
| Learning rate | 0.01 | Unrolling nonlin | Shrinkage |
| Learning rate decay | False | Unrolling number | 20 |
| Learning rate decay step | - | Unrolling mode | FISTA |
| Adam eps | 0.001 | Unrolling alpha | 0.25 |
| Backpropagation type | Truncated |  |  |
| Truncated iterations | 5 |  |  |

**Table S13.** Parameters of SeqNMF for the experiment comparison.

|  |  |
| --- | --- |
| K | 3 |
| L | 80 |
| lambda | 0.01 |
| lambdaL1H | any value ranging from 0.1 to 100 won't result in desirable performance |
| lambdaOrthoH | 1 |
| lambdaL1W | 0 |
| lambdaOrthoW | 0 |

**Table S14.** Parameters for simulated dopamine spiking experiment.

| Data |  |  |  |
| --- | --- | --- | --- |
| Sampling rate | 1 ms | Trial length | 3100 ms |
| Number of neurons | 40 | Number of trials | [14 - 300] |
| Total number of neurons | 40 | Total number of examples | [560 - 1200] |
| Code |  | Kernel |  |
| Non-negativity | False | Non-negativity | True |
| Sparse regularizer $\lambda$ (network) | 0 | Normalization | True |
| Sparse regularizer $\lambda$ (loss) | 0 | Numbers | 3 |
| Support knowledge | True | Length | 600 ms (24) |
| Q regularization | False | Smoother | False |
| Q regularization matrix | - | Smoother penalty | - |
| Q regularization period | - | Initialization | Random Normal |
| Q regularization scale | - | Share among neurons | True |
| Q regularization norm type | - |  |  |
| Top k sparsity | - |  |  |
| Top k period | - |  |  |
| Adam optimizer |  | Other network parameters |  |
| Number of epochs | 200 | Model distribution | Binomial |
| Batch size | 2 | Time bin resolution | 25 ms |
| Learning rate | 0.01 | Unrolling nonlin | Shrinkage |
| Learning rate decay | False | Unrolling number | 100 |
| Learning rate decay step | - | Unrolling mode | FISTA |
| Adam eps | 0.001 | Unrolling alpha | 0.1 |
| Backpropagation type | Truncated |  |  |
| Truncated iterations | 10 |  |  |

The fluorescence of each ROI (14 optical fibers + 1 control patch) was preprocessed by calculating Δ*F* /*F*_0_ as 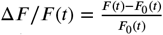 using as *F*_0_ the 6^*th*^ running percentile in a window of 20 s. Possible movement artifacts were corrected by subtracting from the GCaMP signal a prediction of movement signal calculated using linear regression from either the isosbestic or the tdTomato signal, whichever had the most significant correlation coefficient to the original GCaMP signal. This Δ*F* /*F*_0_ signal was used as input to DUNL’s network.

### Supplementary Materials - Figures

**Figure S2.**
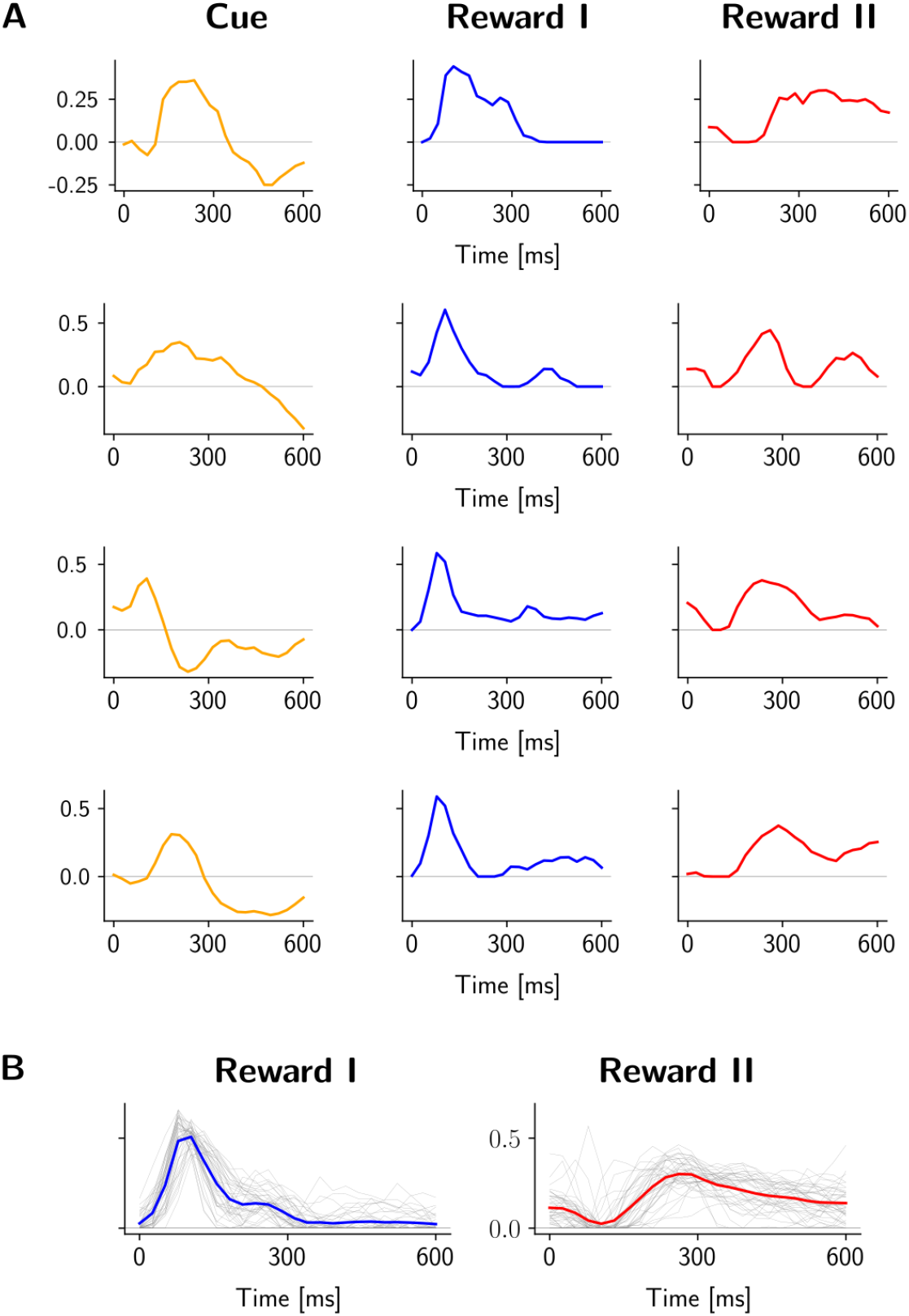
Result of training DUNL while inferring the baseline activity, and finding kernels for individual dopamine neurons (contrary to Figure 2, Figure S3, and Figure S4, in which kernels were shared across neurons). (A) Learned kernels for 4 example neurons. (B)Kernel for all neurons (light gray lines), and population average (blue and red lines for Reward I (salience-like) and Reward II (value-like) respectively).

**Figure S3.**
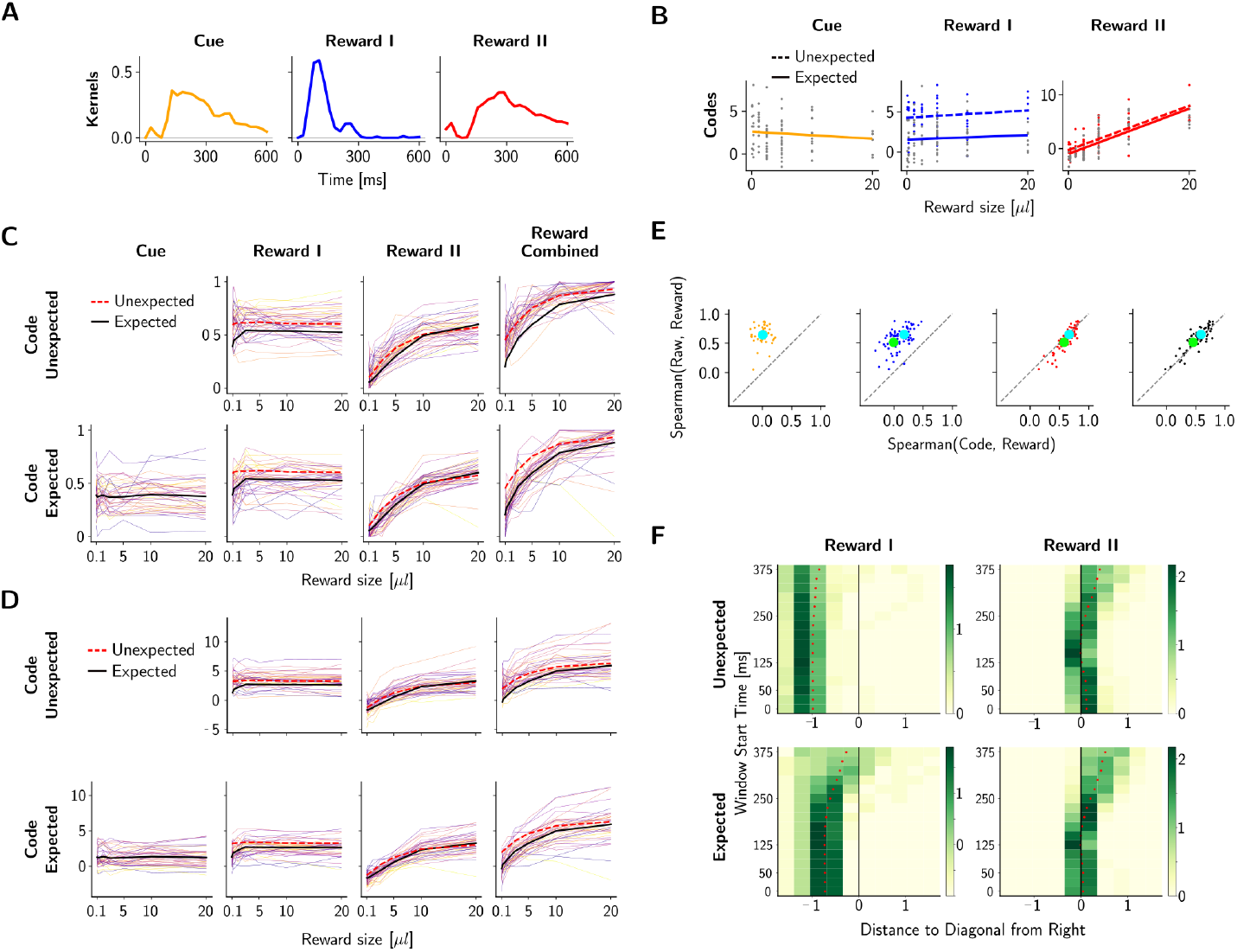
Result of training DUNL without inferring the baseline activity in the unrolling process (instead, we estimate from the pre-odor period). (A)Learned kernels shared across neurons. (B), Codes in all trials as a function of reward size for an example neuron. (C)Mean neural code amplitudes (across trials) as a function of reward size for all neurons (each line is a neuron). (D)Same as (C) without normalization. (E)Spearman’s rank correlation between codes and reward size (x-axis) vs. the windowed average firing rates and reward sizes (y-axis). (F)Histogram of distance of dots from the diagonal in Spearman’s rank correlation in (D); positive distance means below the diagonal and colorbar shows the normalized probability density function at each bin, such that the integral over the shown range in x-axis is 1.

**Figure S4.**
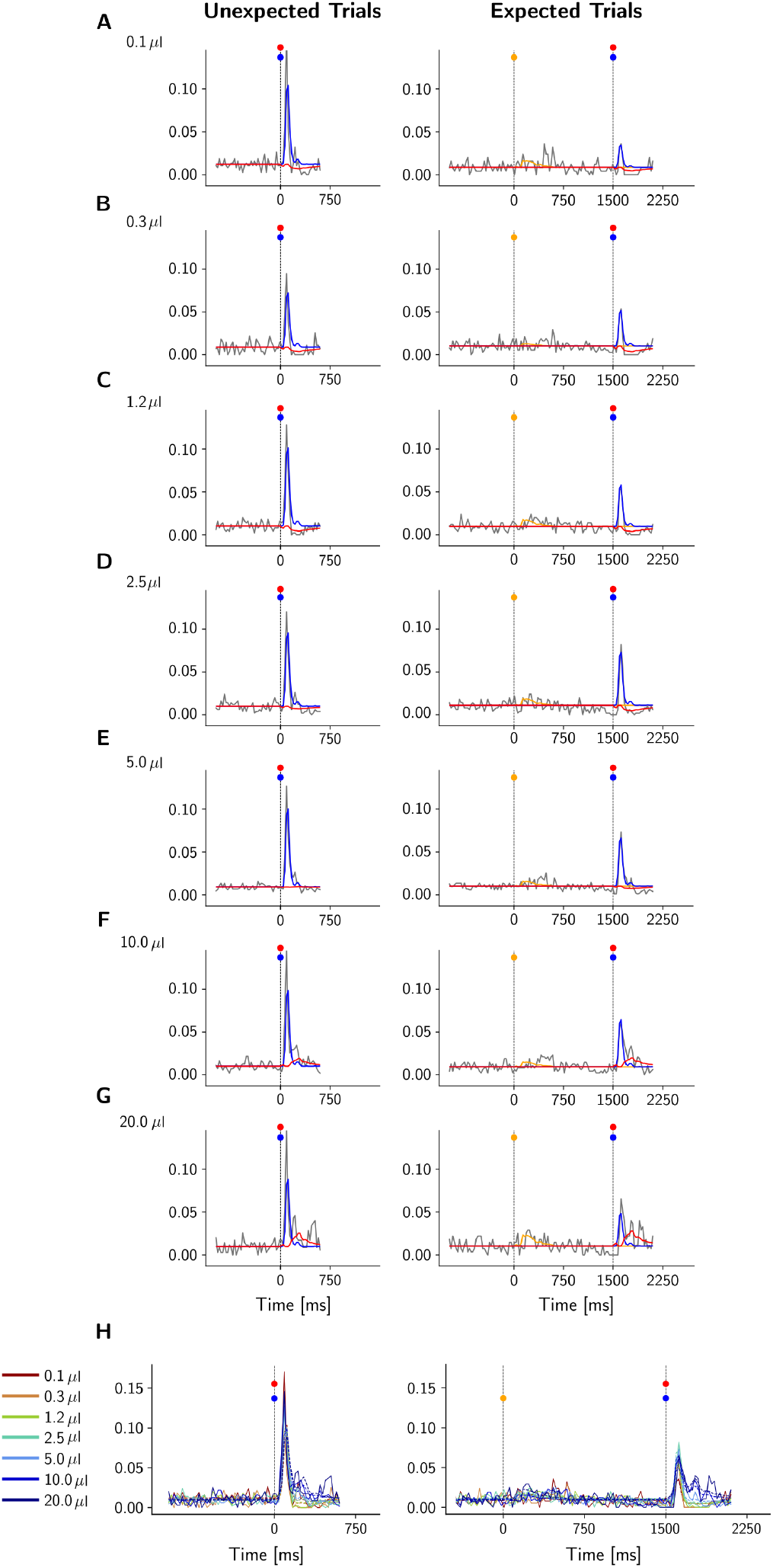
Decomposition of a single dopamine neuron spiking activity using shared kernels and individual codes (Same model as in Figure S3). (A-G) Averaged trial activity for each reward size for unexpected (left) and expected (right) trials (gray traces), were decomposed into Reward I (salience-like, blue) and Reward II (value-like, red) components. The salience kernel contributes in rate estimation of the burst right after the reward onset, and the value kernel contributes in representation of the spikes appearing with around a 100 ms delay. The dip in the neural activity for low reward amount is captured by a negative value code and highlights a negative RPE. (H) Summary of mean neural response for each trial type and reward size (solid lines) and the corresponding DUNL-reconstructed activity trace (dashed lines).

**Figure S5.**
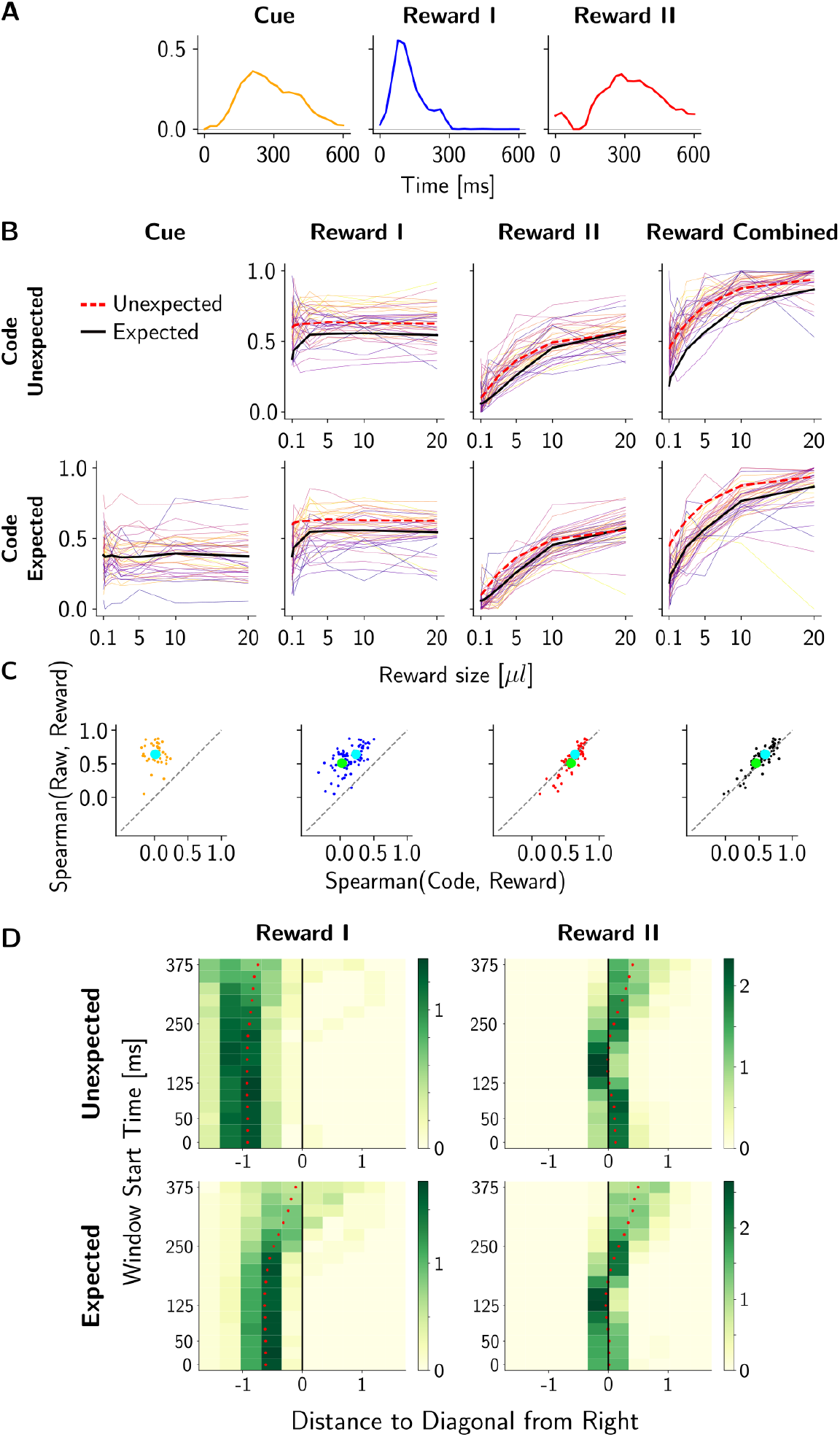
Result of training DUNL with a limited number of trials from dopamine spiking data [87]. (A) Learned kernels shared across neurons. (B)Neural code amplitudes as a function of reward size; the figure demonstrates diversity of neural encodings with each line corresponding to one neuron. (C)Spearman’s rank correlation between codes and reward size (x-axis) vs. the windowed average firing rates and reward sizes (y-axis). (D)Histogram of distance of dots from the diagonal in Spearman’s rank correlation from (C); positive distance means below the diagonal and colorbar shows the normalized probability density function at each bin, such that the integral over the shown range in x-axis is 1.

**Figure S6.**
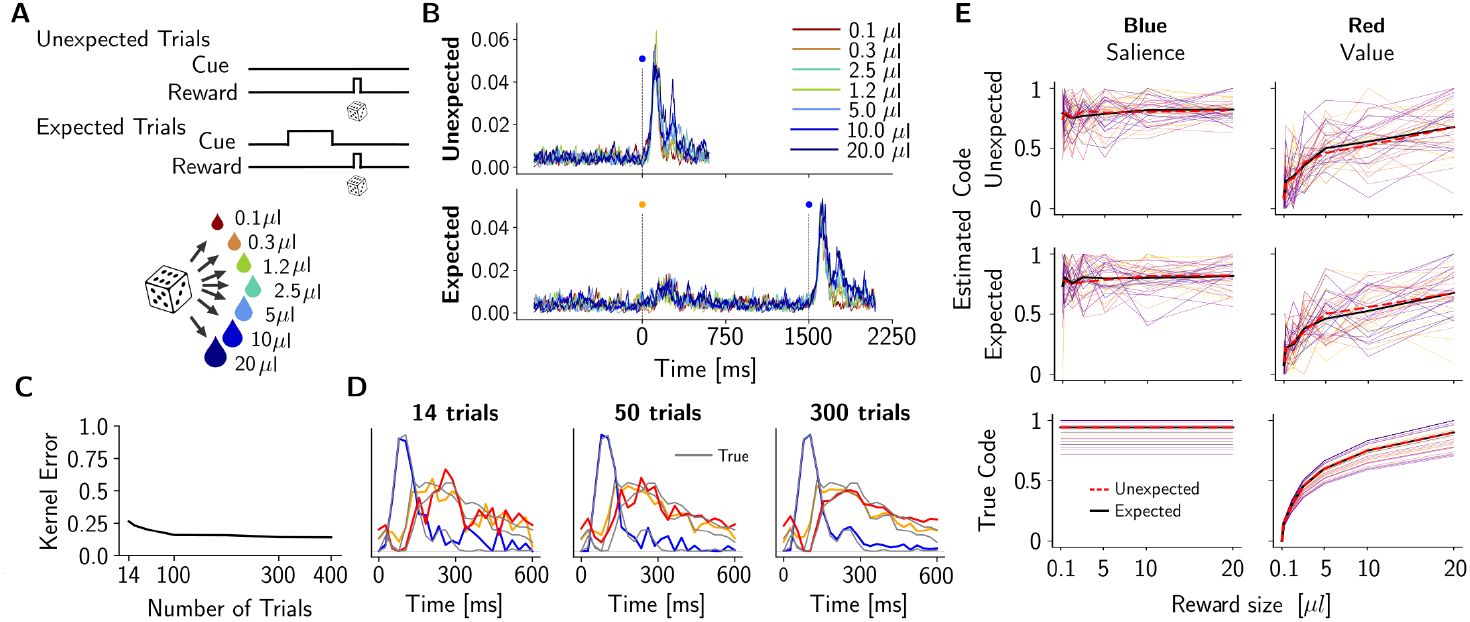
Analysis of kernel quality with the number of trials in simulated dopamine data. (A)The experiment setup used to generate the data. (B)PSTH of simulated neurons over each trial type. (C)The kernel recovery error (i.e., 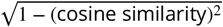. (D)Visualization of the learned kernels in color (the true underlying kernels are shown in gray). (E)DUNL’s code estimates as a function of reward sizes (top), and the true underlying code (bottom).

**Figure S7.**
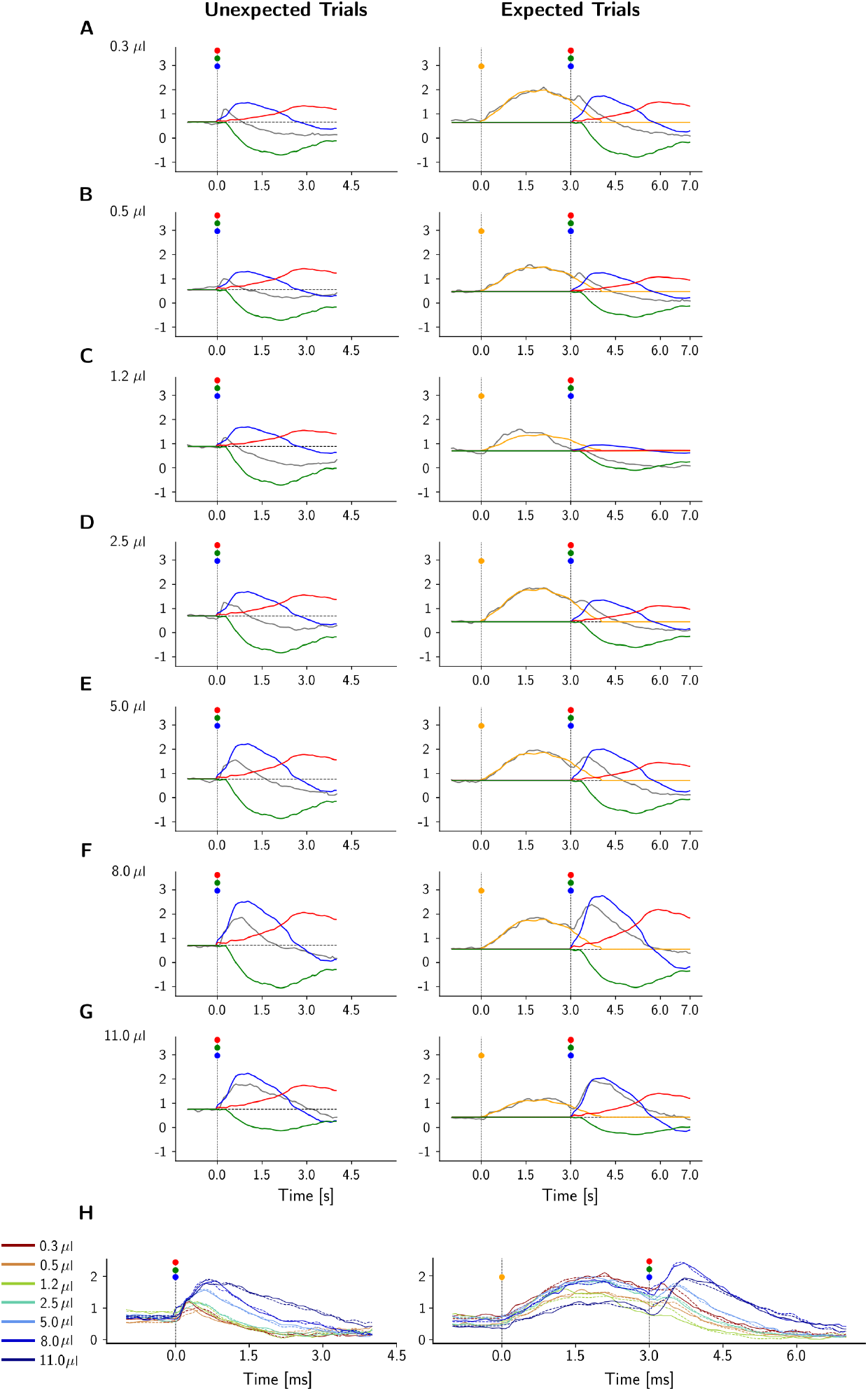
DUNL decomposition of responses from one dopamine neuron recorded using two-photon calcium imaging across reward sizes. (A-G) in unexpected (left) and expected (right) trials. The trial average raw data (gray) and its reconstruction in 4 kernels. Blue models the salience response, red models a positive response for value, and green represents a negative activity for value. (H) Summary of mean neural response for each trial type and reward size (solid lines) and the corresponding DUNL-reconstructed activity trace (dashed lines).

**Figure S8.**
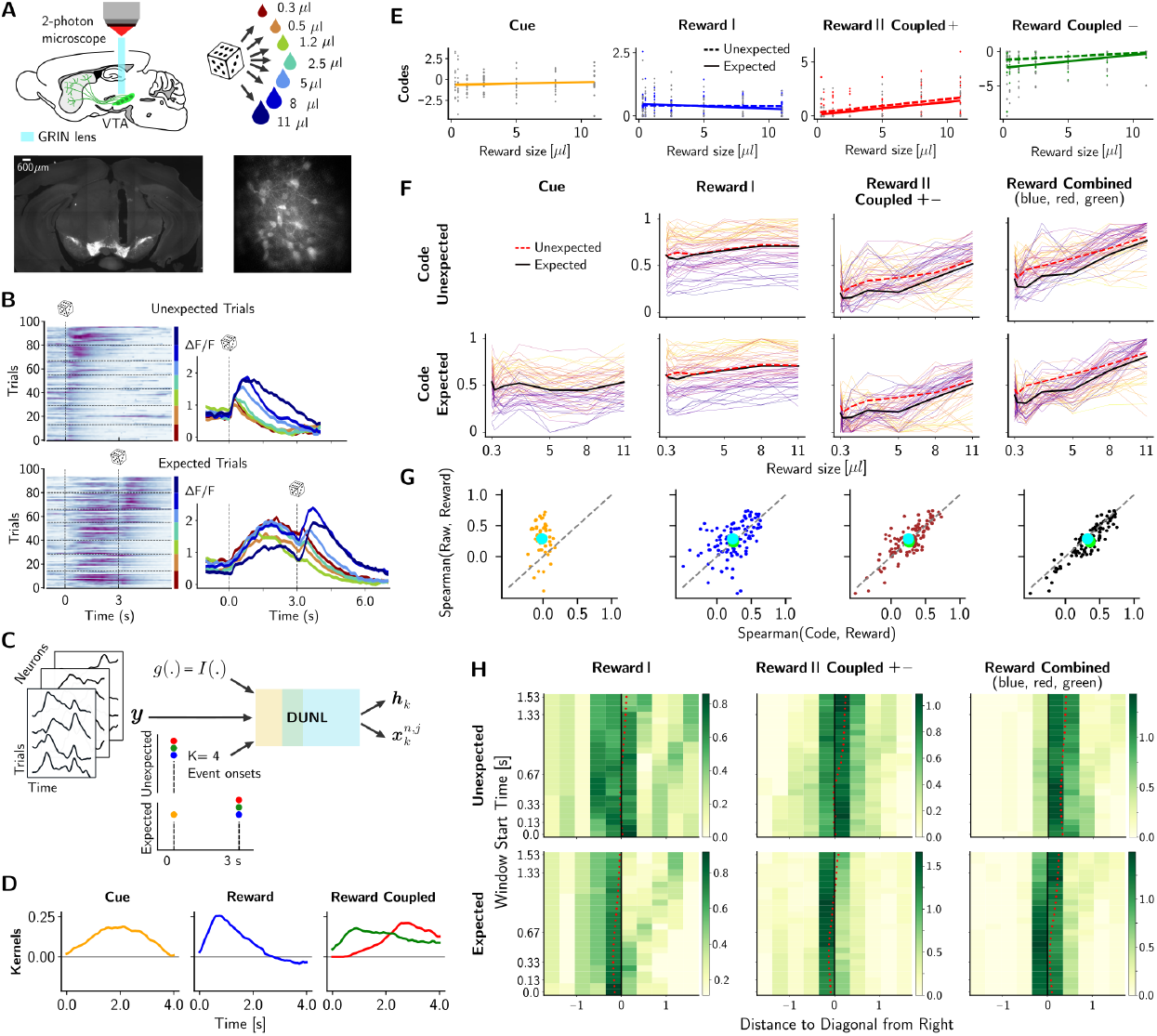
Result of training DUNL on the calcium dopamine data without inferring the baseline activity in the unrolling process (instead, we estimate from the pre-odor period). (A) Top: experimental setup depicted on a sagittal slice of the mouse brain with dopamine neurons represented in green; botton: histology images showing dopamine neurons expressing the fluorescent calcium indicator GCaMP6m: coronal slice of the mouse brain showing GRIN lens track over the VTA (left, scalebar: 600*μ*m), and projection image of the field of view obtained during an experimental acquisition under the two-photon microscope showing individual neurons (right). (B) Left: Heatmap of time-aligned trials at the reward onset. The trials are ordered from low to high reward size, with a horizontal line separating the different trial types. Right: Averaged time-aligned activity of an example neuron for each reward size. (C) Inputs used to run DUNL in this dataset: calcium activity across time and trials, timing of stimuli, number of kernels to learn, and probabilistic generative model for continuous calcium data. (D) Kernel characterization of cue and reward events. Three kernels were used to estimate the reward response: one for salience (blue) and two non-concurrent kernels for positive or negative value (green and red). (E) Code amplitude as a function of the reward size for an example neuron (each dot corresponds to the code inferred from single-trial neural activity; these values are fitted by linear regression). (F) Diversity of neural encodings as a function of reward size for unexpected (top) and expected trials (bottom): each line represents one neuron, the black line shows the average for expected trials, and the red dashed line average for unexpected trials. Activity is normalized per neuron and across trial types, and codes for comparison across subfigures. (G) Spearman correlation of the codes (x-axis) and the windowed average activity of 4 seconds (y-axis) with respect to the reward sizes: each dot represents one neuron and the average across all neurons are shown by blue (expected) and green (unexpected) large marker (Reward Combined has *p* = 0.008, and *p* = 3.468 *∗* 10^−9^ t-test, respectively). The third panel (brown) from the left combines the code from Reward Coupled kernels (positive and negative, depending on the trial). The right panel combines all the reward-related codes (salience-like Reward and value-like Reward-Coupled). (H) Heatmap of the distance of the yellow (unexpected) and green (expected) ‘x’ marker in (F), from the diagonal as a measure of the increased Spearman’s correlation between codes and reward size, as the interval chosen for the ad-hoc window is modified: it shrinks from the bottom to the top of the y-axis to gradually exclude the early activities after the onset. Positive values are located below the diagonal. On the right panel, the marker is closest to the diagonal when 0.4 s of activity at the reward onset is excluded in the ad-hoc window approach. Colorbar: normalized probability density function at each bin, such that the integral over each line in the x-axis is 1.

**Figure S9.**
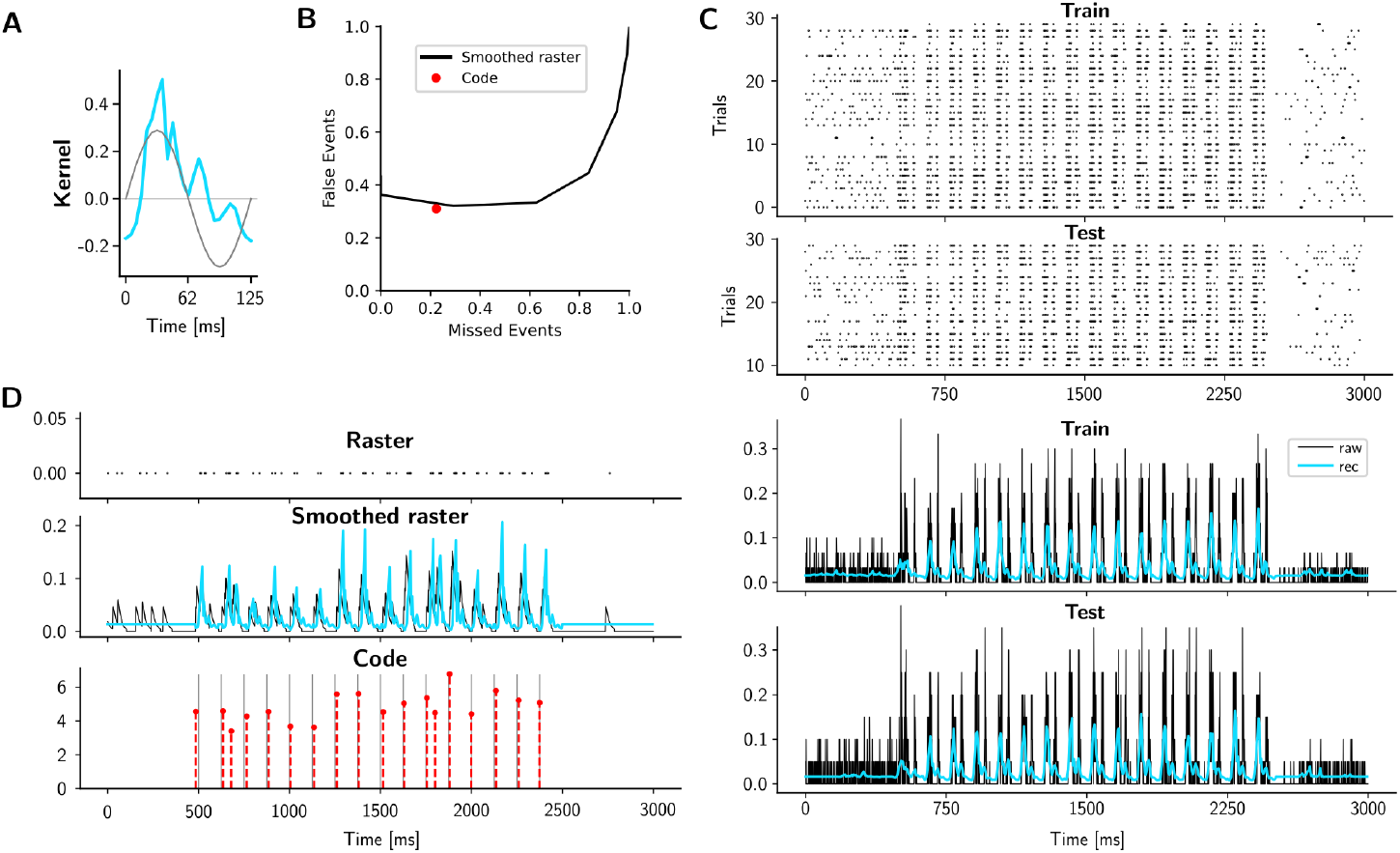
Event detection with DUNL (without group sparsity across neurons) for analysis of spiking data from the somatosensory thalamus. (A) Learned kernel for the whisker motion (blue). The gray sinusoid is the first derivative of stimulus motion. (B) Quantification of the miss/false events detected by DUNL. The red dot represents the performance of DUNL when events are detected on single-trials in the absence of group sparsity across neurons. The black curve shows the performance of a peak-finding algorithm on the smoothed spike rate in black. (C) Raster example from one neuron (top), and peristimulus time histogram from the corresponding neuron (black) along with the DUNL estimate of the firing rate (blue) (bottom). (D) Single trial spike raster (top), smoothed spike rates (black), and the rate estimation in blue (middle), with inferred code from the single trial for detecting 16 + 2 events in time (bottom).

**Figure S10.**
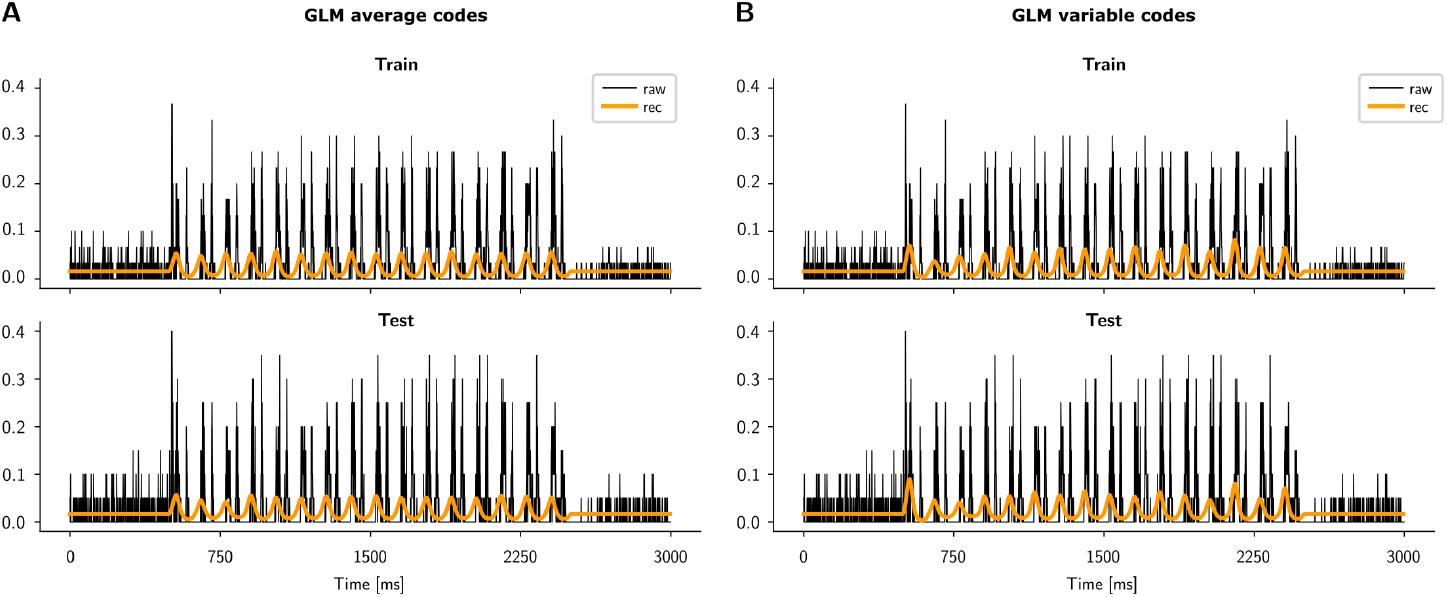
Analysis of spiking data from the somatosensory thalamus using two GLM-based baselines. (A) Using the GLM approach with fixed amplitude and known onsets to fit the data using the optimal kernel that is found and reported by [92]. The Peristimulus time histogram from one neuron (black) and the rate estimate (orange). The average ***R***^2^ across neurons is 0.1136. (B) Using the GLM approach with variable amplitude across events and known onsets to fit the data using the optimal kernel that is found and reported by [92]. The Peristimulus time histogram from the samme neuron as in (A) and the rate estimate (orange). The average ***R***^2^ across neurons is 0.1470.

**Figure S11.**
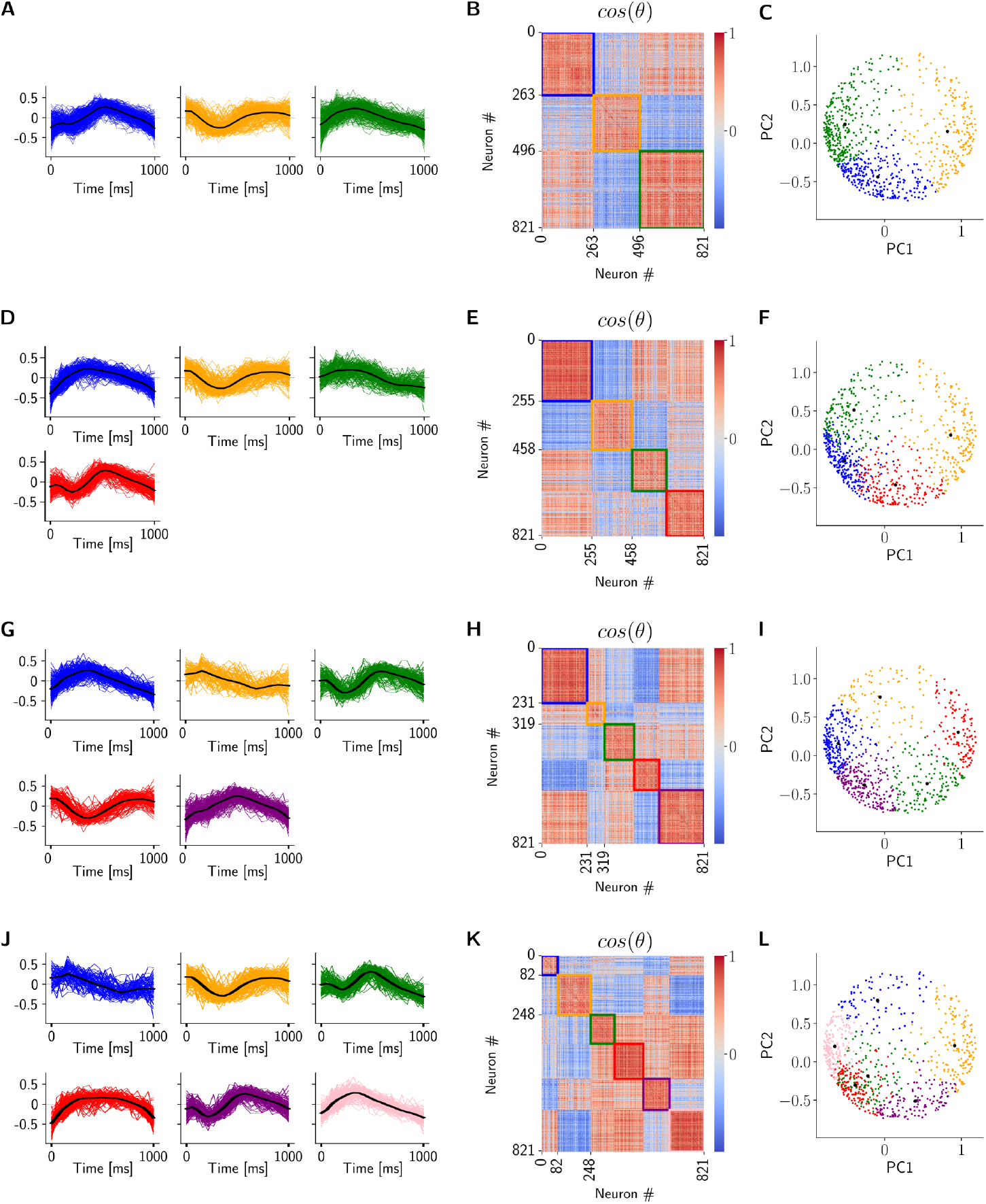
k-means clustering on the DUNL kernels obtained from the piriform cortex neural recordings. (A)Individual (colored traces) and mean (black trace) kernels when using 3 clusters. (B)Similarity matrix across kernels, using cosine distance. (C)Cluster visualizations on the first versus second principal components. (D-F) Same as (A-C) for 4 clusters. (G-I) Same as (A-C) for 5 clusters. (J-L) Same as (A-C) for 6 clusters. We observed similar clustering results when using spectral clustering.

**Figure S12.**
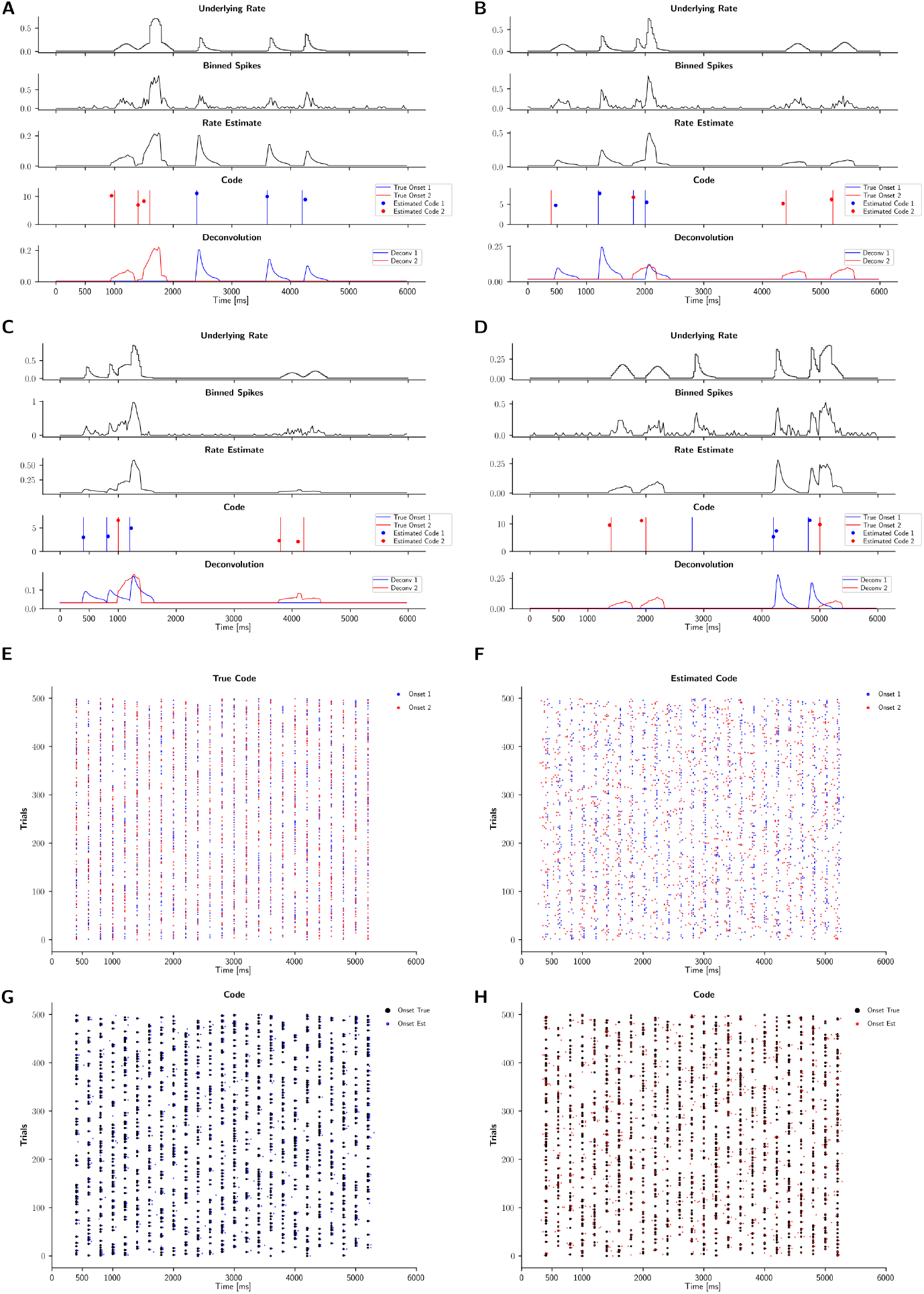
Model characterization with 2 kernels (blue and red). (A-D) Rate estimation and decomposition of 4 example trials. (E)The underlying true event onsets from both kernels across trials (blue and red). (F)The estimated code onsets by DUNL. (G)Same as E-F, but grouping true events and estimated codes in the same plot, for kernel/code 1 (blue). (H) Same as (G) for kernel/code 2 (red).

**Figure S13.**
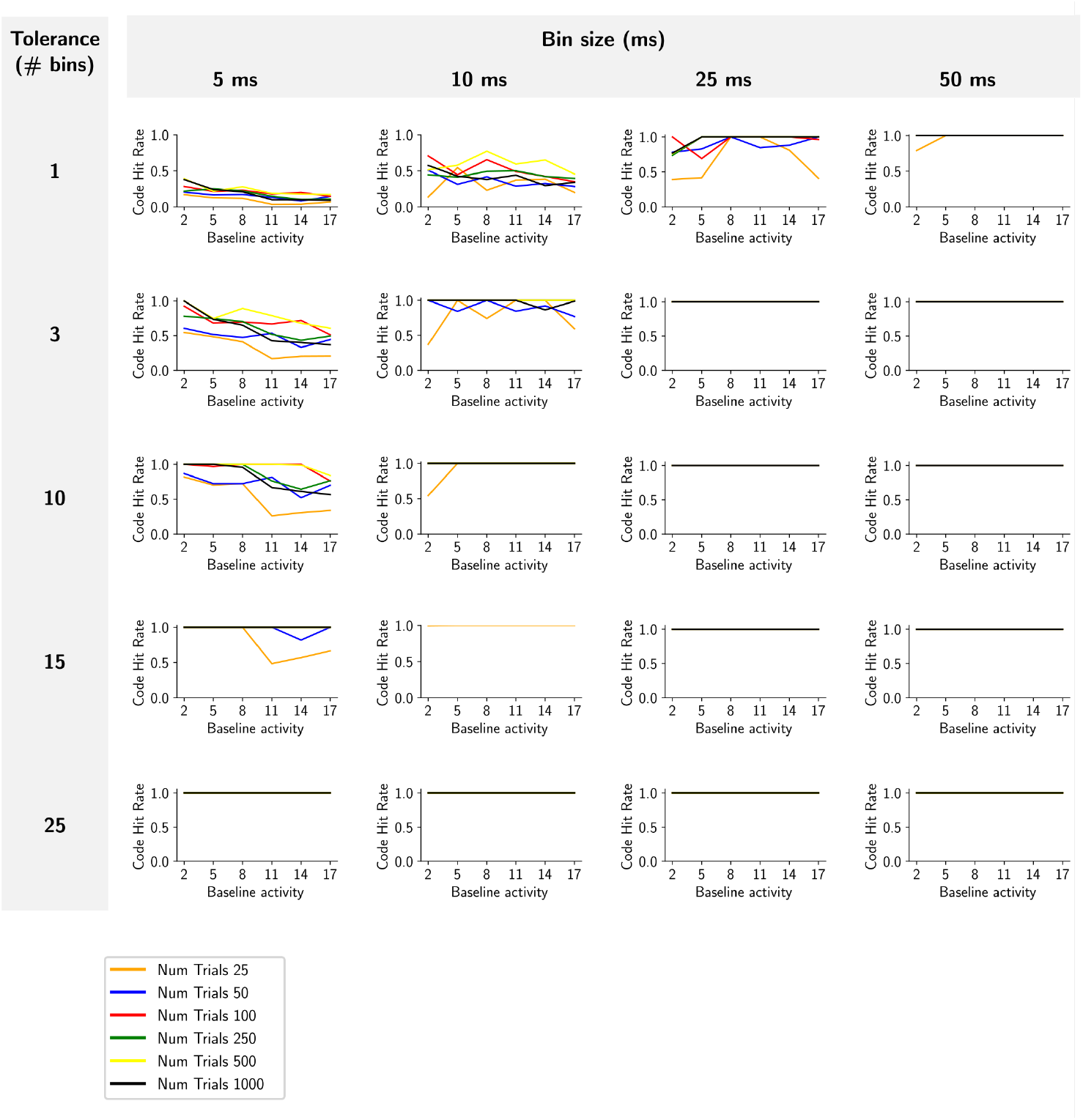
Model characterization with a single kernel. Code hit rate for event identification as a function of bin-size (columns) and time-tolerance (rows).

**Figure S14.**
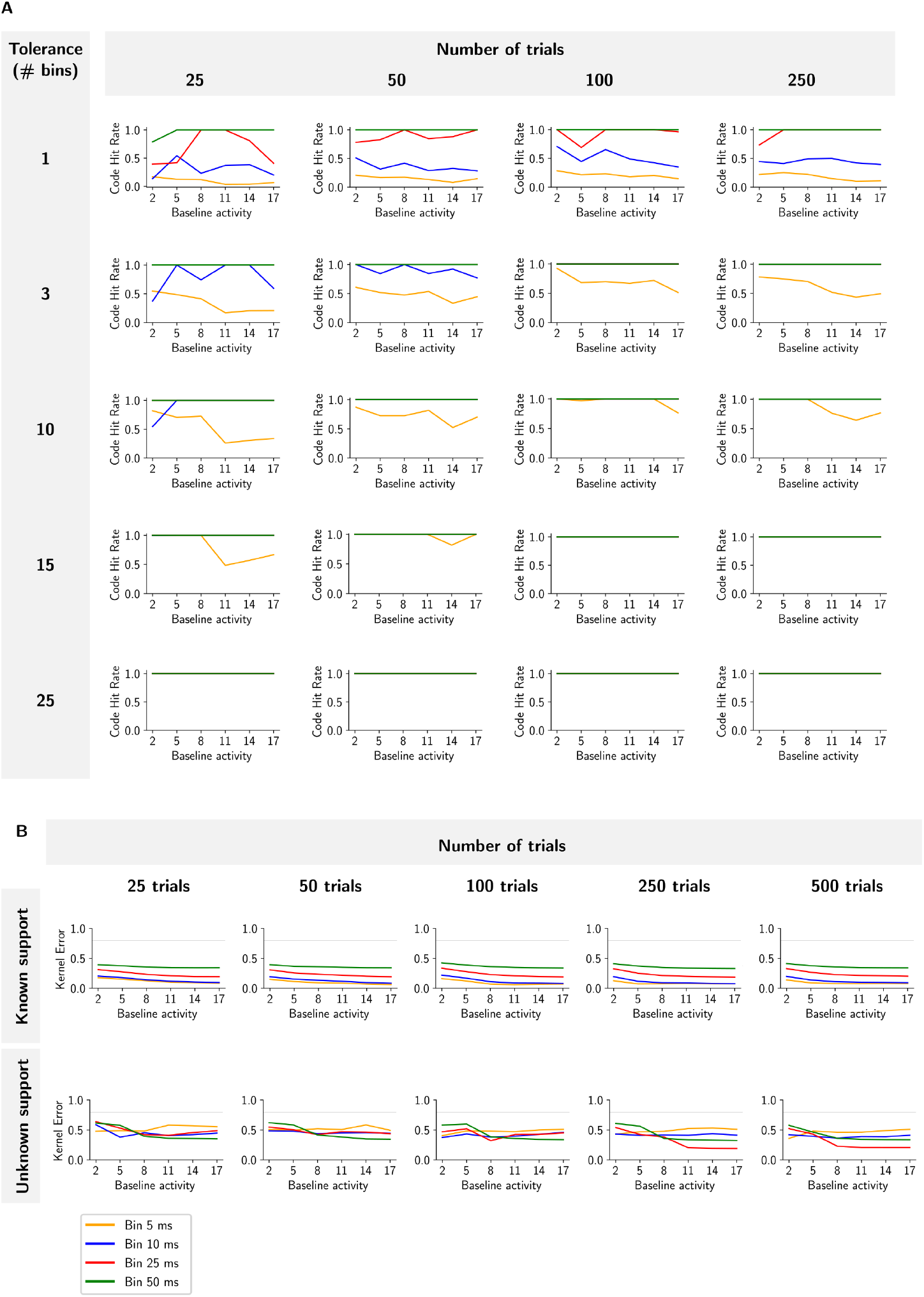
Characterization of DUNL’s performance as a function of the number of trials and time-tolerance. (A)Code hit rate for event identification as a function of the number of trials (columns) and time-tolerance (rows). (B)Kernel recovery error as a function of number of trials available for training for known support (top) and unknown support (bottom) scenarios.

**Figure S15.**
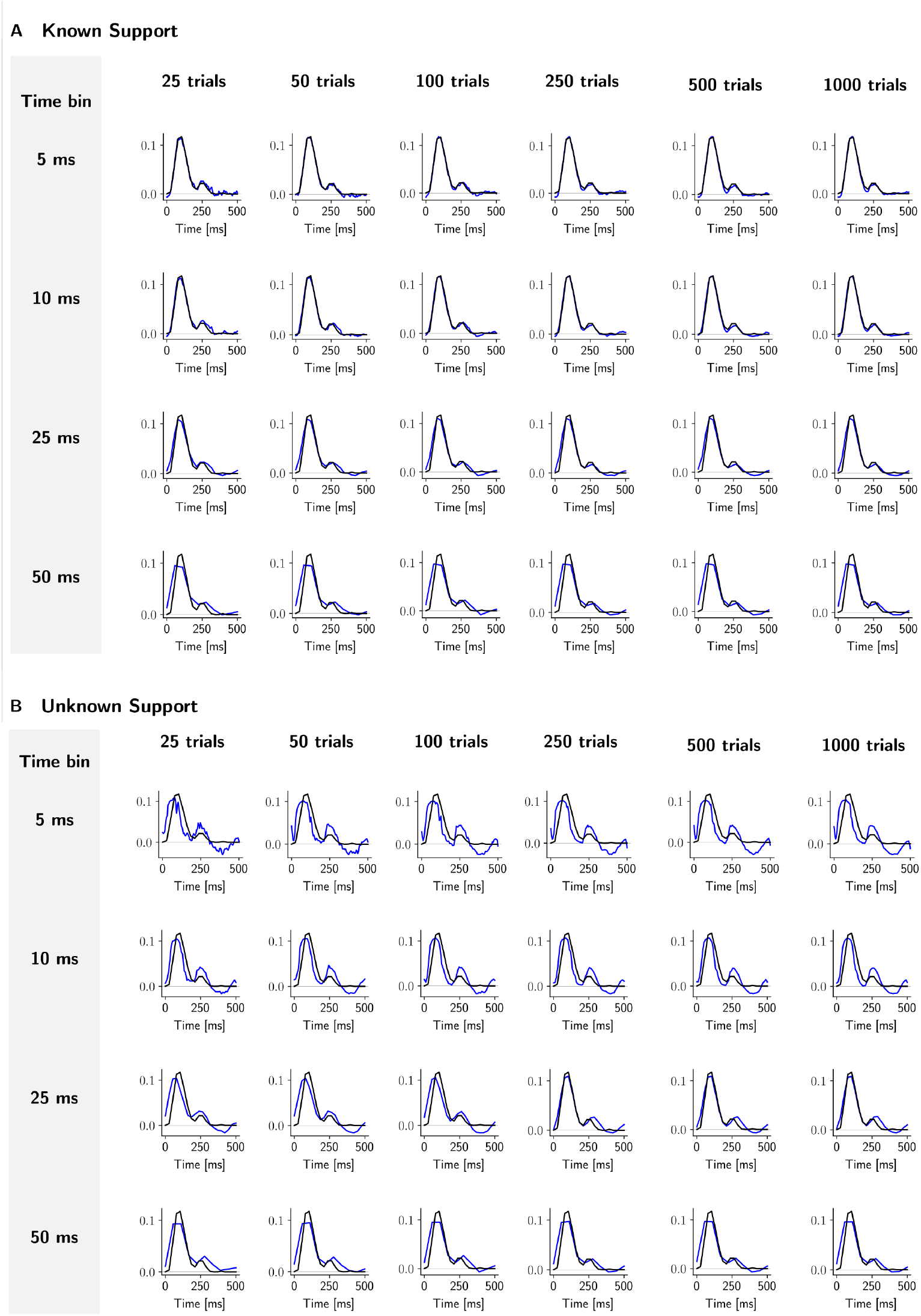
Kernel visualization as a function of trials (columns) and bin-size (rows). (A)Known event onsets (support). (B)Unknown event onsets.

**Figure S16.**
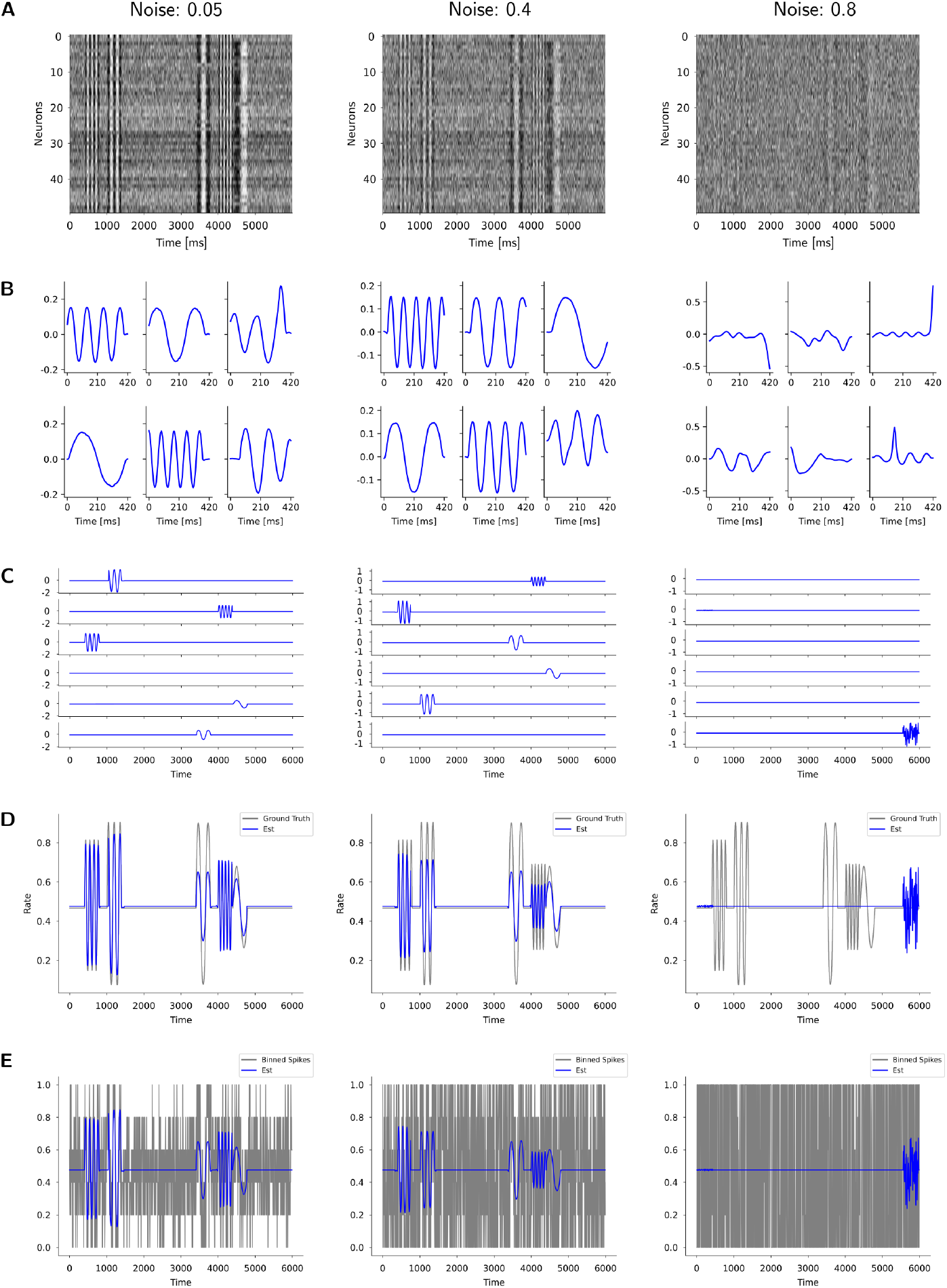
Robustness of kernel shape as a function of SNR. (A)Activity of 50 (out of 1000) synthetic neurons in response to 5 kernels that occur in random order and timing across trials. The same example trial is shown with different levels of additive noise: 0.05, 0.4, and 0.8 from left to right. (B)Kernels recovered by DUNL for each noise level. (C)Location of the kernels recovered by DUNL within the temporal frame of the example trial shown in (A). (D)Reconstruction of the population activity in blue, for the true underlying population activity in gray. (E)Example neuron with its true activity (gray) which is contaminated by noise, and the recovered signal in each noise condition (blue).

**Figure S17.**
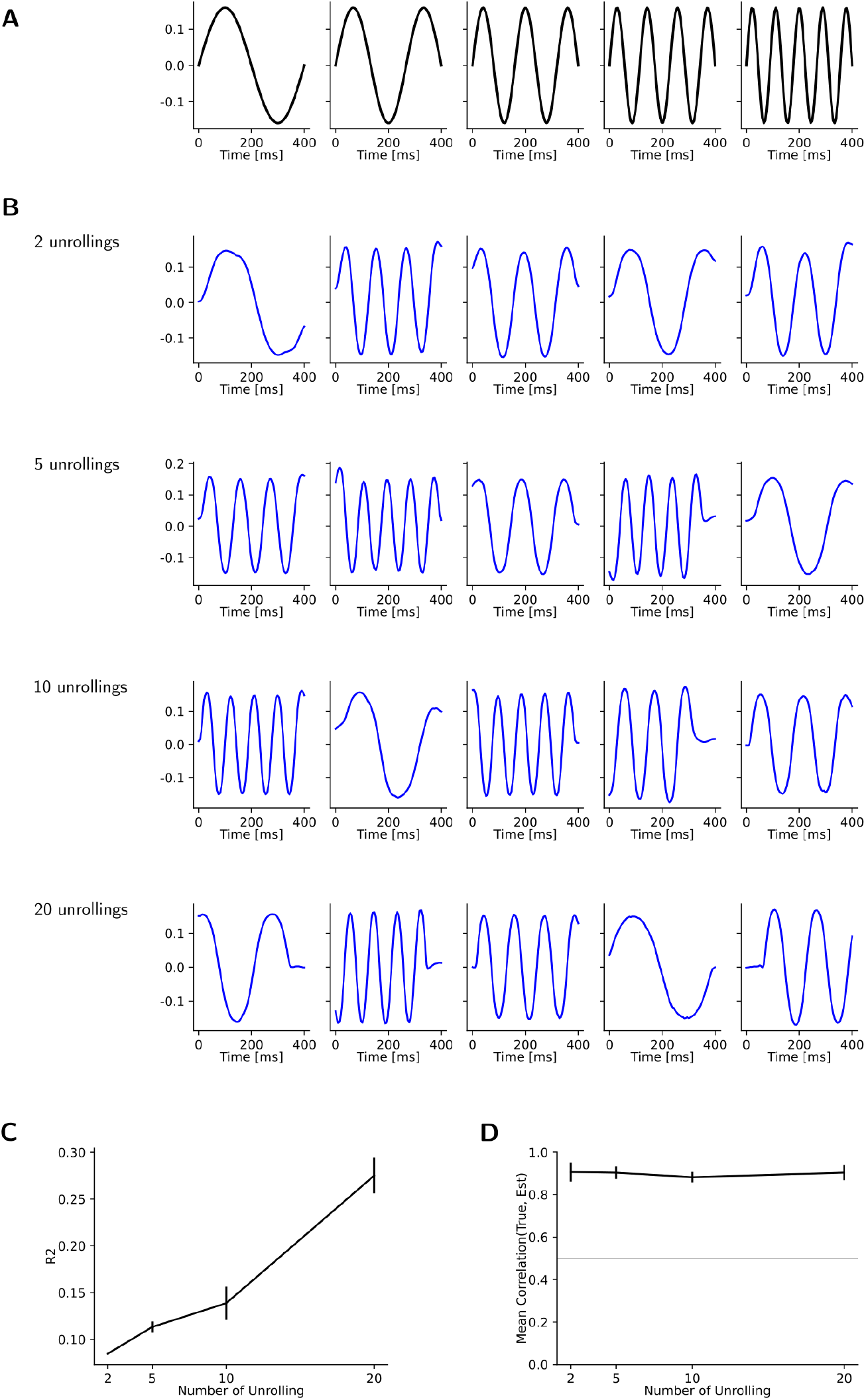
DUNL model fitting as a function of unrolling for the 5 orthogonal kernel simulation. (A) True kernels used for generating the synthetic spiking data. (B) Kernels obtained after 2, 5, 10, and 20 unrolling steps. (C) ***R***^2^ as a function of unrolling number. ***R***^2^ values are low because in this experiment the *l*1 distance is minimized for the code. Despite this, some of the kernels are recovered even after 2 unrolling steps only. Since these kernels often overlap in time during single trials, more unrolling steps result in a better estimate of the kernels. We recommend to use the maximum correlation between two kernels to select the right number of kernels to use.

**Figure S18.**
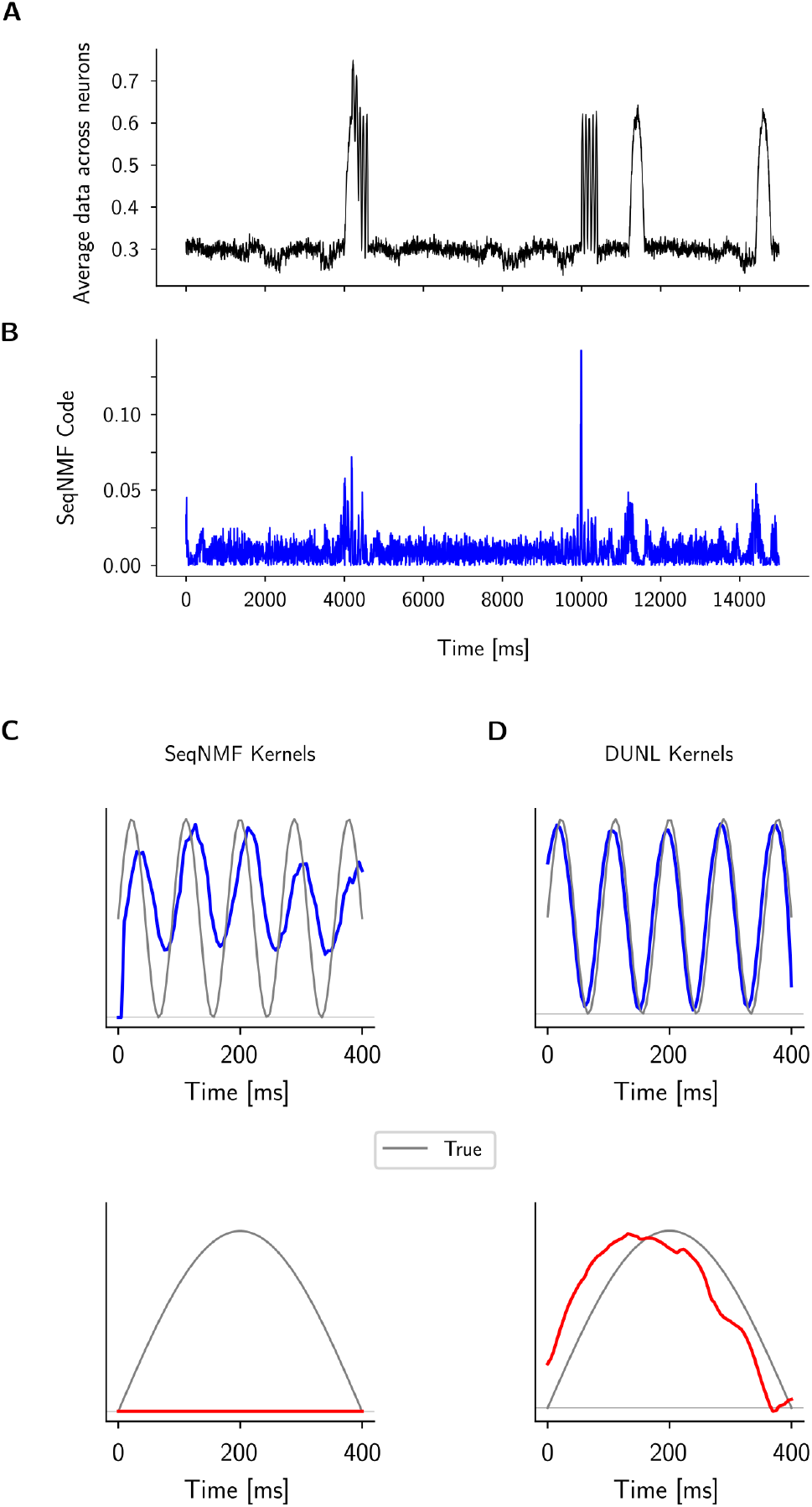
SeqNMF cannot recover the two local kernels expressed across the population of neurons. (A)Average activity of the synthetic neurons responding to two kernels that happen randomly during a trial. (B)SeqNMF code for the trial shown in (A). (C) True kernels (gray) and estimated kernels obtained with SeqNMF. The blue kernel is recovered, but not the red one. (D) DUNL recovers both the blue and the red kernels.

**Figure S19.**
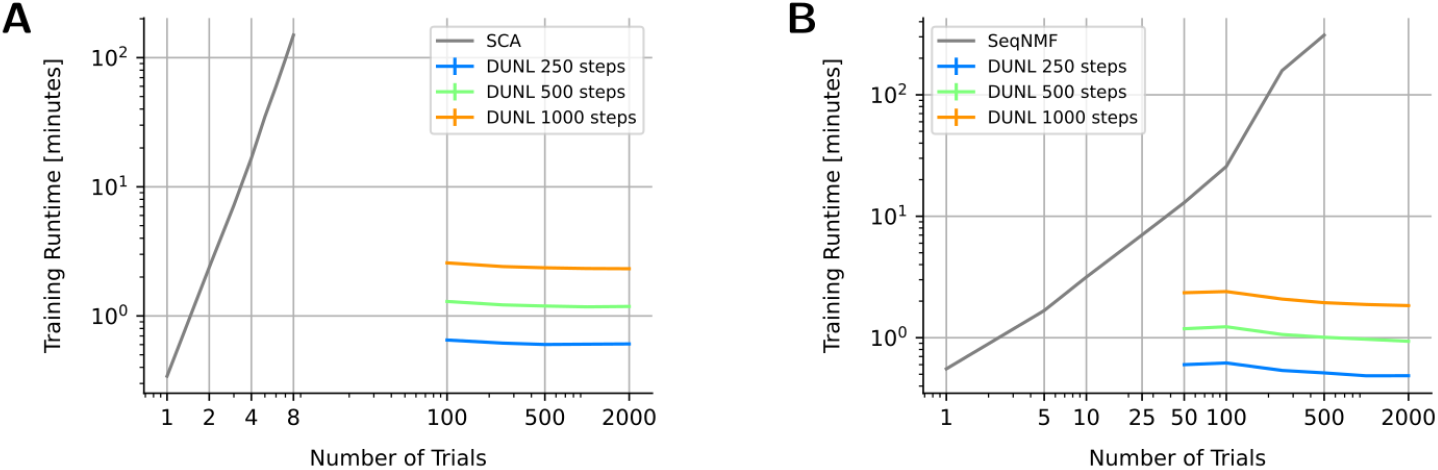
Training runtime. (A) Comparing training runtime of DUNL with SCA as a function of trials used for training. SCA and DUNL are both trained on the same GPU for the data explained in Figure 7. For SCA, trials are stacked to form a matrix. For trials greater than 8, SCA ran out of memory on a 24 GB GPU memory. DUNL’s training is based on batch-based updates; hence, runtimes are reported based on the number of gradient steps (updates). (B) Comparing the training runtime of DUNL with SeqNMF as a function of trials used for training. SeqNMF naturally analyzes a matrix of data of form Neurons × Time; hence, for analysis of multi-trials, trials are stacked to form a matrix. SeqNMF is run on the CPU of Apple M2 Max with 32 GB of memory. Similar to (A), DUNL is trained based on mini-batches; hence, runtime is reported as a function of update steps.

**Figure S20.**
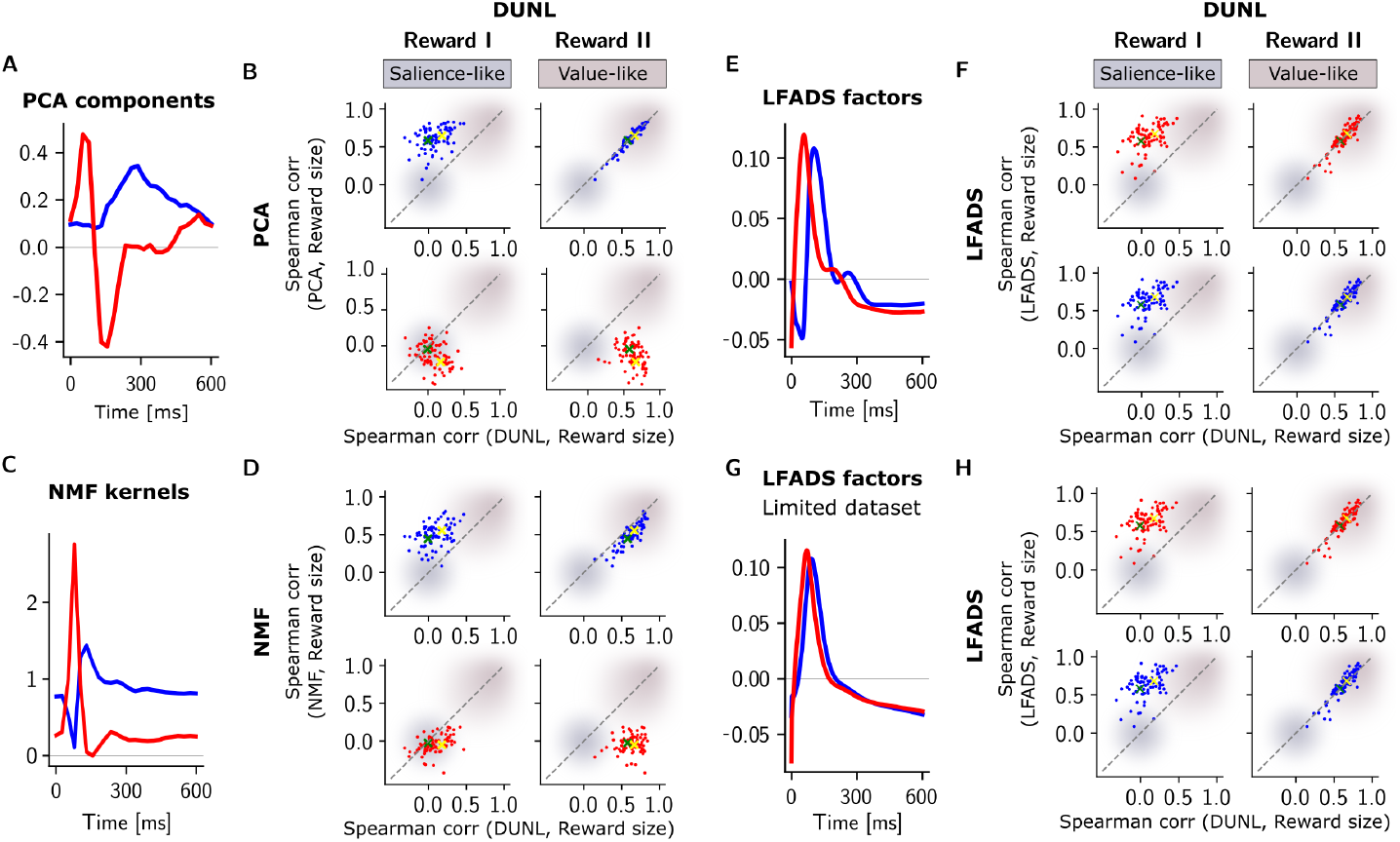
Comparison of DUNL with classical dimensionality reduction and a deep learning framework for the dopamine spiking data. These methods are applied on windowed data of size 600 ms starting from the reward onset from dopamine spiking dataset (Figure 2). PCA is applied to standardized data, NMF is applied to the raw binned data, and LFADS is applied to the raw data. For PCA and NMF, the neuron’s baseline activity at each trial is subtracted before pre-processing. (A) PCA kernels with PC1 in blue and PC2 in red color. (B) Scatter plot of Spearman’s rank correlation of DUNL codes for the Reward I (salience-like) and Reward II (value-like) kernels (left and right, respectively) in the x-axis and of Spearman’s rank correlation of PCs and reward size on the y-axis. Salience-prone region for both methods is shaded in gray-blue (low correlation with reward size) while value-prone region for both methods is shaded in mountbatten pink (larger values for the correlation with reward-size). The blue PC contains value information similar to DUNL’s Reward II but the red PC contains salience and an anti-correlation with value. For a fair comparison, the neurons’ baseline activity at each trial is pre-estimated (instead of inferred) for DUNL. (C) NMF kernels. (D) Scatter plot of Spearman’s rank correlation of DUNL codes for the Reward I and Reward II kernels (left and right, respectively) in the x-axis and of Spearman’s rank correlation of NMF coefficients and reward size on the x-axis. The blue kernel is salience-like, and the red kernel is value-like, but DUNL’s Reward II kernel still outperforms the red NMF kernel at representing value. Similar to PCA, for a fair comparison, the neurons’ baseline activity at each trial is pre-estimated (instead of inferred) for DUNL. (E) LFADS: the average of two factors over the dataset learned by LFADS (the factors are zero-mean and normalized for visualization purposes). (F) Spearman’s rank correlation of DUNL codes and reward size (x-axis) in comparison to Spearman’s rank correlation of the temporal average of the LFADS factors and reward size (y-axis). The comparison of Spearman correlations from DUNL and LFADS shows that both LFADS factors capture similar statistics, only similar to the Reward II kernel in DUNL (value-like); LFADS fails to deconvolve the reward response into salience and value. (G-H) Same as (E-F) for LFADS run on the limited dataset using < 8% of the data (same dataset as Figure S5). Compared to the full dataset scenario, the Spearman’s correlation results hold; however, certain details on the factors such as the local bump around 200 ms is not captured.

**Figure S21.**
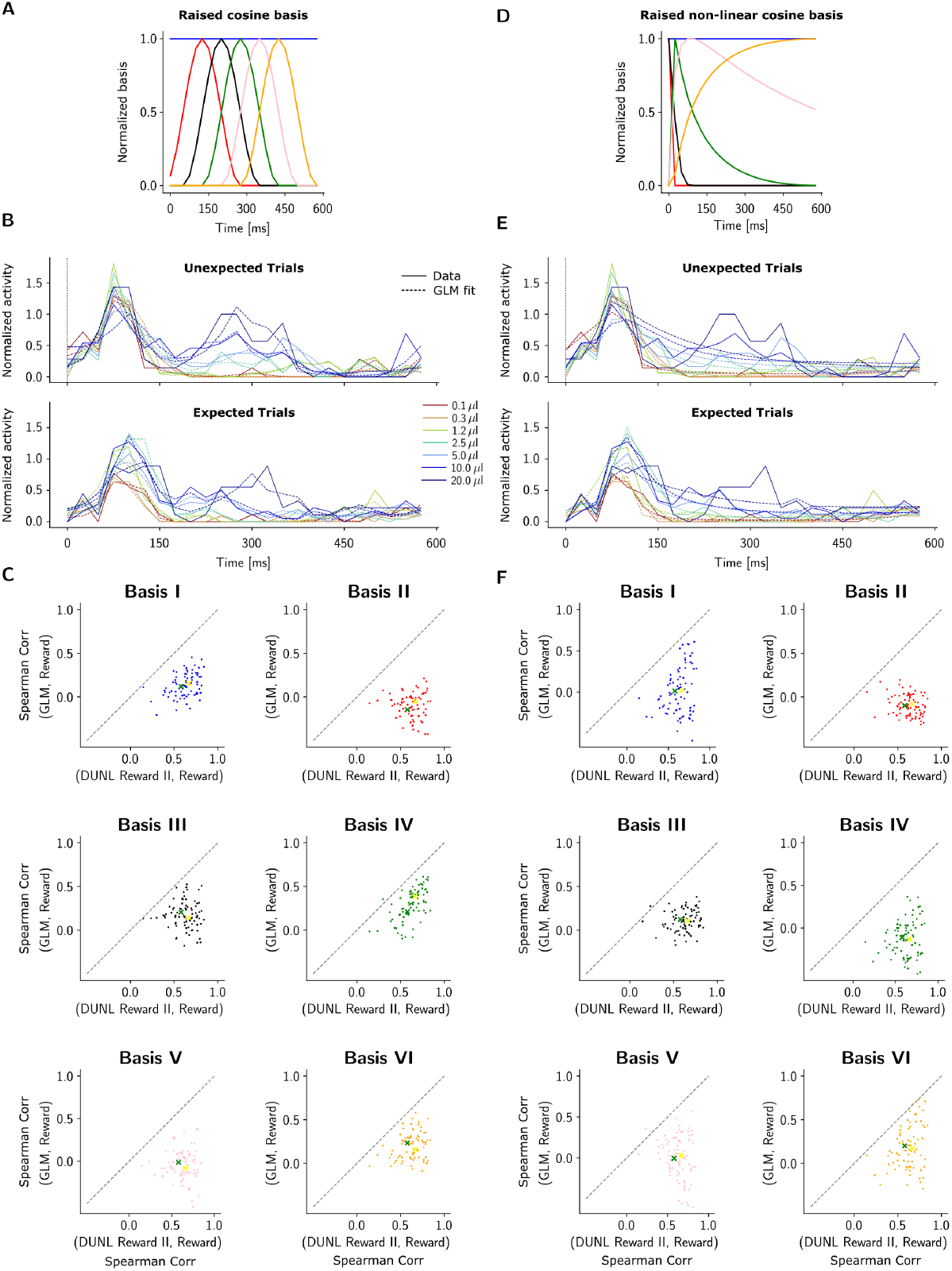
Comparison of DUNL with GLM [34]. We used Poisson GLM regression with a set of pre-defined family of basis functions, for the dopamine spiking data. Similar to the comparison with the dimensionality reduction, the methods are applied on windowed data of size 600 ms starting from the reward onset from dopamine spiking data (Figure 2). The goal of this experiment is to highlight the benefit of learning kernels, as in DUNL, as opposed to designing them as in [34]. (A) Raised cosine bases (normalized bases with 0/1 min/max are shown). The bias is shown as the first base (blue). (B) An example of trial reconstruction by GLM averaged over trial types. (C) The Spearman’s rank correlation of DUNL value code and reward size (x-axis) in comparison to Spearman’s rank correlation of coefficients of each of the GLM basis and reward size (y-axis). The comparison shows that neither of the bases is representative of value response; The predefined bases do not offer interpretability from the point of view of deconvolving the reward response into salience and value. The yellow and green markers show the average across *unexpected* and *expected* trials. (D) Nonlinear raised cosine bases (normalized bases with 0/1 min/max are shown). The first (blue) bases represents the constant bias term. (E) An example of trial reconstruction by GLM averaged over trial types using the nonlinear raised cosines. (F) The Spearman’s rank correlation of DUNL value code and reward size (x-axis) in comparison to Spearman’s rank correlation of coefficients of each of the GLM bases and reward size (y-axis) for the nonlinear raised cosine case. The presence of dots below the diagonal line indicates that the value code offered by DUNL is a better representative of the reward amount. Overall, this emphasizes the lack of interpretability of GLM with pre-defined family of basis functions within the context of deconvolving the single-trial spiking data into interpretable components.

#### Supplementary Discussion: DUNL vs. Prior Works

The proposed method is built on a convolutional dictionary learning generative model that learns localized kernels and infers sparse representations, deconvolving single-trial neuronal activity into local low-rank components. While DUNL shares the convolution operation with methods like convNMF [35, 36], seqNMF[37], and *PP-Seq* [15], it distinguishes itself through scalability, flexibility, representation properties, and the use of the unrolling framework, which leads to the design of a deep neural network. This manuscript introduces the unrolling concept and demonstrates how to integrate domain knowledge (generative models) into neural networks. Below, we describe in detail how DUNL distinguishes itself from prior works.

Incorporating a generative model into deep learning offers three main benefits: for classical methods, it translates generative models into deep neural architectures, enhancing scalability for large datasets with faster runtimes, efficient optimizers like Adam, and flexible trial designs. For deep learning methods, it strengthens the neural network’s inductive bias, improving generalization with limited data and adding domain knowledge for better interpretability. For hybrid approaches, the unrolling technique adds architectural flexibility, allowing models to learn more from data, which classical methods lack, potentially boosting performance [68].

Below, we summarize methodological differences between DUNL and previously published tools for the analysis of neural data, with a focus on convolutional methods and single-trial analysis methods:

- **DUNL vs. convNMF/seqNMF [35–37]**.
  - ConvNMF and seqNMF, like DUNL, use convolution but are designed to detect temporal patterns across neuron populations, analyzing data in a matrix format (*Neurons* × *Trial length*). SeqNMF focuses on learning population-wide kernels with one shared code, whereas DUNL can learn localized kernels per neuron or across all neurons, offering individual codes per neuron and trial. DUNL is optimized for detecting events that trigger impulse-like responses, using *ℓ*_1_ norm minimization and k-sparse coding for high sparsity, a feature missing in seqNMF. Our experiments highlight DUNL’s superiority over seqNMF in inferring highly sparse event representations.
  - ConvNMF/seqNMF are designed to analyze a data matrix of neural population activity over time (Neurons × Trial length), requiring trials to be stacked along the time axis for multi-trial data. In contrast, DUNL can handle a tensor (*Trials* × *Neurons* × *Time*), allowing it to learn kernels across both neurons and trials simultaneously. This batch-based, gradient-based approach enables DUNL to scale efficiently, handling trials of different lengths and large datasets. SeqNMF, on the other hand, is limited by memory constraints and struggles with scaling to a large number of trials. Our new experiments demonstrate DUNL’s scalability and compare its runtime with seqNMF.
  - DUNL models spikes as binary events, dividing spiking neural activity into small time bins (5 ms to 25 ms at 1 ms resolution) and using a binomial distribution to model firing rates. For calcium imaging data, DUNL adapts by using a Gaussian model to respect the continuous nature of the signal. In contrast, SeqNMF assumes continuous data and employs least square minimization for non-negative matrix factorization, requiring data smoothing or trial averaging. SeqNMF also searches for non-negative factors, which limits its ability to capture negative values, such as the value-like factor observed in the dopamine experiments.
  - In DUNL terms, SeqNMF decomposes the neural activity matrix into a code and dictionary matrix, where the code amplitude is shared across the neuron population, and amplitude differences between neurons are encoded in the dictionary (W). This assumes that the relative response amplitude across neurons is fixed, which may not hold in certain cases. In contrast, DUNL provides a code amplitude for each event per neuron, allowing it to capture a broader range of response strengths. Lastly, DUNL can encourage neurons to fire together and share kernels, a feature SeqNMF lacks explicitly (though, in theory, it can recover patterns where multiple neurons are active simultaneously).
- **DUNL vs. PP-seq [15]**. From a modeling perspective, DUNL’s generative model differs from PP-seq in key ways. PP-seq focuses on identifying sparse sequences of patterns across the neural population, similar to convNMF/seqNMF, while providing uncertainty estimates. Its goal is to explain neural activity by assuming impulse responses follow a Gaussian form [15], parameterized by amplitude, mean, and variance. In contrast, DUNL aims to not only fit the data but also characterize neuron activity in response to localized events of various types. DUNL learns non-parametric kernels, enabling it to capture a broader range of impulse responses beyond Gaussian forms. Additionally, DUNL allows flexibility in its latent representation by modifying the activation and loss functions during training. For example, in Figure 4 (whisker data), we applied group sparsity regularization to encourage neurons to fire together, a feature missing in PP-seq.
- **DUNL vs. SCA [33]**. Sparse component analysis (SCA) is a newly released method that decomposes data into sparse components, and there are key differences between SCA and DUNL. First, SCA enforces orthogonal factors, whereas DUNL does not. DUNL’s unrolling architecture allows it to learn correlated kernels, requiring more iterations to deconvolve them, but it can do so effectively. Second, SCA is not a convolutional method—its factors cover the full trial duration, limiting it to finding low-dimensional structures only in time-aligned or windowed data. In contrast, DUNL is convolutional and can detect low-dimensional structures that shift across trials. Third, SCA only handles trials of the same length, whereas DUNL can process trials of varying lengths. Fourth, SCA works with a data matrix rather than a tensor, forcing trials to be stacked in the time dimension. Lastly, SCA is not scalable, as shown in this manuscript.
- **DUNL vs. GPFA [28]**: Gaussian-Process Factor Analysis (GPFA) was developed for spiking data, and it may not easily apply to other recording modalities, as calcium or neuromodulator sensor activities might not benefit from the square-root transform. GPFA focuses on independent factors in population activity, discarding single-neuron variability as *independent variability*. In contrast, DUNL provides single-neuron, single-event codes, useful for neurons that multiplex activity from different events in a single trial. Additionally, GPFA does not scale well—fitting a smoothing kernel for each neuron becomes impractical in large datasets. A related method, variational latent Gaussian Process (vLGP) [29], is faster and provides good predictability at fine timescales for spiking data.
- **DUNL vs. mTDR [30]**: Model-based targeted dimensionality reduction (mTDR) identifies neural population coding of task variables by linearly mapping stimuli to neural activity, with weights specific to each neuron. In contrast, DUNL estimates a code per neuron and per trial, even when kernels are shared. mTDR requires the support (task variables) to be provided, similar to GLMs, while DUNL can infer the support or accept it as input. Additionally, DUNL can handle tasks with variable trial lengths and does not rely on a global linear mapping between task variables and neural responses.

